# mRNA structural dynamics shape Argonaute-target interactions

**DOI:** 10.1101/822452

**Authors:** Suzan Ruijtenberg, Stijn Sonneveld, Tao Ju Cui, Ive Logister, Dion de Steenwinkel, Yao Xiao, Ian J. MacRae, Chirlmin Joo, Marvin E. Tanenbaum

## Abstract

Small RNAs (such as miRNAs, siRNAs and piRNAs) regulate protein expression in a wide variety of biological processes and play an important role in cellular function, development and disease. Association of small RNAs with Argonaute (AGO) family proteins guide AGO to target RNAs, generally resulting in target silencing through transcriptional silencing, translational repression or mRNA degradation. Here we develop a live-cell single-molecule imaging assay to simultaneously visualize translation of individual mRNA molecules and their silencing by human AGO2-siRNA complexes. We find that siRNA target sites are commonly masked *in vivo* by RNA secondary structures, which inhibit AGO2-target interactions. Translating ribosomes unmask AGO2 binding sites, stimulating AGO2-target interactions and promoting mRNA degradation. Using a combination of mathematical modeling and experiments, we find that mRNA structures are highly heterogeneous and continuously refolding. We show that structural dynamics of mRNAs shape AGO2-target recognition, which may be a common feature controlling mRNA-protein interactions.

## Introduction

Precise regulation of gene expression is essential for cell fate and function. A family of small RNAs of 20-32 nucleotides (nt), including microRNAs (miRNAs), small interfering RNAs (siRNAs), and Piwi interacting RNAs (piRNAs), is an important class of molecules that can regulate RNA and protein levels in cells (Bartel, 2018; Gebert and MacRae, 2019; Ghildiyal and Zamore, 2009; Malone and Hannon, 2009; Ozata et al., 2019). Small RNAs associate with a member of the Argonaute (AGO) family of proteins, and together they form the RNA-induced silencing complex (RISC) (Liu et al., 2004; Meister, 2013; Schirle et al., 2014; Song et al., 2004). Association with a small RNA guides AGO to a target RNA via Watson-Crick base pairing between the small RNA and the target RNA. Upon binding to the target RNA, AGO either cleaves the target RNA (also referred to as slicing) or recruits additional effector proteins to induce other types of target repression, including heterochromatin formation, mRNA deadenylation and decay, and translational repression (Iwasaki et al., 2015; Jonas and Izaurralde, 2015). The precise mechanism of repression that occurs upon AGO recruitment to the RNA target likely depends on the AGO family member, cellular context, and the pattern of sequence complementarity between the small RNA and its target (Bartel, 2009; Filipowicz et al., 2008; Ghildiyal and Zamore, 2009; Jonas and Izaurralde, 2015).

Mechanisms of AGO-target interactions have been extensively studied using *in vitro* single-molecule imaging, as well as biochemical and structural approaches (Bartel, 2018; Chandradoss et al., 2015; Jo et al., 2015; Salomon et al., 2015; Schirle et al., 2014; Wee et al., 2012; Yao et al., 2015). These studies have shown that initial AGO-target interactions are established by nt 2-4 of the small RNA, and are subsequently extended to nt 2-8 (referred to as the seed region), which further stabilizes the interaction between AGO and the target site (Chandradoss et al., 2015; Jo et al., 2015; Salomon et al., 2015; Wee et al., 2012). Further base pairing beyond the seed region is accompanied by conformational changes in AGO, and is required for target cleavage by AGO2, the major human AGO family member with endonucleolytic cleavage activity in somatic cells (Jung et al., 2013; Liu et al., 2004; Meister et al., 2004; Schirle et al., 2014; Sheu-Gruttadauria and MacRae, 2017). However, AGO-target interactions *in vivo* are likely more complex. First, the cytoplasm contains many different RNA species, providing a far more challenging venue for AGO to correctly identify the target site. Second, hundreds of RNA binding proteins (RBPs) exist *in vivo*, which may compete with AGO for target site binding (Hentze et al., 2018; Jankowsky and Harris, 2015; Strobel et al., 2018). Third, *in vivo*, RNA targets are often translated by ribosomes, which may displace AGO proteins from the target RNA during translation through physical collisions between ribosomes and AGO proteins. Indeed, translating ribosomes have been suggested to hinder miRNA-dependent silencing of mRNAs, possibly explaining why miRNA binding sites are generally enriched in the 3’ UTRs of mRNAs (Bartel, 2018; Grimson et al., 2007; Gu et al., 2009). Finally, *in vivo* RNA targets are typically at least an order of magnitude longer than the RNA oligonucleotides that are frequently used as targets for *in vitro* studies. These long RNA targets have a far greater potential to adopt secondary and tertiary structures, which can affect association of RBPs, as well as AGO target site availability (Ameres et al., 2007; Beaudoin et al., 2018; Becker et al., 2019).

RNA structures are highly pervasive in the mammalian transcriptome (Aw et al., 2016; Bevilacqua et al., 2016; Ding et al., 2014; Gong et al., 2018; Lu et al., 2016; Mustoe et al., 2018; Rouskin et al., 2014; Strobel et al., 2018; Wan et al., 2014; Zubradt et al., 2017). These mRNA structures are often conserved and can be functionally important for RNA regulation, either by promoting or inhibiting interactions between RBPs and their RNA target (Al-Hussaini et al., 2008; Dominguez et al., 2018; Hentze et al., 2018; Liu et al., 2015). For example, RNA targets in which AGO binding sites are embedded in highly structured regions poorly interact with AGO both *in vitro* and *in vivo* (Ameres et al., 2007; Beaudoin et al., 2018; Becker et al., 2019). Recent studies have also revealed that mRNAs can undergo structural rearrangements throughout an mRNA’s life. For example, when zebrafish embryos undergo the maternal-to-zygotic transition (Beaudoin et al., 2018), or as mRNAs transition from the nucleus to the cytoplasm (Adivarahan et al., 2018; Sun et al., 2019), or upon translation of mRNAs by ribosomes (Beaudoin et al., 2018; Mizrahi et al., 2018; Mustoe et al., 2018). While these studies demonstrate that mRNAs can adopt distinct structural conformations, they provide limited insights into the dynamics of mRNA structure, especially on the short timescales at which AGO-target interactions likely occur. Dynamics of mRNA folding, un-folding and re-folding on short timescales may be critical for AGO-target interactions, as well as of many other RBPs, and may thus be important for mRNA post-transcriptional regulation.

To study the dynamics of AGO-target interactions *in vivo* in the context of translated target mRNAs, we have developed an imaging approach to simultaneously visualize mRNA translation and AGO-mediated repression of individual mRNA molecules in live cells. As a model system, we have used AGO2 in complex with an siRNA in human somatic cells. We find that many AGO2 target sites are masked to varying extents by secondary structures in the target mRNA, which inhibit AGO2-target interactions. mRNAs adopt distinct structural conformations, and are continuously refolding on the seconds-to-minutes timescale. Moreover, we find that translating ribosomes unfold mRNA structure, thereby unmasking AGO2 binding sites and promoting AGO2-target interactions and mRNA cleavage. Taken together, our study provides a new method to study AGO target repression *in vivo* and reveals that mRNA structural dynamics shape AGO2-target interaction dynamics, which may be relevant for many RBP-mRNA interactions.

## Results

### An assay to study AGO2-dependent mRNA target silencing by single-molecule live-cell imaging

Recently, an imaging method was developed to visualize cleavage of single mRNAs (Horvathova et al., 2017). However, this method does not allow simultaneous observation of mRNA translation, so the effects of AGO on translation efficiency, and the effects of translating ribosomes on AGO-dependent mRNA silencing cannot be studied using this approach. To study AGO2 activity in the context of a translated target mRNA we adapted a live-cell imaging method that we and others recently developed to visualize translation of individual mRNA molecules (Figure 1A, left panel) (Morisaki et al., 2016; Pichon et al., 2016; Wang et al., 2016; Wu et al., 2016; Yan et al., 2016). Our translation imaging approach uses the ‘SunTag’ system (Tanenbaum et al., 2014), which consists of a genetically-encoded single-chain variable fragment antibody fused to GFP (scFv-GFP), and a reporter mRNA encoding 24 copies of the SunTag antibody peptide epitope upstream of a gene of interest (for example the kinesin KIF18B) (Figures 1A, left panel, and 1B). Translation of the reporter mRNA leads to rapid co-translational binding of the scFv-GFP to the SunTag peptide epitopes, resulting in fluorescence labeling of the nascent polypeptide (Figure 1A, left panel), thereby allowing visualization of reporter mRNA translation. As multiple ribosomes generally translate an mRNA molecule simultaneously, a very bright fluorescence signal is associated with each translating mRNA molecule. mRNAs are also labeled independently of translation by introducing 24 copies of the PP7 coat protein (PCP) binding site (PBS) into the 3’ UTR of the reporter mRNA and by co-expression of the PCP fused to two copies of mCherry (Chao et al., 2008; Ruijtenberg et al., 2018; Yan et al., 2016). To facilitate imaging and long-term tracking of single mRNAs, we developed a system to tether mRNAs to the plasma membrane by fusing a C-terminal membrane anchor (a CAAX motif) to the PCP-2xmCherry (Figure 1A, left panel), which does not detectably affect translation dynamics (Hoek et al., 2019; Yan et al., 2016). Finally, the translation reporter is placed under control of a doxycycline-inducible promoter to allow temporal control of reporter mRNA expression.

**Figure 1.**
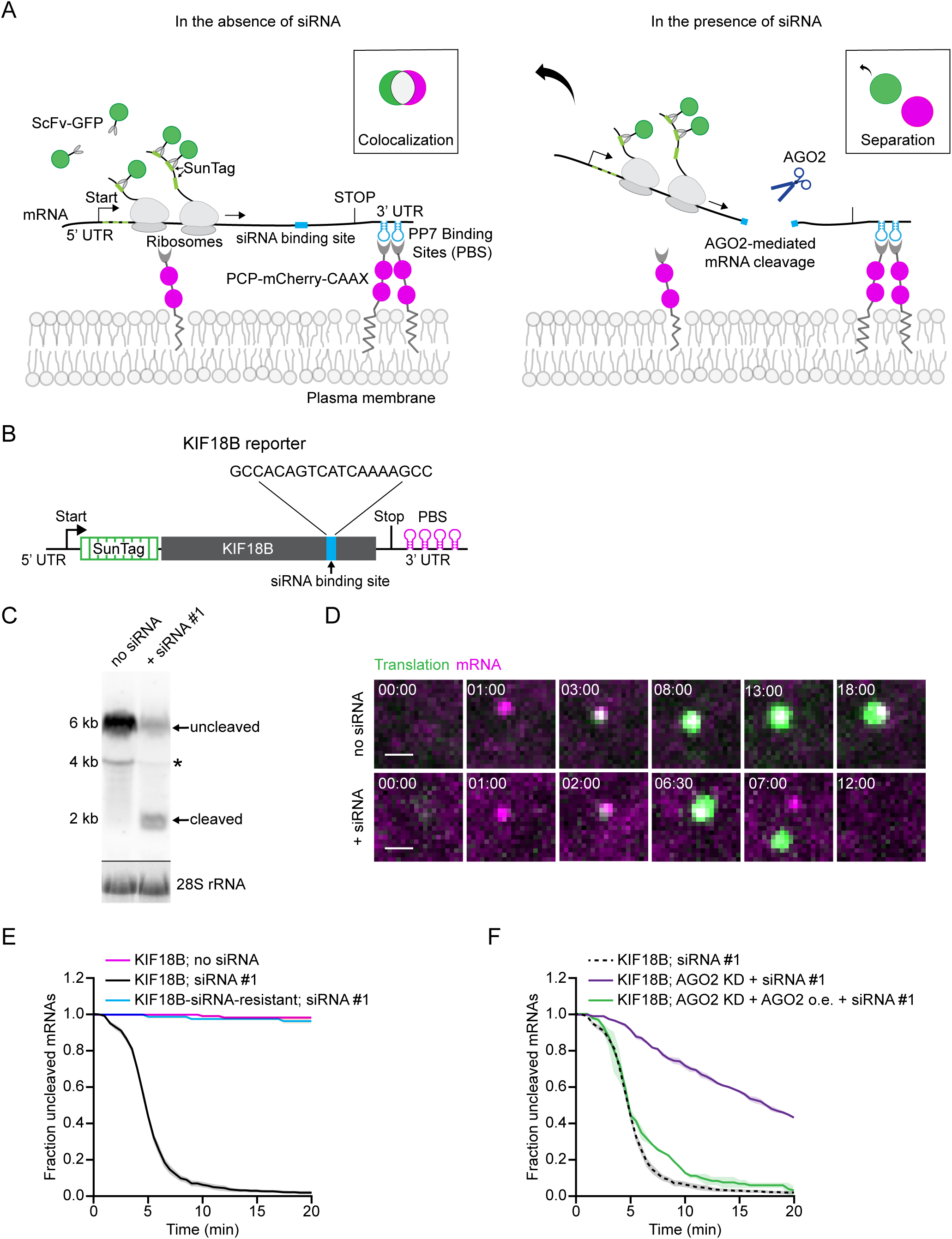
Observing AGO2-dependent mRNA target silencing by single-molecule live-cell imaging. (A) Schematic of the single-molecule imaging assay used to visualize AGO2-mediated mRNA silencing in the absence (left) or presence (right) of siRNA. Green and magenta spots (insets) show nascent polypeptides (translation) and mRNA, respectively, as observed by microscopy. (B) Schematic of the mRNA reporter. (C) Northern blot of cells expressing the reporter mRNA shown in (B), either without siRNA or transfected with KIF18B siRNA #1. (top) Upper band (uncleaved) represents full-length reporter mRNA, lower band (cleaved) represents the 3’ cleavage fragment. Asterisk indicates an additional 4 kB band that may represent a shorter isoform of the reporter mRNA. (bottom) 28S rRNA acts as a loading control. (D) Representative images of mRNA molecules of the reporter shown in (B) expressed in SunTag/PP7 cells without (top) or with siRNA (bottom). Scale bar, 1 µm. Time is shown in min:sec. (E-F) SunTag/PP7 cells expressing indicated reporters were transfected with 10 nM KIF18B siRNA #1, where indicated. The time from first detection of translation until separation of GFP and mCherry foci (i.e. mRNA cleavage) is shown. Solid lines and corresponding shaded regions represent mean ± SEM. (F) Cells expressing dCas9-KRAB were infected with sgRNA targeting endogenous AGO2 (AGO2 KD), or with full length AGO2 (AGO2 o.e. (overexpression)), where indicated. Dotted lines indicate that the data is replotted from an earlier figure panel for comparison. Number of measurements for each experiment is listed in Table S1. See also Figure S1 and Videos S1, and S2.

To study AGO2-dependent silencing of individual mRNA molecules in live cells, we designed an siRNA with full complementarity to a site in the coding sequence of our reporter gene (KIF18B) (Figure 1B). Analysis by northern blot, qPCR, and single-molecule FISH (smFISH) of cells transfected with the siRNA revealed a strong reduction in reporter mRNA levels, and the formation of 3’ and 5’ cleavage fragments (Figures 1C, and S1A-F), indicating that the reporter mRNA was efficiently targeted for endonucleolytic cleavage by the siRNA. Next, we wanted to visualize siRNA-mediated mRNA cleavage of single mRNAs in living cells by time-lapse microscopy. In our reporter, the siRNA target site is located in the coding sequence of KIF18B, 3891 nt downstream of the start codon and 470 nt upstream of the stop codon. Endonucleolytic cleavage at the siRNA binding site will result in the formation of a 5’ cleavage fragment, which contains ∼90% of the KIF18B coding sequence and is therefore expected to be associated with the large majority of ribosomes (and associated GFP signal). The 3’ cleavage fragment is associated with a small part of the KIF18B coding sequence (and thus a small fraction of the ribosomes) and also with all the PCP-mCherry fluorescence (Figures 1A, right panel). Since mRNAs are tethered to the plasma membrane through the PCP system in the 3’ UTR, the mCherry-positive 3’ mRNA cleavage fragment is expected to remain in the field-of-view upon cleavage (until it is degraded by an RNA exonuclease), while the GFP positive 5’ fragment is expected to diffuse away out of the field-of-view (Figure 1A, right panel). Thus, we reasoned that mRNA cleavage would result in a separation of GFP and mCherry foci in microscopy-based images.

Human U2OS cells stably expressing scFv-GFP, PCP-2xmCherry-CAAX (referred to as SunTag/PP7 cells), and the reporter mRNA (Figure 1B) were treated with doxycycline for 15-30 min to induce expression of the reporter and imaged for ∼1 hr using a spinning disk confocal microscope at a time interval of 30 s. While few mRNAs were present in the field-of-view before transcription induction by doxycycline addition, many newly transcribed mRNAs (∼10-50) appeared in the cytoplasm during imaging. We focused on the newly-transcribed mRNAs that appeared in the field-of-view before the first round of translation had initiated and followed these mRNAs over time. In the absence of siRNA, translation of newly-transcribed reporter mRNAs initiated rapidly (2.4 ± 3.1 min, mean ± SD) after mRNA appearance in the field-of-view (as indicated by the appearance of a GFP signal), and was generally maintained at high levels for the duration of the movie (∼1 hr) (Figure 1D, upper panel; Video S1).

In siRNA-transfected cells, translation initiated with similar kinetics as in control cells (2.1 ± 2.1 min, mean ± SD). However, in contrast to control cells, GFP and mCherry foci frequently separated within minutes of translation initiation (92% of mRNAs in 10 min) (Figures 1D, lower panel, and 1E; Video S2). Separation of GFP and mCherry foci was due to AGO2-dependent endonucleolytic cleavage, rather than translation termination of individual ribosomes (which could in principle also result in separation of GFP and mCherry foci), as foci separation was largely eliminated by mutation of the siRNA binding site in the mRNA reporter or by depletion of AGO2 (Figures 1E, 1F, and S1G). Furthermore, the fluorescence intensity of the large majority (97%) of GFP foci after GFP-mCherry foci separation was higher than the intensity of a single SunTag polypeptide (Figure S1H; see methods), consistent with endonucleolytic cleavage, but not with translation termination of individual ribosomes. These results show that GFP and mCherry foci separation reflects endonucleolytic cleavage of mRNAs by AGO2 and show that GFP-mCherry foci separation can be used as a tool to study AGO2 activity on single mRNA molecules in real-time in living cells.

To determine whether mechanisms of target repression other than mRNA cleavage occurred upon siRNA transfection, we performed two additional sets of experiments. First, we found that transcription of the reporter mRNA was unaffected by the AGO2/siRNA complex, as determined by smFISH of nascent RNAs (Figure S1I). Second, we examined the translation efficiency of reporter mRNAs in the presence or absence of siRNA, but did not observe a significant effect on translation rates (Figures S1J and S1K). Together, these results show that endonucleolytic cleavage is the predominant mechanism of action of AGO2/siRNA complexes on their target mRNAs.

### Ribosomes stimulate AGO2-dependent mRNA cleavage

Upon closer examination of the kinetics of mRNA cleavage, we noticed that the majority of mRNAs (56%) were cleaved in a short time window between 4-6 minutes after the start of translation (i.e. after first appearance of GFP signal on an mRNA) (see Figure 1E). Intriguingly, this time window closely coincides with the time at which the first ribosome is expected to arrive at the AGO2 cleavage site based on measurements of ribosome translocation speed (∼3 codons/s) (Yan et al., 2016) (see also Figure S2A, pink bars). To test whether arrival of the first ribosome at the AGO2 binding site stimulates mRNA cleavage, we introduced a spacer sequence of 1443 nt between the SunTag and AGO2 binding site (Figure 2A, left), which should delay the arrival of the ribosome at the cleavage site by ∼2.5 min. Analysis of the timing of mRNA cleavage confirmed a delay in cleavage of ∼2.5 min (Figure 2B), suggesting that ribosomes arriving at the AGO2 binding site indeed stimulate AGO2-dependent mRNA cleavage. Furthermore, treatment of cells with the ribosome translocation inhibitors cycloheximide (CHX) or Emetine (Eme), or introduction of a stop codon 1050 nt upstream of the AGO2 binding site all strongly inhibited AGO2-dependent mRNA cleavage (Figures 2A, right, 2C, and S2B). Finally, we found that the cleavage rate over the first ∼3 min from GFP appearance was similar for all reporters (Figure 2C), consistent with the fact that ribosomes have not arrived at the cleavage site yet during this time period. Together, these results show that arrival of translocating ribosomes at the siRNA binding site stimulates mRNA cleavage by AGO2.

**Figure 2.**
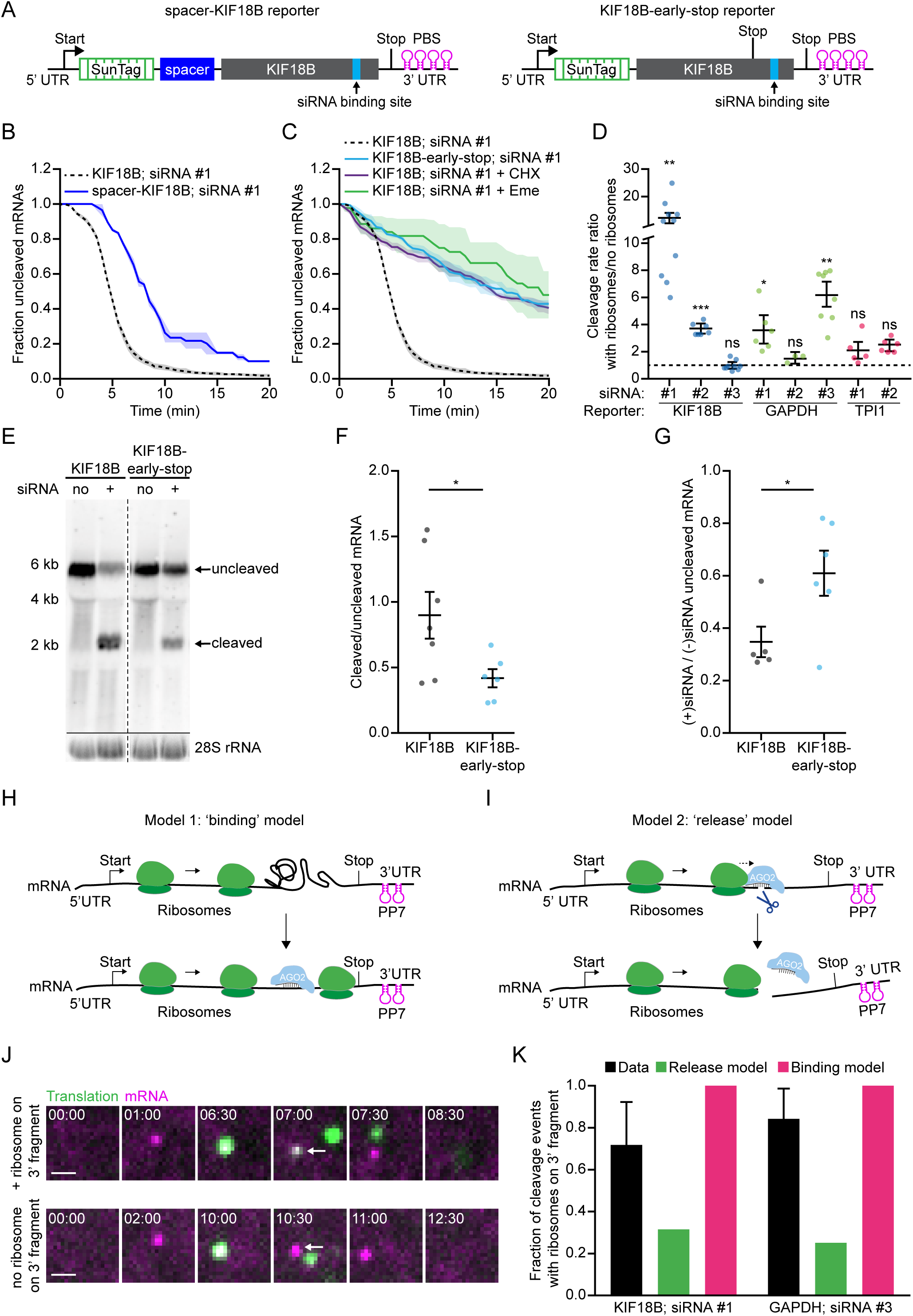
Ribosomes stimulate AGO2-dependent mRNA cleavage by promoting AGO2-target interactions. (A) Schematic of indicated reporters. (B-C) SunTag/PP7 cells expressing indicated reporters were transfected with 10 nM KIF18B siRNA #1 and treated with CHX or Emetine, where indicated. The time from first detection of translation (B, C) or from CHX/Eme addition (C; + Eme, + CHX), until separation of GFP and mCherry foci (i.e. mRNA cleavage) is shown. Solid lines and corresponding shaded regions represent mean ± SEM. Dotted lines indicate that the data is replotted from an earlier figure panel for comparison. (D) Ratio of the cleavage rates in the presence and absence of translating ribosomes is shown for the indicated siRNAs and reporters (see methods). (E) Northern blot of cells expressing the KIF18B or KIF18B-early-stop reporter, either non-transfected (no siRNA) or transfected with 10nM KIF18B siRNA #1 (+ siRNA). (top) Upper band (uncleaved) represents the full-length reporter mRNA, the lower band (cleaved) represents the 3’ cleavage fragment. (bottom) 28S rRNA acts as a loading control. (F) Ratio of the northern blot band intensity for bands representing cleaved and uncleaved mRNAs for the + siRNA condition. (G) Ratio of the intensity of the + siRNA and - siRNA uncleaved bands. (F-G) Each dot represents a single experiment and lines with error bars indicate the mean ± SEM. (H-I) Schematics for (H) ‘binding’ and (I) ‘release’ models explaining how ribosomes could stimulate AGO2-mediated mRNA cleavage. (J) Representative images of mRNA molecules in SunTag/PP7 cells expressing the KIF18B-ext reporter showing cleavage events with (top) or without (bottom) a ribosome on the 3’ cleavage fragment. Arrows indicate 3’ cleavage fragments. Time is indicated as min:sec. (K) The fraction of mRNAs that contains a ribosome on the 3’ cleavage fragment is shown for the data (black bars) and for the indicated models (green and pink bars). P-values in (D, F-G) are based on a two-tailed Student’s t-test. P-values are indicated as * (p < 0.05), ** (p < 0.01), *** (p < 0.001), ns = not significant. Number of measurements for each experiment is listed in Table S1. See also Figure S2.

To determine whether ribosome-stimulated cleavage by AGO2 was observed for other siRNA and mRNA sequences as well, we designed new reporters where the KIF18B sequence was replaced by the GAPDH or TPI1 coding sequence, and we designed 3 siRNAs targeting GAPDH, 2 siRNAs targeting TPI1, as well as two additional siRNAs targeting KIF18B. Overall, 4 of 8 tested siRNAs showed a significantly (p < 0.05, two-tailed t-test) higher cleavage rate by AGO2 in the presence compared to the absence of translating ribosomes (Figures 2D and S2C-I). Ribosome-stimulated cleavage by AGO2 was further confirmed by northern blot analysis, which revealed a significant reduction in mRNA cleavage upon introduction of a stop codon upstream of the AGO2 binding site (Figures 2E-G, and S2J). Together, these results demonstrate that ribosome-stimulated cleavage by AGO2 is a commonplace phenomenon in living cells.

### Ribosomes promote AGO2-target interactions

We considered two models explaining how ribosomes could stimulate AGO2-dependent mRNA cleavage. First, it is possible that ribosomes stimulate cleavage by promoting AGO2-mRNA target interactions (‘binding’ model; Figure 2H). For example, ribosomes may clear the AGO2 binding site of RBPs or unfold RNA structures that mask the AGO2 binding site. Second, it is possible that ribosome collisions with AGO2 stimulate release of the 5’ and 3’ cleavage fragments from AGO2 after endonucleolytic cleavage has occurred (‘release’ model; Figure 2I) (note that our imaging approach cannot distinguish between mRNA slicing and fragment release).

A key prediction of the ‘release’ model is that the ribosome that stimulates cleavage will be located upstream of AGO2 at the moment of cleavage, and will thus remain on the 5’ cleavage fragment. In contrast, in the ‘binding’ model the ribosome that stimulates cleavage will frequently pass over the AGO2 binding site to unmask it and will end up on the 3’ cleavage fragment. Therefore, to distinguish between the ‘release’ and ‘binding’ models, we examined the location of the first ribosome after mRNA cleavage (Figure 2J). For these experiments, we made use of two reporter-siRNA combinations for which cleavage rates were strongly stimulated by ribosomes, KIF18B siRNA #1 and GAPDH siRNA #3 (see Figure 2D). To increase the sensitivity of detection of single ribosomes on the 3’ cleavage fragment, we modified image acquisition conditions (see methods) and extended the coding sequence of the reporters downstream of the cleavage site to increase the time that a ribosome would remain on the 3’ cleavage fragment (Figure S2K). After normalization of the data (see methods) we found that one or more ribosomes were present on the 3’ cleavage fragment in both the KIF18B-ext and the GAPDH-ext reporter for the majority of ribosome-stimulated cleavage events (76% and 85%, respectively) (Figure 2K, black bars). To determine whether these results are more consistent with the ‘binding’ or ‘release’ model, we calculated the expected fraction of cleaved mRNAs with ribosomes on the 3’ cleavage fragment for both models (Figure 2K, green and pink bars; see methods), which revealed that the data was most consistent with the ‘binding’ model. The fraction of cleavage events with a ribosome on the 3’ cleavage fragment was somewhat lower than predicted by the ‘binding’ model though (Figure 2K, compare black and pink bars), which is likely due to imperfect detection of such ribosomes (Figure S2L).

### In vivo kinetics of the AGO2 cleavage cycle

The kinetics of the AGO2 cleavage cycle (i.e. target binding, AGO2 structural rearrangements, mRNA slicing and cleavage fragment release) likely affects the interplay between AGO2 and translating ribosomes. For example, if the slicing activity of AGO2 is slow, ribosomes will frequently collide with AGO2 before slicing has occurred, potentially inhibiting target slicing. While several studies have determined the kinetics of each step of the cleavage cycle of AGO2 *in vitro* (Ameres et al., 2007; Chandradoss et al., 2015; Jo et al., 2015; Lam et al., 2015; Salomon et al., 2015; Wee et al., 2012; Yao et al., 2015), very little is known about the cleavage kinetics *in vivo*. Our system provides an opportunity to determine the cleavage cycle dynamics *in vivo*, from the first step (target binding) until the last step (cleavage fragment release). While the end of the cleavage cycle (fragment release) can be directly established from the fluorescence images, the start of the cleavage cycle (target binding) is not observed directly. However, the start of the cleavage cycle can be inferred; the target binding step can initiate when the target site becomes unmasked, which occurs when the first ribosome arrives at the AGO2 binding site, and the moment of arrival of the first ribosome can be calculated from the data (See Figures 2C and S2A).

To estimate the duration of the entire cleavage cycle *in vivo* (from binding site availability to fragment release), we computed cleavage curves using different theoretical AGO2 cleavage cycle durations (ranging from 1 s to 30 min) (Figure 3A, colored lines show a few example curves; see methods) and compared the computed cleavage curves with the experimentally-derived cleavage curve (Figure 3A). This analysis revealed that a cleavage cycle duration of ≤1 min best fit the data (Figures 3A and 3B, black line). Therefore, we conclude that the entire cleavage cycle, including AGO2/siRNA target binding, mRNA slicing and fragment release occurs within ∼1 min of target site unmasking. Thus, each individual step of the cleavage cycle takes less than 1 min.

**Figure 3.**
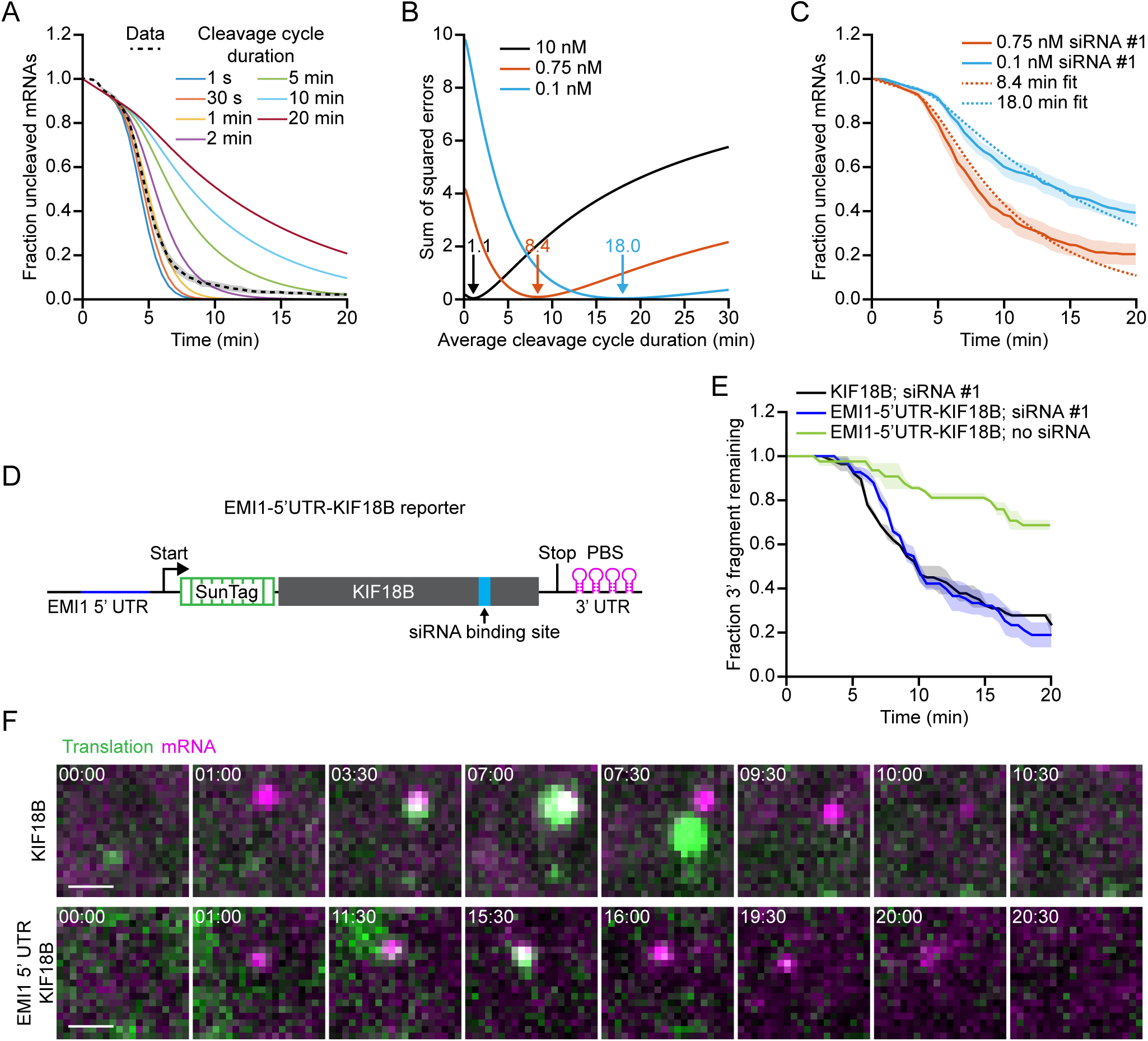
*In vivo* kinetics of the AGO2 cleavage cycle. (A) Simulated cleavage curves for indicated (theoretical) durations of the cleavage cycle (solid colored lines) are compared to the data (KIF18B + 10 nM siRNA #1; black dotted line). Time represents time since GFP appearance. (B) Sum of squared error values are shown for different average cleavage cycle durations for indicated siRNA concentrations. Arrows and values indicate average cleavage cycle duration of the optimal fit. (C) Dotted lines indicate the best cleavage curve fit for the indicated siRNA concentrations. (C, E, F) SunTag/PP7 cells expressing the KIF18B reporter or (E, F) the EMI1-5’UTR-KIF18B reporter were transfected with KIF18B siRNA #1 at indicated concentrations (10 nM in E and F). (C, E) The time from first detection of translation until (C) separation of GFP and mCherry foci (i.e. mRNA cleavage) or (E) mCherry disappearance (i.e. exonucleolytic decay of the 3’ cleavage fragment) is shown. Solid lines and corresponding shaded regions represent mean ± SEM. (D) Schematic of the EMI1-5’UTR-KIF18B reporter. (F) Representative images of a time-lapse movie are shown. Scale bar, 1 µm. Time is shown in min:sec. Note that fluorescent intensities for the KIF18B reporter and the EMI1-5’UTR-KIF18B reporter images are scaled differently to allow visualization of the very dim GFP signal associated with translation by a single ribosome (bottom). Number of measurements for each experiment is listed in Table S1.

Next, we focused on the binding step in more detail. We decreased the concentration of siRNA from 10 nM to 0.75 nM or 0.1 nM to slow down the binding step (Figure 3C, solid lines). Comparison of the cleavage curves for 0.75 nM and 0.1 nM siRNA with simulated cleavage time distributions, revealed a good fit with an average cleavage cycle duration of ∼8 min and ∼18 min, respectively (Figures 3B, orange and blue line, and 3C, dotted lines). Since the catalysis and release steps are unlikely affected by a decrease in the siRNA concentration, these results suggest that even at moderately high siRNA concentrations (i.e. at least between 0.75 nM and 10 nM siRNA), target binding is the rate-limiting step, whereas AGO2 structural rearrangements, catalysis and fragment release all occur relatively fast (<1 min).

The experiments described above show that fragment release occurs rapidly (<1 min) after slicing. It is possible, however, that this estimated time for the release step reflects ribosome-stimulated release. Our earlier results show that the *first* ribosome promotes AGO2-target binding by unmasking the target site (see Figure 2K), but does not exclude the possibility that a *following* ribosome stimulates fragment release by colliding with AGO2 after catalysis has occurred. To determine the release rate in the absence of (possible) ribosome-AGO2 collisions, we set out to determine the kinetics of the cleavage cycle of a transcript translated by a single ribosome. We introduced a 5’ UTR sequence of the EMI1 gene in our reporter (Figure 3D), which strongly represses translation initiation (Tanenbaum et al., 2015; Yan et al., 2016). We selected mRNAs that were translated by a single ribosome and used exonucleolytic decay of the 3’ cleavage fragment (i.e. disappearance of the mCherry spot) as a read-out for mRNA cleavage, since translation termination and mRNA cleavage cannot be distinguished based on GFP-mCherry foci separation for mRNAs translated by a single ribosome (see methods) (Hoek et al., 2019). We compared the duration of the entire cleavage cycle (from target site unmasking until exonucleolytic decay of the 3’ fragment) of the EMI1-5’UTR-KIF18B reporter with the KIF18B reporter and found very similar cleavage cycle durations for both reporters (Figures 3E and 3F). These results demonstrate that fragment release is not substantially affected by ribosomes colliding with AGO2 after cleavage and confirm that fragment release occurs rapidly after mRNA slicing *in vivo*. Finally, these results also show that a single ribosome translating the siRNA binding site is sufficient to stimulate binding site accessibility for AGO2/siRNA complexes.

### Interactions of the AGO2 target sequence with flanking mRNA sequences drive target site masking

Translating ribosomes can promote binding site accessibility either by displacing RBPs from the binding site, or by unfolding structure(s) in the mRNA that mask the AGO2 binding site. To distinguish between RBP displacement and mRNA structure unfolding, we designed a new set of reporters in which ribosomes can unfold mRNA structure without displacing RBPs from the AGO2 binding site. In these new reporters, the siRNA binding site is placed close to the 3’ end of the mRNA, immediately upstream of the PCP binding site array. The PCP binding sites are well-folded RNA hairpins and thus likely unavailable for base-pairing with the AGO2 binding site. Therefore, structures masking the AGO2 binding site will mostly arise from interactions between the AGO2 binding site and upstream mRNA sequences. Stop codons were introduced into the coding sequence (a fusion of BFP and luciferase, referred to as luciferase reporters) either 27 nt or 110 nt upstream of the siRNA binding site (‘late stop’ reporter) (Figure 4A). In these reporters ribosomes can disrupt interactions of the AGO2 binding site with upstream mRNA sequences, without displacing RBPs from the binding site. As controls, we generated reporters in which the stop codon is positioned downstream of the siRNA binding site (‘downstream stop’ reporter) to remove both structures and RBPs, or reporters with a stop codon 1677 nt upstream of the binding site (‘early stop’ reporters), to remove neither structures nor RBPS (Figure 4A). If RNA structures are masking the AGO2 binding site, then the ‘late stop’ and ‘downstream stop’ reporters should be cleaved efficiently, while the early stop should not. In contrast, if RBPs mask the binding site, only the ‘downstream stop’ reporter should be cleaved efficiently. For these experiments, we selected the target sites of KIF18B siRNAs #1 and #2, and GAPDH siRNA #3, as each of these target sites showed strong stimulation of cleavage by ribosomes (i.e. target site masking) in their native contexts (see Figure 2D).

**Figure 4.**
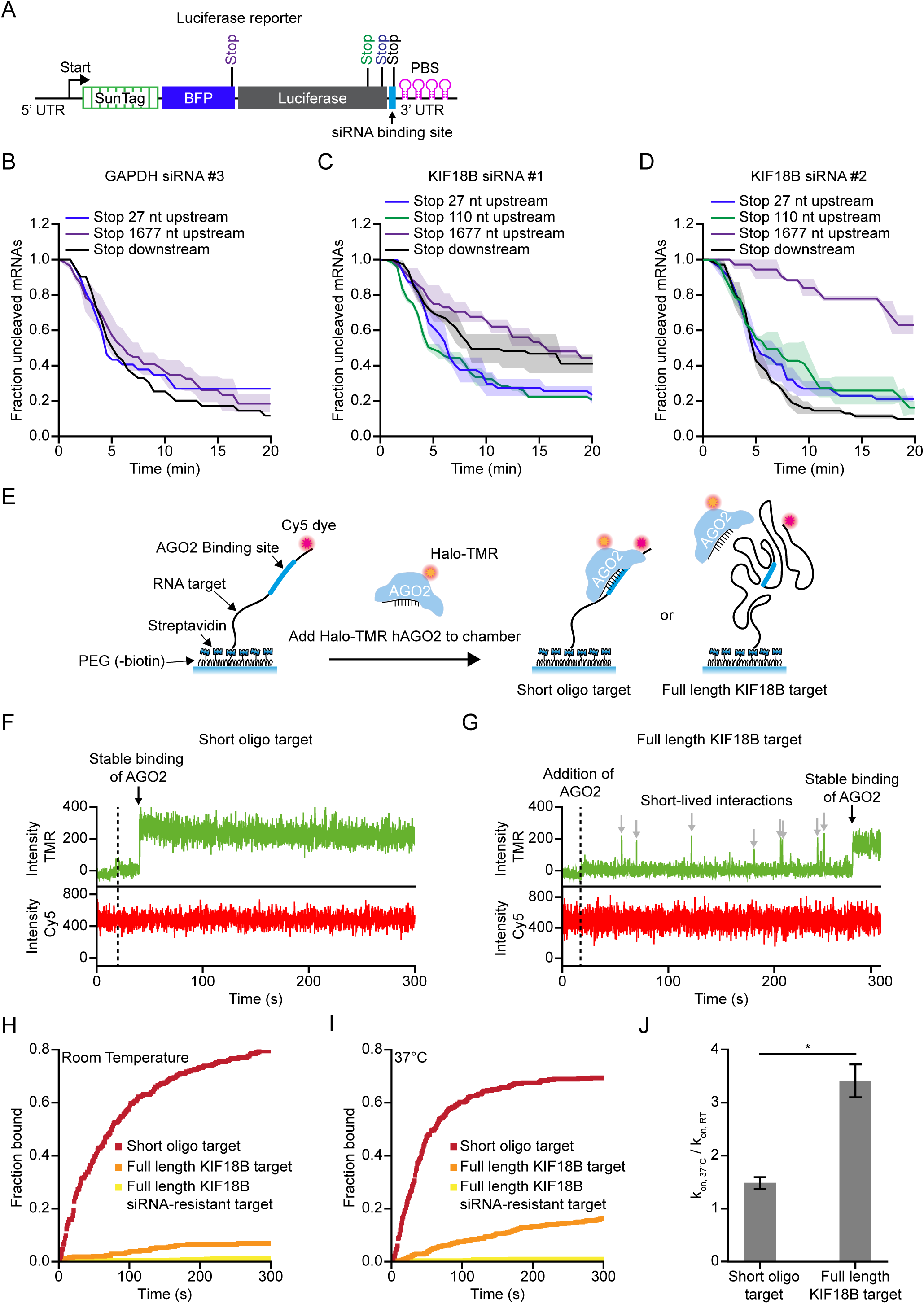
Masking of mRNA target sites by RNA structures inhibits AGO2-target interactions. (A) Schematic of the luciferase reporter. Position of different stop codon are indicated. (B-D) SunTag/PP7 cells expressing the indicated reporters were transfected with 10 nM of the indicated siRNA. The time from first detection of translation until separation of GFP and mCherry foci (i.e. mRNA cleavage) is shown. Solid lines and corresponding shaded regions represent mean ± SEM. (E) Schematic of the *in vitro* single-molecule binding assay. (F, G) Representative traces of AGO2/siRNA complex binding to the short oligo target (F) or the full length KIF18B target (G). Green line represents Halo-TMR AGO2 (top) and red line represents Cy5 RNA signal (bottom). Grey arrows indicate short binding events by AGO2 and black arrow indicates stable binding by the AGO2/siRNA complex. (H, I) The cumulative fraction of target RNAs bound by Halo-TMR AGO2 is plotted as a function of time for the indicated reporters at room temperature (RT) (H) and 37°C (I). (J) Ratio of kon at 37° C and RT for the short oligo target and the full length KIF18B target. P-value is based on a two-tailed Student’s t-test. Number of measurements for each experiment is listed in Table S1. See also Figure S3

mRNAs containing the GAPDH siRNA binding site showed fast cleavage rates in the context of all three stop codons, suggesting that, in these new reporters, the GAPDH siRNA #3 binding site is not masked (Figure 4B). In contrast, for the reporters containing the KIF18B siRNA #1 or #2 binding sites, cleavage was substantially faster for the ‘downstream stop’ reporters compared with the ‘early stop’ reporters (Figures 4C and 4D, compare black and purple lines), indicating strong target site masking. Importantly, the cleavage rates of the ‘late stop’ reporters were at least as fast as cleavage rates of the ‘downstream stop’ reporter (Figures 4C and 4D, compare blue and green to black lines), indicating that ribosomes stimulate AGO2 mRNA binding and cleavage by unfolding mRNA secondary structure, rather than displacing RBPs from the binding site. Surprisingly, for the KIF18B siRNA #1, the rate of cleavage of the ‘early stop’ reporters was even faster than that of the ‘downstream stop’ reporter (Figure 4C, compare blue and green lines to black line). A possible explanation for this result is that ribosomes passing over the AGO2 binding site impair mRNA cleavage by displacing AGO2 from the mRNA upon collision before slicing has occurred.

Interestingly, while cleavage by all three siRNAs (KIF18B siRNAs #1 and #2, and GAPDH siRNA #3) showed strong stimulation by ribosomes in their native context, the GAPDH siRNA #3 is no longer ribosome-stimulated in the above-mentioned luciferase reporter (Figure 4B). Furthermore, in the KIF18B reporter KIF18B siRNA #1 shows a stronger ribosome-dependent stimulation of cleavage than KIF18B siRNA #2 (see Figures 2C and S2C), while in the luciferase reporter KIF18B siRNA #2 is stimulated more strongly (Figures 4C and 4D). These findings suggest that the interactions between the AGO2 binding site sequence and flanking mRNA sequences determines the degree of binding site masking. Indeed, when AGO2 binding sites were inserted in different mRNAs and at different positions in an mRNA, the magnitude of target site masking (i.e. the ribosome-dependent cleavage stimulation) was variable (Figures S3A-G).

To directly test the role of flanking sequence in AGO2 target site masking, we established an *in vitro* assay to study AGO2-target interactions. We purified recombinant human AGO2, loaded it with KIF18B siRNA #1, and synthesized either a short RNA oligonucleotide or the full length KIF18B mRNA as target. As a control, we mutated the siRNA binding site to generate a KIF18B-siRNA-resistant mRNA as target. To visualize AGO2-target interactions directly, AGO2 was fluorescently labeled using a HaloTag and TMR dye, and the RNA targets were conjugated to a single Cy5 dye (Figure 4E). Fluorescently-labeled targets were biotinylated and conjugated to a glass slide (Figure 4E) (Chandradoss et al., 2015). AGO2/siRNA complexes were then perfused into the sample chamber and the binding of fluorescently-labeled AGO2 to either the oligo target or full length KIF18B targets was assessed at room temperature (RT) (Figures 4F and 4G). AGO2 binding to the oligo target occurred very rapidly (t1/2 = 73 ± 8 s, mean ± SD), while binding to the full length mRNA target occurred much slower (t1/2 = 4.1 ± 0.7 × 10^3^ s, mean ± SD) (Figure 4H). As the target sequence was identical in the short and long targets, and as no other RBPs were present in the reaction, these results suggest that RNA structures formed in the full length transcript inhibit binding of AGO2 to the target site. Since RNA folding is strongly dependent on temperature (with higher temperature resulting in reduced RNA folding), we repeated the binding assay at 37°C and found that AGO2 bound to the full length KIF18B target 3.4-fold faster at 37°C, while binding to the oligo target was only 1.5-fold faster at 37°C (Figures 4I and 4J), further indicating that interactions of the AGO2 target site with nucleotides control AGO2 binding site availability.

### Multiple weak intramolecular mRNA interactions result in potent AGO2 target site masking

Next, we sought to characterize the type of RNA structures that cause AGO2 binding site masking. We performed structure prediction using mfold (Zuker, 2003) to determine the nature of the structures that mask the AGO2 target sites. We focused on the GAPDH reporter and GAPDH siRNA #3, due to its strong structural masking and the relatively small size of the GAPDH reporter, which simplifies RNA structure predictions. We generated a new reporter (‘mfold’ reporter) that contained 19 nucleotide substitutions in the mRNA sequence flanking the target site, which disrupted all the strongest predicted RNA structures involving the AGO2 binding site (see methods). Analysis of the cleavage kinetics of the ‘mfold’ reporter revealed that mRNA cleavage was still strongly stimulated by ribosomes (Figure 5A), indicating that the AGO2 binding site was still masked by RNA structures in the mutated reporter mRNA.

**Figure 5:**
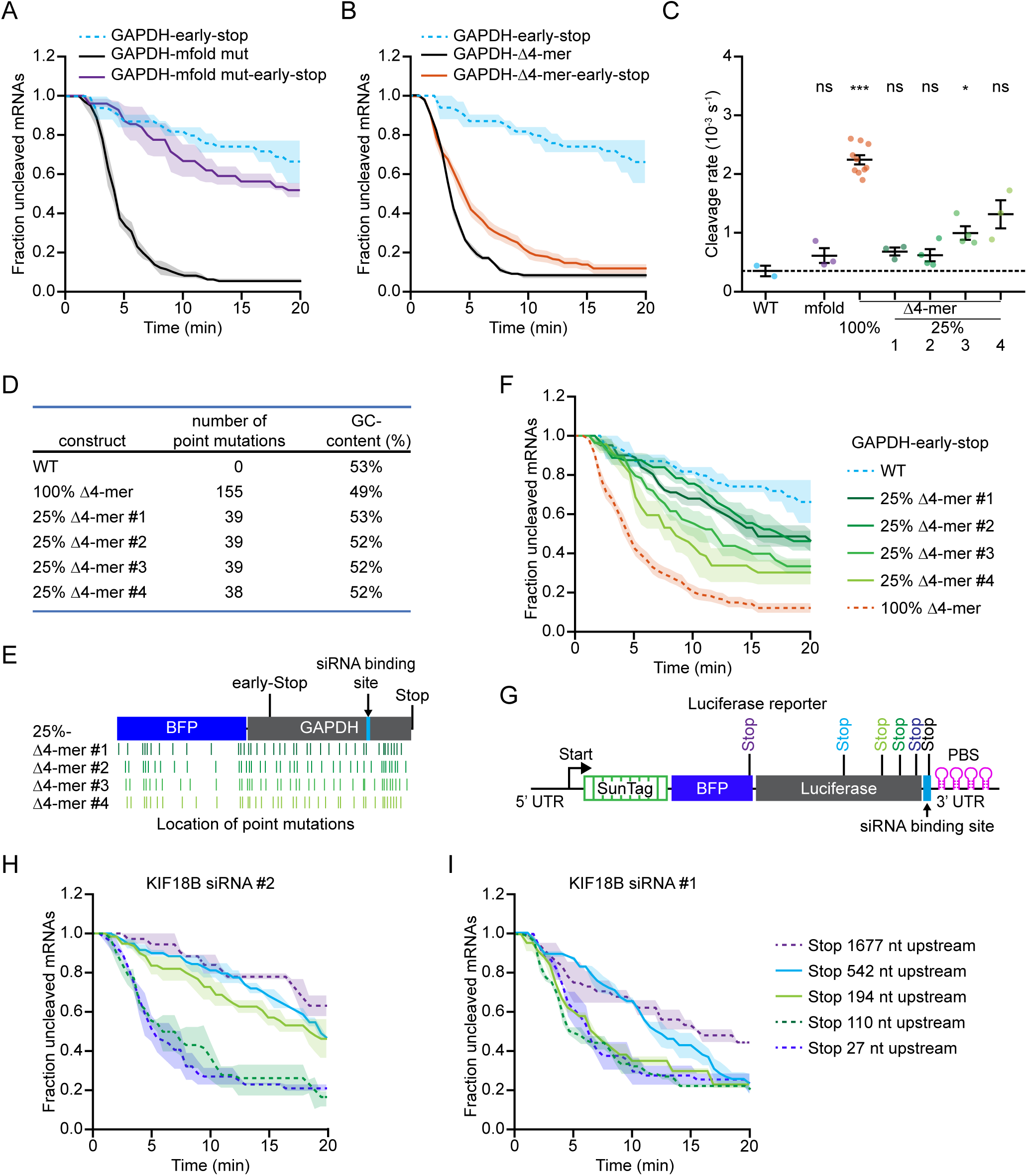
Multiple weak intramolecular mRNA interactions cooperatively mask AGO2-target sites. (A, B, F, H, I) SunTag/PP7 cells expressing the indicated reporters were transfected with 10 nM siRNA of GAPDH siRNA #3 (A, B, F) or indicated siRNAs (H, I). The time from first detection of translation until separation of GFP and mCherry foci (i.e. mRNA cleavage) is shown. Solid lines and corresponding shaded regions represent mean ± SEM. Dotted lines indicate that the data is replotted from an earlier figure panel for comparison. (C) Calculated cleavage rates in the absence of ribosomes translating the siRNA target site are shown for indicated reporters treated with GAPDH siRNA #3 (see methods). Each dot represents a single experiment and lines with error bars indicate the mean ± SEM. P-values are based on a two-tailed Student’s t-test. P-values are indicated as * (p < 0.05), ** (p < 0.01), *** (p < 0.001), ns = not significant. (D) Characteristics of the different Δ4-mer reporters. (E) Schematic overview of the location of the single nucleotide substitutions in the 25% Δ4-mer reporters. (G) Schematic overview of the luciferase reporters used in (H, I) containing stop codons at variable distances from the siRNA binding site. Number of measurements for each experiment is listed in Table S1. See also Figure S4

Since disrupting the strongest predicted structures involving the AGO2 binding site did not prevent robust structural masking of the binding site, it is likely that binding site masking is not due to a single strongly folded structure. Instead, AGO2 binding site masking may arise from numerous weak interactions between the AGO2 binding site and short complementary nucleotide sequences in the target mRNA. To test this hypothesis, we mutated, through single nucleotide substitutions, all 4-mer sequences in the GAPDH mRNA reporter that showed complementarity to the binding site of GAPDH siRNA #3 (105 single nucleotide substitutions in GAPDH and 50 in the BFP upstream of the GAPDH coding sequence, referred to as the ‘Δ4-mer’ reporter, see methods). Removal of 4-mers substantially increased the cleavage rate in the absence of ribosomes translating the AGO2 binding site (∼6 fold) (Figures 5B, 5C, compare blue and orange dots), indicating that AGO2 binding site masking was largely disrupted in this mutant mRNA. Removal of complementary 4-mers for KIF18B siRNA #1 in the KIF18B reporter also substantially reduced ribosome-dependent stimulation of mRNA cleavage, although residual cleavage stimulation could still be observed (Figure S4A), possibly due to other short sequences with complementarity (e.g. 3-mers or 6 or 7-mers with single mismatches). Together, these results confirm that ribosomes stimulate AGO2-target interactions by unfolding mRNA structures, and suggest that multiple nucleotide sequences, each with weak affinity for the AGO2 binding site, together drive strong target site masking.

Next, we generated four new ‘Δ4-mer’ reporters; in each of these reporters a non-overlapping set of 25% of the single nucleotide substitutions were introduced that disrupt the complimentary 4-mers (Δ25% 4-mers) (Figures 5D and 5E; see methods). If only a small number of all 4-mer sequences that are mutated in the ‘Δ4-mer’ reporter are involved in AGO2 target site masking, then a subset of the ‘Δ25% 4-mer’ reporters should show a strong reduction in target site masking, while others should be unaffected. In contrast, if many different 4-mers provide a small contribution to target site masking, all four reporters should show a *partial* reduction in target site masking. Analysis of cleavage kinetics revealed that all four ‘Δ25% 4-mer’ reporters showed an intermediate phenotype (Figures 5F, 5C, compare blue and green dots, and S4B-E), demonstrating that multiple different 4-mer sequences are cooperatively contributing to AGO2 target site masking. The heterogeneity in RNA structures that mask the AGO2 binding site (i.e. multiple different 4-mers can bind to the target site), as well as the low affinity of 4-mer interactions suggest that structures causing target site masking may not be readily identified using *in silico* RNA folding predictions or sequencing-based structure determinations.

Next, we sought to map the distances over which flanking sequences can act to mask the AGO2 target site. For this, we made use of the ‘luciferase’ reporters containing the stop codon either 27 nt, 110 nt, or 1677 upstream of the AGO2 target site (Figure 5G). As the ‘late stop’ reporters showed no structural masking of the target site, while the ‘early stop’ reporter showed strong structural masking (see Figures 4C and 4D), sequences located between these stop codons are likely contributing to the observed AGO2 binding site masking. To identify the positions of these sequences, we generated additional reporters in which we gradually moved the stop codon further away from the AGO2 binding site (Figure 5G) and assessed mRNA cleavage of these additional reporters. For the reporter containing KIF18B siRNA #2, moving the stop codon 194 nt from the AGO2 binding site already caused substantial target site masking. Moving the stop even farther away (542 nt from AGO2 binding site) further increased target site masking, resulting in a similar cleavage rate as observed for the ‘early stop’ reporter (Figures 5H and S4F). Similar effects were observed when using KIF18B siRNA #1, although the relative contribution of different regions of the mRNA that were involved in AGO2 target site masking were different (Figures 5I and S4F), consistent with a different nucleotide composition of the two target sites. These results show that structures spanning several hundred nucleotides can contribute to AGO2 target site masking (Figure S4F), consistent with other studies showing that base-pairing interactions can occur over large distances (Lu et al., 2016; Metkar et al., 2018).

### Kinetics of mRNA folding shape AGO2-mRNA interactions dynamics

While several methods are available to generate ‘snapshots’ of RNA structures (Aw et al., 2016; Bevilacqua et al., 2016; Ding et al., 2014; Gong et al., 2018; Lu et al., 2016; Mustoe et al., 2018; Rouskin et al., 2014; Strobel et al., 2018; Wan et al., 2014; Zubradt et al., 2017), very little is known about the structural dynamics of mRNAs *in vivo*. The dynamics of mRNA (un-)folding are likely important, as we find that structural unmasking of binding sites is a key driver of AGO2-target interactions. We therefore wished to characterize mRNA structural dynamics in more detail.

Inhibiting ribosome translocation by addition of CHX results in a decreased cleavage rate (see Figures 2C and S2C-I), suggesting that mRNAs refold after ribosome-dependent unfolding. We reasoned that upon inhibition of ribosome translocation, the cleavage rate will decrease over time as re-folding of the target site occurs. Thus, a transition from a fast cleavage rate to a slow cleavage rate is expected after CHX addition, and the time over which this transition occurs, reports on the mRNA folding rate. To assess the kinetics of mRNA refolding, we first determined that CHX slows ribosome translocation within 30-60 s after drug addition to the cell culture medium (Figure S5A). Next, we re-analyzed the mRNA cleavage kinetics upon CHX addition for a subset of reporters. In our initial analysis, we included all translating mRNAs, however, a subset of the analyzed mRNAs initiated translation just before cleavage occurred, so the first ribosome had not yet reached (and unfolded) the siRNA binding site for those mRNAs. Cleavage kinetics of those mRNAs therefore does not report on the refolding rate of the structure surrounding the AGO2 target site. In our re-analysis, we only assessed the cleavage kinetics of mRNAs on which the first ribosome had recently arrived at the siRNA binding site (i.e. mRNAs for which the target site is expected to be mostly in an unfolded state upon CHX addition; referred to as ‘unfolded mRNAs’). For 3 out of 4 reporter-siRNA combinations, we indeed observed a fast initial cleavage rate after CHX addition, followed by a slower cleavage rate at later time-points (Figures 6A and S5B-D, red lines). Fitting these cleavage curves with a double exponential decay distribution revealed that the fast cleavage rate occurred with a half-life of around 1-2 min for different reporters (Figures 6A, S5B, S5C, dotted lines, and S5E). After correction for the delay in ribosome stalling upon addition of CHX, (30-60 s) (Figure S5A), these results suggest that AGO2 target sites are masked again within ∼30-90 s of ribosome-dependent unfolding. For the fourth reporter-siRNA condition (GAPDH siRNA #3) the cleavage rate was substantially faster for the set of ‘unfolded mRNAs’ when compared to all mRNAs for all time points (compare Figures S5D, red line, and S2G, purple line), suggesting that the target site remains in a (partially) unmasked state after unfolding in the presence of stalled ribosomes for this reporter. Possibly, stalled ribosomes near the target site inhibit mRNA re-folding for this region of the mRNA.

**Figure 6:**
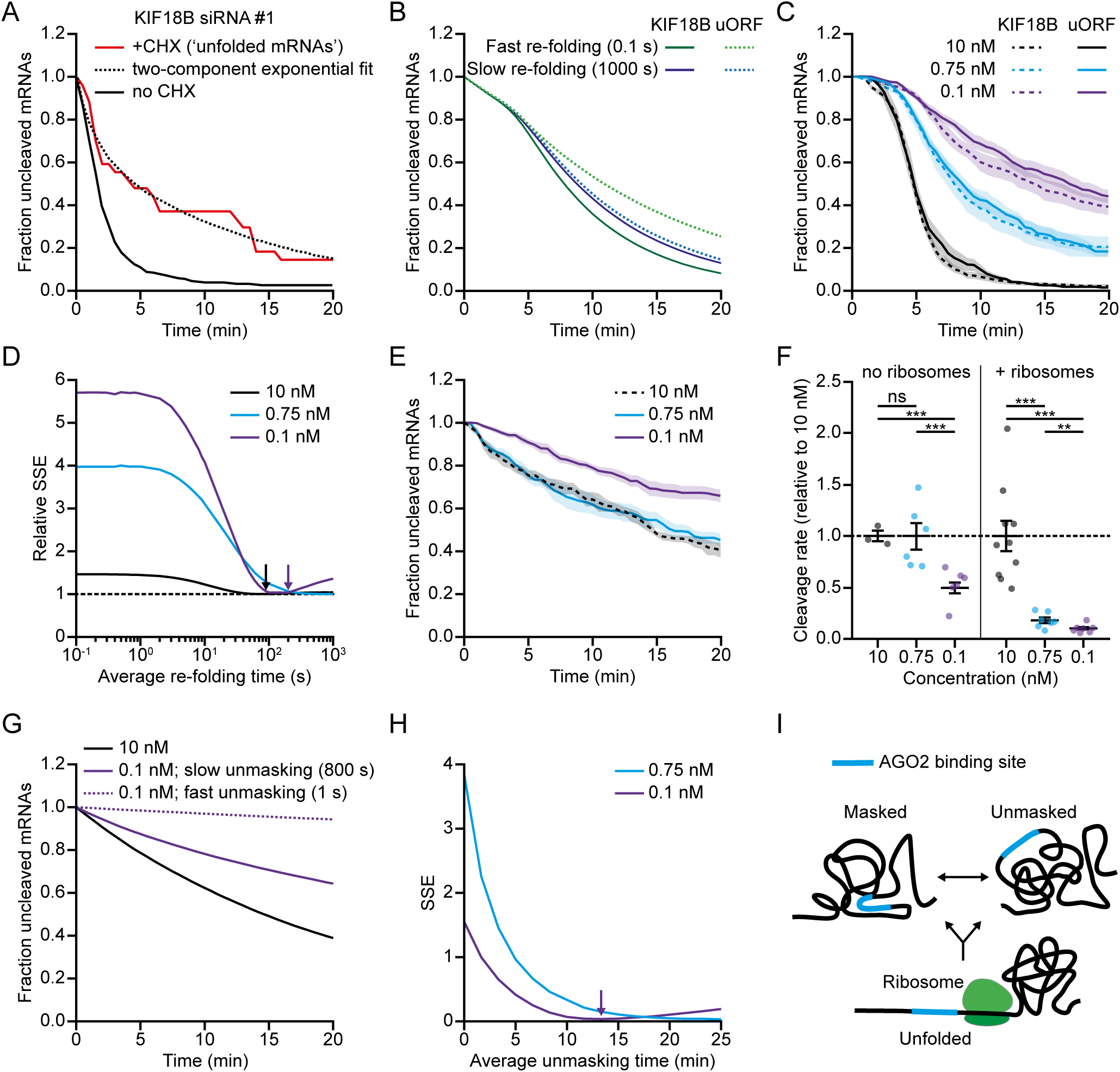
Kinetics of mRNA folding shape AGO2-mRNA interactions. (A) SunTag/PP7 cells expressing the KIF18B reporter were transfected with 10 nM siRNA KIF18B #1 and treated with CHX, where indicated. Only mRNAs for which translation initiated between 3.5-6.5 min before CHX addition were included (see methods). The time since CHX addition is shown for the ‘+CHX’ cleavage curve. Dotted line represents optimal fit with a two-component exponential decay distribution. The no CHX cleavage curve is re-normalized and plotted from 3.5 min after the start of translation. (B) Simulated cleavage curves for the KIF18B reporter and uORF reporter simulated with a fast or slow mRNA re-folding time. (C, E) SunTag/PP7 cells expressing indicated reporters were transfected with either 10 nM, 0.75 nM or 0.1 nM of KIF18B siRNA #1 and treated without (C) or with (E) CHX. The time from first detection of translation (C) or CHX addition (E) until separation of GFP and mCherry foci (i.e. mRNA cleavage) is shown. Solid lines and corresponding shaded regions represent mean ± SEM. Dotted lines indicate that the data is replotted from an earlier figure panel for comparison. (D) Goodness-of-fit score (relative SSE) at different simulated mRNA re-folding times for the data shown in (C) at indicated siRNA concentrations (see methods). Arrows indicate re-folding time of best fit. (F) Calculated cleavage rates for different siRNA concentrations (relative to 10 nM siRNA) for the data shown in (C, E). Each dot represents a single experiment and lines with error bars indicate the mean ± SEM. P-values are based on a two-tailed Student’s t-test. P-values are indicated as * (p < 0.05), ** (p < 0.01), *** (p < 0.001), ns = not significant. (G) Simulated cleavage curves for 10 and 0.1 nM siRNA concentration using fast or slow unmasking rates (average unmasking time of 1 s and 800 s, respectively). (H) Goodness-of-fit score (SSE) at different simulated unmasking times (see methods). Arrow indicates unmasking time of the best fit. (I) Schematic model of AGO2 target site masking/unmasking and the role of ribosomes in target site unfolding. Number of measurements for each experiment is listed in Table S1. See also Figure S5.

To examine mRNA re-folding kinetics after ribosome-induced unfolding through an independent method, we examined the relationship between the translation initiation rate and the mRNA cleavage rate. We previously found that a ribosome initiates translation every ∼25 s on these reporters (Yan et al., 2016) (and thus will pass the siRNA binding site every ∼25 s). If mRNA re-folding occurs very rapidly after unfolding by a translating ribosome (<25 s), then the mRNA will be refolded by the time the next ribosome arrives at the target site. In contrast, if the mRNA re-folds relatively slowly (>25 s), the target site will still be unmasked when the next ribosome arrives. Since the cleavage rate depends on the fraction of time that the target site is unmasked, the cleavage rate can be used as a proxy for the structural state of the mRNA. For example, if an mRNA re-folds within 60 s after unfolding by a ribosome, and a new ribosome initiates every 25 s, the target site will be mostly in an unfolded state during steady state translation. If the initiation rate is reduced to one ribosome per 50 s, the mRNA will still be unfolded most of the time, and the cleavage rate would be largely unchanged. However, if the initiation rate is reduced further, to one ribosome per 100 s, the mRNA will frequently re-fold before the next ribosome arrives and the target site will be in a masked state a substantial fraction of the time, which will reduce the cleavage rate. Thus, by measuring the cleavage rate at different translation initiation rates, one can estimate the folding rate of the mRNA sequence involved in AGO2 target site masking.

We introduced an upstream open reading frame (uORF) into the KIF18B reporter (‘uORF-KIF18B’ reporter), which reduced translation initiation by 3.3 fold (Figure S5F; see methods). The mean time interval between ribosomes passing the AGO2 binding site was thus increased from ∼25 s to ∼80 s. We then developed a computational framework to simulate mRNA cleavage rates at different theoretical mRNA folding rates and translation initiation rates (see methods). As expected, at fast re-folding rates the simulations predict a relatively large difference between the cleavage rate of the KIF18B and ‘uORF-KIF18B’ reporter, while a small difference in cleavage rate is predicted when simulating the cleavage rate using slow mRNA re-folding rate as an input (Figure 6B). We then measured mRNA cleavage rates for both the KIF18B and ‘uORF-KIF18B’ reporters and compared these cleavage rates to the simulated rates using a goodness-of-fit score (sum of squared errors (SSE)) (Figures 6C and 6D). Both experimental measurements and simulations were performed for three different siRNA concentrations, 10 nM, 0.75 nM and 0.1 nM to increase the accuracy of the analysis. For both the 10 nM and 0.1 nM, the optimal fit was achieved when simulating an mRNA re-folding time of ∼30-180 s, while a somewhat slower re-folding time (>180 s; a precise value could not be given due to the absence of a local minimum) was found for the 0.75 nM condition (Figure 6D). Overall, these results are in good agreement with the measurements of mRNA re-folding upon CHX treatment (30-90 s). These mRNA re-folding rates suggest that the AGO2 target site is likely unmasked for the majority of time when placed in the coding sequence of a gene which is translated at a moderate to high translation rate (e.g. the KIF18B reporter mRNA). These results provide the first real-time *in vivo* measurements of mRNA folding kinetics during translation and reveal how the translation initiation rate can affect mRNA target binding by AGO2.

When positioned in non-translated regions of the mRNA (i.e. 3’ UTR), AGO2 binding sites are not unfolded by ribosomes and structural unmasking of the target site must occur through alternative mechanisms. One possibility is that mRNA structures rearrange (stochastically) over time and that AGO2 target sites are only masked in a subset of all possible structural configurations. However, little is known about the dynamics of these structural rearrangements (i.e. masking and unmasking of the target site), which are likely important for AGO2-target binding in non-translated regions of the mRNA. If structural rearrangements occur on a timescale that is much faster than AGO2-target binding (∼1-18 min for 0.1-10 nM siRNA, see Figure 3B), AGO2-target binding will be rate-limiting for mRNA cleavage, and the mRNA cleavage rate will depend primarily on the AGO2/siRNA concentration. In contrast, if structural rearrangements occur at rates similar to or slower than AGO2-target binding, structural unmasking becomes an (additional) rate-limiting step, and the mRNA cleavage rate will become less sensitive to the siRNA concentration. Thus, the sensitivity of the mRNA cleavage rate to the siRNA concentration reports on the dynamics of mRNA structural rearrangements. Interestingly, while in the presence of translating ribosomes the cleavage rate is strongly affected by changes in the siRNA concentration, the cleavage rate in the absence of ribosomes showed a substantially weaker dependency on siRNA concentration (Figures 6E-F, and S5G), suggesting that target site unmasking through structural rearrangements becomes a rate-limiting step in the absence of translating ribosomes.

To quantitatively investigate the dynamics of ribosome-independent structural rearrangements, we used a computational approach. We simulated the effect of decreasing the siRNA concentration on the mRNA cleavage rate in the absence of ribosomes (see methods). For slow structural dynamics, we found that the simulated mRNA cleavage rate is less sensitive to siRNA concentration (0.1-10 nM) than for fast dynamics (Figure 6G, compare the solid and dotted purple lines to the black line), which is more consistent with the experimental data (Figure 6E, compare the purple line to the black line). To determine the unmasking time of our reporter mRNA, we compared the experimental cleavage curves at different siRNA concentrations (0.75 nM and 0.1 nM) to multiple simulated cleavage curves (each with different unmasking times) using a goodness-of-fit score (SSE). This analysis revealed that the best fits were obtained with unmasking times of >10 min (Figure 6H), indicating that target site unmasking becomes a rate-limiting step in mRNA cleavage, even at higher concentrations of siRNA (between 0.75 nM and 10 nM). Furthermore, these simulations indicate that target site unmasking in the 3’ UTR (i.e. in the absence of translating ribosomes) is much slower (>10 min) than the unmasking rate in the coding sequence (where target sites are unfolded every ∼25 s by a translating ribosome for our reporters), highlighting the importance of ribosome-mediated unmasking of AGO2 target sites.

### Discussion

In this study, we use a live-cell imaging approach to visualize translation and AGO2-mediated cleavage of individual mRNA molecules. This work provides *in vivo* measurements of AGO2 cleavage kinetics, reveals that the interplay between AGO2 and ribosomes controls AGO2 cleavage efficiency, and more broadly, reveals how structural dynamics of mRNAs shape AGO-target interactions.

### Mechanisms of target repression by AGO2

Several types of mRNA target repression have been proposed for AGO family members, including direct mRNA cleavage, or recruitment of additional proteins that mediate translational repression and/or mRNA decay. Here, we developed a new single-molecule imaging approach to investigate mRNA target silencing by AGO2 complexed with an siRNA in living human somatic cells. We find that AGO2 efficiently silences its target mRNA through endonucleolytic cleavage, rapidly after nuclear export of the mRNA. We did not find evidence for mRNA cleavage in the nucleus or inhibition of target mRNA transcription by AGO2, nor did we observe translational repression of target mRNAs. However, it is important to note that AGO2/siRNA complexes associate with their target only very briefly (∼1 min), as mRNA slicing and fragment release occur rapidly upon target site binding (see Figure 3B). Translational repression may require longer association of AGO proteins with their target, as recruitment of additional proteins is likely required for translational repression (Filipowicz et al., 2008; Jonas and Izaurralde, 2015). It will be interesting to apply our single-molecule imaging assay to visualize target silencing by other classes of small RNAs (e.g. miRNAs or piRNAs) to determine the kinetics of other types of target silencing, including translational repression.

### Kinetics of the AGO2 cleavage cycle

*In vitro* studies have aimed at determining the kinetics of each step in AGO2’s cleavage cycle (Ameres et al., 2007; Chandradoss et al., 2015; Jo et al., 2015; Lam et al., 2015; Salomon et al., 2015; Wee et al., 2012; Yao et al., 2015), but little was known about the cleavage kinetics in cells. Our study provides the first *in vivo* measurements of the kinetics of the cleavage cycle of AGO2. AGO2 binds its target site in <1 min at high (10 nM) siRNA concentration if the target site is unmasked (as is the case for target sites in the coding sequence). However, the average time to binding is substantially longer at lower siRNA concentrations (∼8 min at 0.75 nM siRNA and ∼18 min at 0.1 nM siRNA) (see Figure 3B) or for target sites that are frequently masked (as is the case for many target sites in 3’UTRs). After the initial AGO2-target interaction, the remaining steps of the cleavage cycle (conformational rearrangements of AGO2, mRNA slicing, and fragment release) occur within ∼1 min of target binding (see Figure 3B), indicating that in most cases initial binding of AGO2 to its target is the rate-limiting step.

### Paradoxical roles of ribosomes in controlling AGO2-mRNA target interactions

Recent reports showed that ribosomes reduce the overall degree of structure in the coding sequence of the transcriptome (Adivarahan et al., 2018; Beaudoin et al., 2018; Mizrahi et al., 2018; Mustoe et al., 2018). In addition, several studies have shown that AGO target sites are less efficiently recognized if they are embedded within a strong structure (Ameres et al., 2007; Beaudoin et al., 2018). Here, we show that ribosome-dependent unfolding of mRNA structures stimulates AGO-target interactions, providing a direct, causal link between mRNA translation and AGO2-target binding. The observation that siRNA-mediated mRNA cleavage is more efficient in an actively translated region appears to contrast previous reports that miRNAs repress their target more efficiently when bound to the 3’UTR (Grimson et al., 2007; Gu et al., 2009). It is possible that ribosomes also inhibit AGO-target interactions, for example by displacing AGO from the mRNA through physical collisions (Bartel, 2009; Grimson et al., 2007). Thus, ribosomes may have two opposing activities that affect AGO-target interactions and the net effect of ribosomes on AGO-dependent target silencing may depend on a number of different factors.

First, the kinetics of target silencing by AGO may determine whether ribosomes have a stimulatory or inhibitory role. We find that AGO2/siRNA complexes cleave their target relatively fast after initial binding. Therefore, cleavage will frequently occur before AGO2-ribosome collisions can displace AGO2 from the mRNA. In contrast, if miRNAs act more slowly, the frequency of ribosome-AGO collisions is increased, which in turn could increase the inhibitory effect of ribosomes and make 3’ UTRs more effective target sites for miRNAs. Second, the net effect of ribosomes depends on the degree of target site masking. We observed that target site masking, and thus the stimulatory role of ribosomes, varies greatly for different AGO2 binding sites and corresponding flanking sequences. Finally, the translation initiation rate likely affects both the kinetics of mRNA unfolding and the frequency of ribosome-AGO collisions, and therefore also controls the net effect of ribosomes on AGO target repression.

Interestingly, previous analysis revealed that miRNA target sites positioned immediately downstream of the stop codon are highly active (Grimson et al., 2007). Consistent with this, we find that AGO binds very efficiently to target sites positioned immediately downstream of the stop codon, because ribosomes translating upstream mRNA sequences can stimulate unmasking of target sites immediately downstream of the stop codon (see Figures 4C and 4D). Therefore, binding sites just downstream of the stop codon may benefit from the stimulatory activity of ribosomes, while being protected from the inhibitory effect of ribosome-AGO collisions. In summary, we find that ribosomes may have a paradoxical role in controlling AGO-target interactions. This paradoxical role of ribosomes may not be limited to AGO family proteins, but may broadly shape the interactions of RBPs with their target RNAs. Future work will hopefully reveal the role of ribosomes and mRNA structural dynamics in controlling RBP-mRNA interactions and uncover the parameters that determine whether ribosomes stimulate or inhibit other RBP-mRNA interactions.

### mRNA structural dynamics and heterogeneity

Many AGO2 binding sites tested in our study showed substantial target site masking. Removing the strongest *predicted* structures involving the binding site did not eliminate target site masking (see Figure 5A). In contrast, mutation of all or subsets of short complementary 4-mer sequences in the target mRNA reduced target site masking (see Figure 5F), indicating that multiple (or many) sequences in the target mRNA cooperatively contribute to target site masking and that different structural configurations exist that can mask the AGO2 target site.

Individual 4-mer base-pair interactions have very rapid binding and unbinding kinetics. Surprisingly though, our data suggest that target sites can remain masked for >10 min in the absence of ribosomes (see Figure 6H). So how can we reconcile these two apparently contradictory findings? One speculative model (Figure 6I) is that mRNAs form stable 3-dimensional structures in which multiple sequences with weak/moderate affinity for the AGO2 target site are positioned in close proximity to the target site, resulting in frequent interactions and strong target site masking. In this model, the key activity of ribosomes would be to unfold the stable 3-dimensional structure that facilitates target site masking, rather than directly disrupting target site interactions with complementary sequences. In the absence of translating ribosomes such structures could persist for long periods of time (>10 min), explaining the very slow cleavage kinetics for some reporters in which the AGO2 binding is located in the 3’ UTR (e.g. see Figures 2C and S2G). Possibly, these structures stochastically rearrange over time, naturally resulting in AGO2 target site unmasking (without complete unfolding of the mRNA), or perhaps structures are unfolded or refolded sporadically by cellular helicases. Together, these results provide a high temporal resolution analysis of the structural dynamics of an mRNA molecule *in vivo* and provide a framework for understanding the role of mRNA structural dynamics in shaping RBP-mRNA interactions.

## Acknowledgements

We thank Martin Depken for helpful discussions with the computational modelling. We thank Loes Steller, Iris Bally, and Rupa Banerjee for help with experiments. We would also like to thank the Tanenbaum lab members for helpful discussions, and Tim Hoek and Deepak Khuperkar for critical reading of the manuscript. This work was financially supported by the European Research Council (ERC) through an ERC starting grant (ERCSTG 677936-RNAREG) to M.E.T., a VENI grant from the Netherlands Organization for Scientific Research (NWO) (NWO 016.VENI.171.050) to S.R., an ERC consolidator grant (819299) and a VIDI grant from NWO (864.14.002) to C.J., and the National Institute of General Medical Sciences (R35 GM127090) to I.J.M.; M.E.T., S.R., S.S., D.d.S and I.L. are supported by the Oncode Institute that is partly funded by the Dutch Cancer Society (KWF).

## Author contributions

S.R., S.S., and M.E.T. conceived the project; S.R., S.S., I.L., and D.d.S. performed the *in vivo* experiments and analyzed the data; S.S. performed the computational modeling; T.J.C. performed the *in vitro* experiments and analyzed the data under supervision of C.J.; Y.X. purified the hAGO2 complex under supervision of I.J.M.; S.R., S.S., and T.J.C. prepared the figures; S.R., S.S. and M.E.T. wrote the manuscript and T.J.C, Y.X., I.J.M., and C.J. provided input.

## Figure legends

**Supplemental figure S1.**
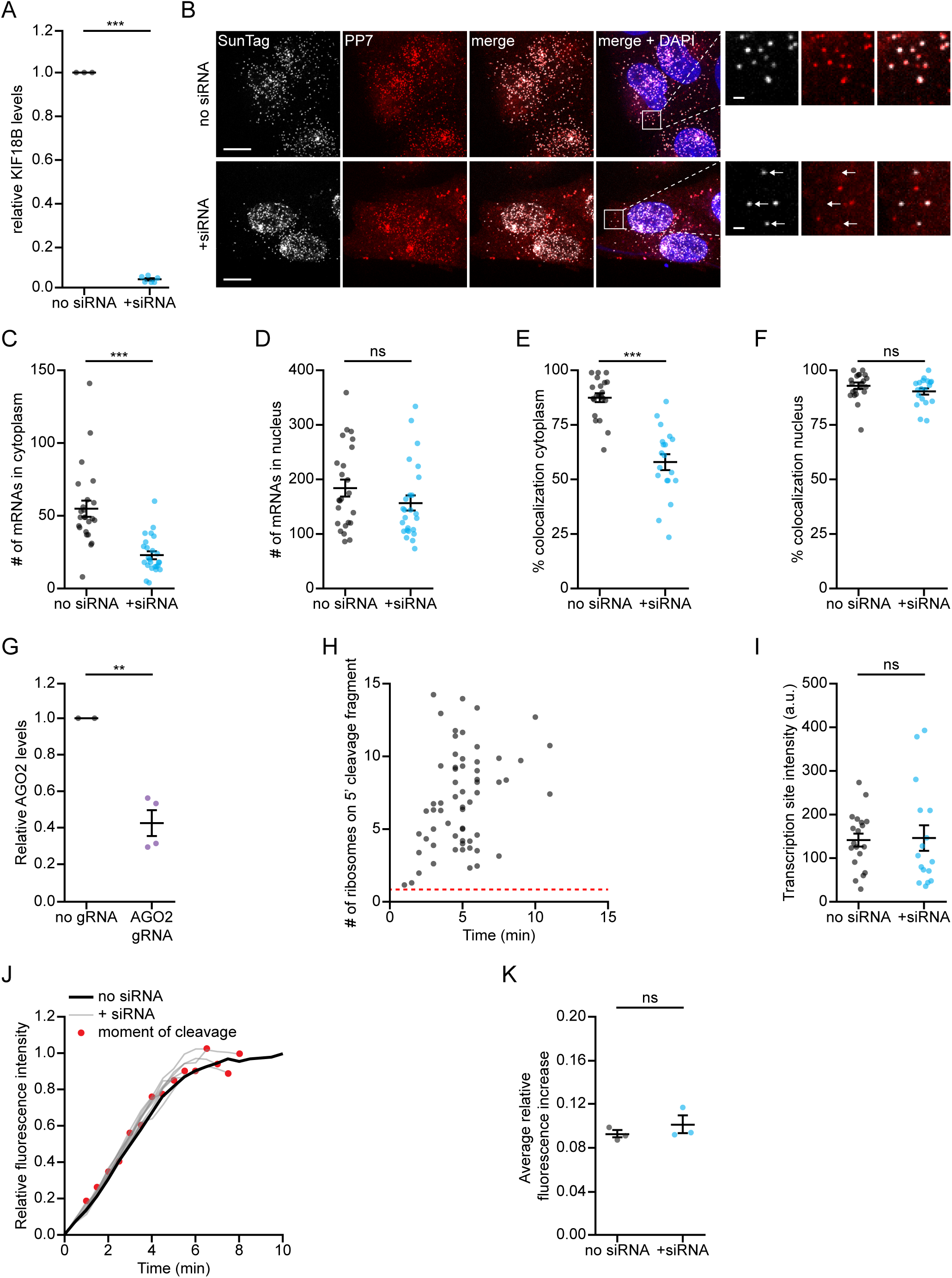
Effects of AGO2/siRNA complexes on mRNA transcription and translation - related to Figure 1. (A) Relative mRNA levels of endogenous KIF18B based on qPCR in non-transfected cells (no siRNA) and cells transfected with KIF18B siRNA #1 (+ siRNA). Each dot represents an independent experiment and lines with error bars indicate the mean ± SEM. (B-F, I) Cells expressing the KIF18B reporter without siRNA (no siRNA) or transfected with 10 nM KIF18B siRNA #1 (+ siRNA) were fixed and incubated with smFish probes to visualize reporter mRNAs. (B) Representative images of cells incubated with smFISH probes targeting the KIF18B reporter (SunTag-Cy5 and PP7-Alexa594) in no siRNA cells (upper panel) and + siRNA cells (lower panel). Arrows in insets indicate mRNA molecules for which the 5’ end (SunTag-Cy5 probe) and 3’ end (PP7-Alexa594 probe) do not co-localize. Scale bar, 10 µm in large images and 1 µm in insets. (C-D) Number of mRNAs in no siRNA and + siRNA cells in the cytoplasm (C) and the nucleus (D) determined based on smFISH using probes targeting the SunTag sequence. Each dot represents a single cell and lines with error bars indicate the mean ± SEM. (E-F) Percentage of mRNAs for which the 5’ end (labeled with SunTag probes) and 3’ end (labeled with PP7 probes) co-localized in no siRNA and + siRNA cells, either in the cytoplasm (E) or in the nucleus (F). Each dot represents a single cell and lines with error bars indicate the mean ± SEM. (G) Relative AGO2 mRNA levels based on qPCR in control cells (no gRNA) and cells treated with a CRISPRi guide targeting endogenous AGO2 (AGO2 gRNA). Each dot represents an independent experiment and lines with error bars indicate the mean ± SEM. (H) SunTag/PP7 cells expressing the KIF18B reporter were transfected with KIF18B siRNA #1. The number of ribosomes present on the 5’ cleavage fragment was determined one frame after the moment of cleavage (see methods). Dotted red line indicates the intensity of a single SunTag array (i.e. the intensity associated with a single ribosome). (I) Cells were treated for 40 min with dox and the integrated intensity of transcription sites was determined with smFISH probes targeting the SunTag sequence. Each dot represents a single transcription site and lines with error bars indicate the mean ± SEM. (J-K) SunTag/PP7 cells expressing the KIF18B reporter were untransfected (no siRNA) or transfected with KIF18B siRNA #1 (+ siRNA). (J) GFP intensity over time associated with individual mRNAs is shown for no siRNA cells (black line) and + siRNA cells (grey lines). Black line indicates average of all mRNAs in no siRNA cells, while each grey line represents the average GFP intensity of all mRNAs cleaved at the same moment relative to the start of translation (see methods). The red dot indicates the moment of cleavage. (K) Average increase in GFP fluorescence intensity either between 1.5-4 min after the start of translation (no siRNA) or at the moment preceding mRNA cleavage (+ siRNA) is shown (see methods). Each dot represents the average of an independent experiment and lines with error bars indicate the mean ± SEM. (A, C-F, G, I) P-values are based on a two-tailed Student’s t-test. (K) P-value is based on a paired two-tailed t-test. P-values are indicated as * (p < 0.05), ** (p < 0.01), *** (p < 0.001), ns = not significant. Number of measurements for each experiment is listed in Table S1.

**Supplemental figure S2.**
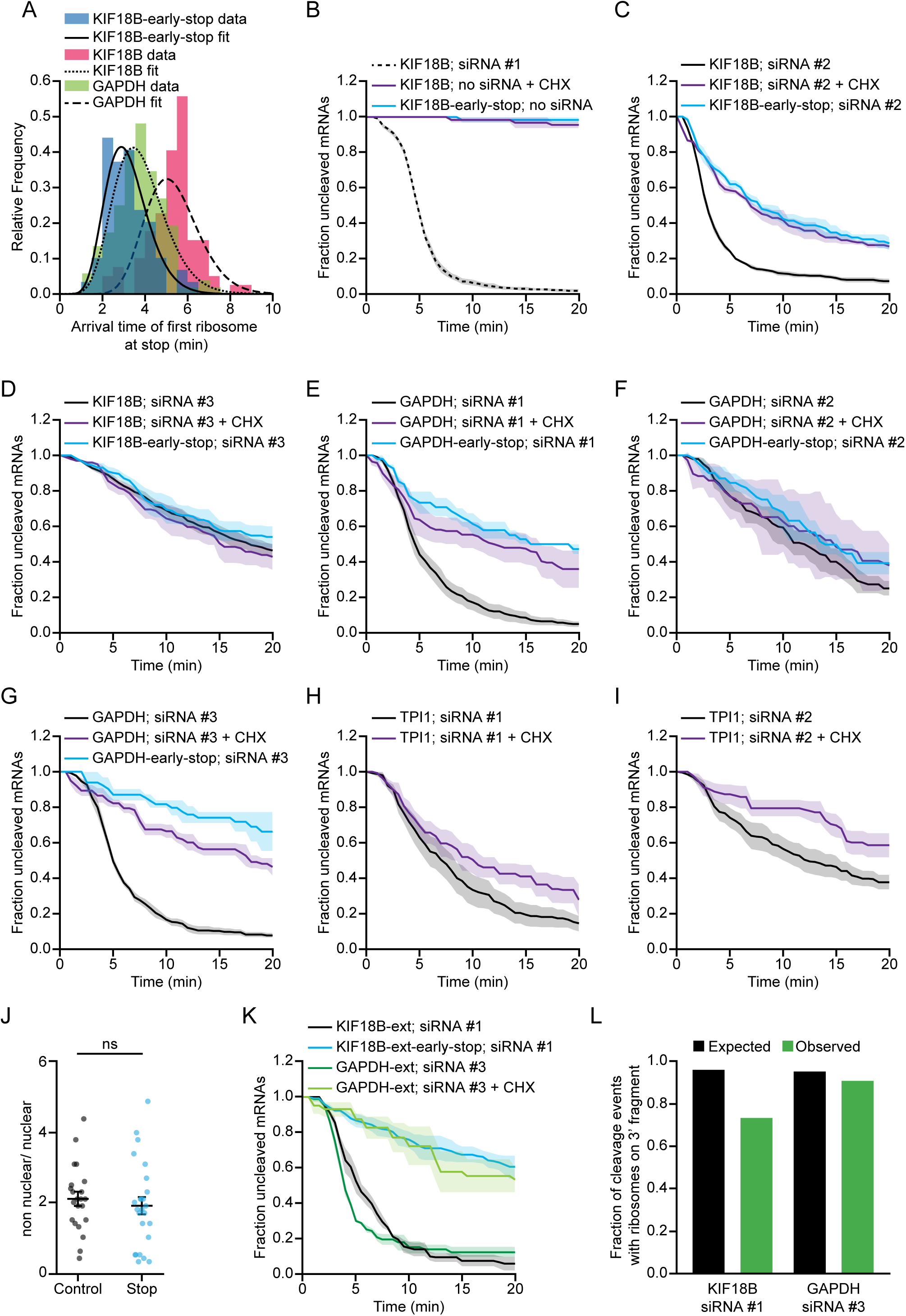
Ribosomes stimulate AGO2-dependent mRNA cleavage - related to Figure 2. (A) The moment at which the first ribosome arrived at the stop codon was calculated for indicated reporters. The experimental data (colored bars) was fit with a gamma distribution (black lines) (See methods). (B-I) SunTag/PP7 cells expressing the indicated reporters were transfected with 50 nM (KIF18B siRNA #3) or 10 nM (all others) siRNA and treated with CHX, where indicated. The time from first detection of translation or from CHX addition until separation of GFP and mCherry foci (i.e. mRNA cleavage) is shown. Solid lines and corresponding shaded regions represent mean ± SEM. Dotted line indicates that the data is replotted from an earlier figure panel for comparison. (J) Ratio of non-nuclear and nuclear mRNAs 90 min after addition of dox in cells expressing the KIF18B reporter (control) or KIF18B-early-stop reporter (Stop) as determined by smFISH using SunTag probes. Note that mRNA localization is similar for the two cell lines used for northern blot analysis (see Figure 2E). Each dot represents one cell and lines with error bars indicate the mean ± SEM. P-value is based on a two-tailed Student’s t-test. (K) SunTag/PP7 cells expressing the indicated reporters were transfected with 10 nM siRNA and treated with CHX, where indicated. The time from first detection of translation or from CHX addition (+ CHX) until separation of GFP and mCherry foci (i.e. mRNA cleavage) is shown. Solid lines and corresponding shaded regions represent mean ± SEM. (L) The fraction of mRNAs that contains a ribosome on the 3’ cleavage fragment is shown for mRNAs on which translation started at least 7.5 minutes (KIF18B) or 6 minutes (GAPDH) before the moment of cleavage. On these mRNAs it is expected that the first ribosome has passed the AGO2 target site in ∼95% of mRNAs (indicated by black bars) based on the experimentally-derived ribosome elongation rate. The expected fraction (black bars) and observed fraction (green bars) of mRNAs that contains a ribosome on the 3’ cleavage fragment is shown. Number of measurements for each experiment is listed in Table S1.

**Supplemental Figure S3.**
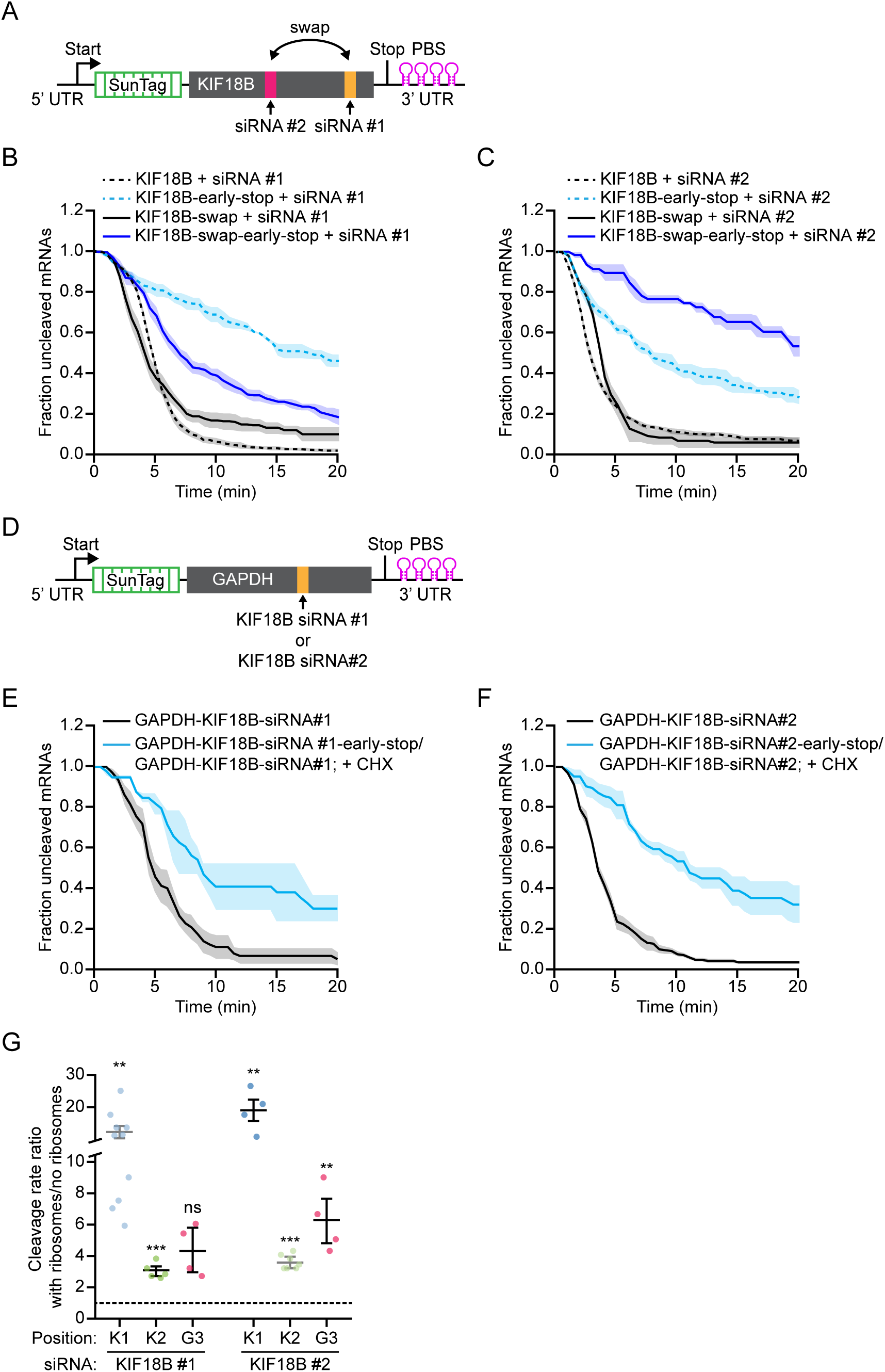
Degree of structural masking depends on the AGO2 binding sequence and the surrounding sequence - related to Figure 4. (A) Schematic of the KIF18B reporter in which the position of the siRNA #1 and siRNA #2 binding sites are swapped. (B, C) SunTag/PP7 cells expressing indicated reporters were transfected with 10 nM KIF18B siRNA #1 (B) or 10 nM KIF18B siRNA #2 (C). The time from first detection of translation until separation of GFP and mCherry foci (i.e. mRNA cleavage) is shown. Solid lines and corresponding shaded regions represent mean ± SEM. Dotted lines indicate that the data is replotted from an earlier figure panel for comparison. (D) Schematic of the GAPDH reporter in which the KIF18B siRNA #1 or KIF18B siRNA #2 binding site is placed at the position of GAPDH siRNA #3. (E, F) SunTag/PP7 cells expressing the indicated reporters were transfected with 10 nM KIF18B siRNA #1 (E) or 10 nM KIF18B siRNA #2 (F). The time from first detection of translation or CHX addition until mRNA cleavage is shown. Note that data of the KIF18B-early-stop reporter and KIF18B reporter treated with CHX are combined to generate the cleavage curve for cleavage in the absence of ribosomes. Solid lines and corresponding shaded regions represent mean ± SEM. (G) Ratio of cleavage rate in the presence and absence of ribosomes is shown for the indicated siRNAs and reporters (see methods). Each dot represents a single experiment and lines with error bars indicate the mean ± SEM. P-values are based on a two-tailed Student’s t-test. P-values are indicated as * (p < 0.05), ** (p < 0.01), *** (p < 0.001). K1, K2 and G3 indicate the position of the indicated siRNA. K1 refers to the position of KIF18B siRNA #1, K2 to KIF18B siRNA #2 and G3 to GAPDH siRNA #3. Light blue and light green data points are replotted from an earlier experiment. Number of measurements for each experiment is listed in Table S1.

**Supplemental figure S4.**
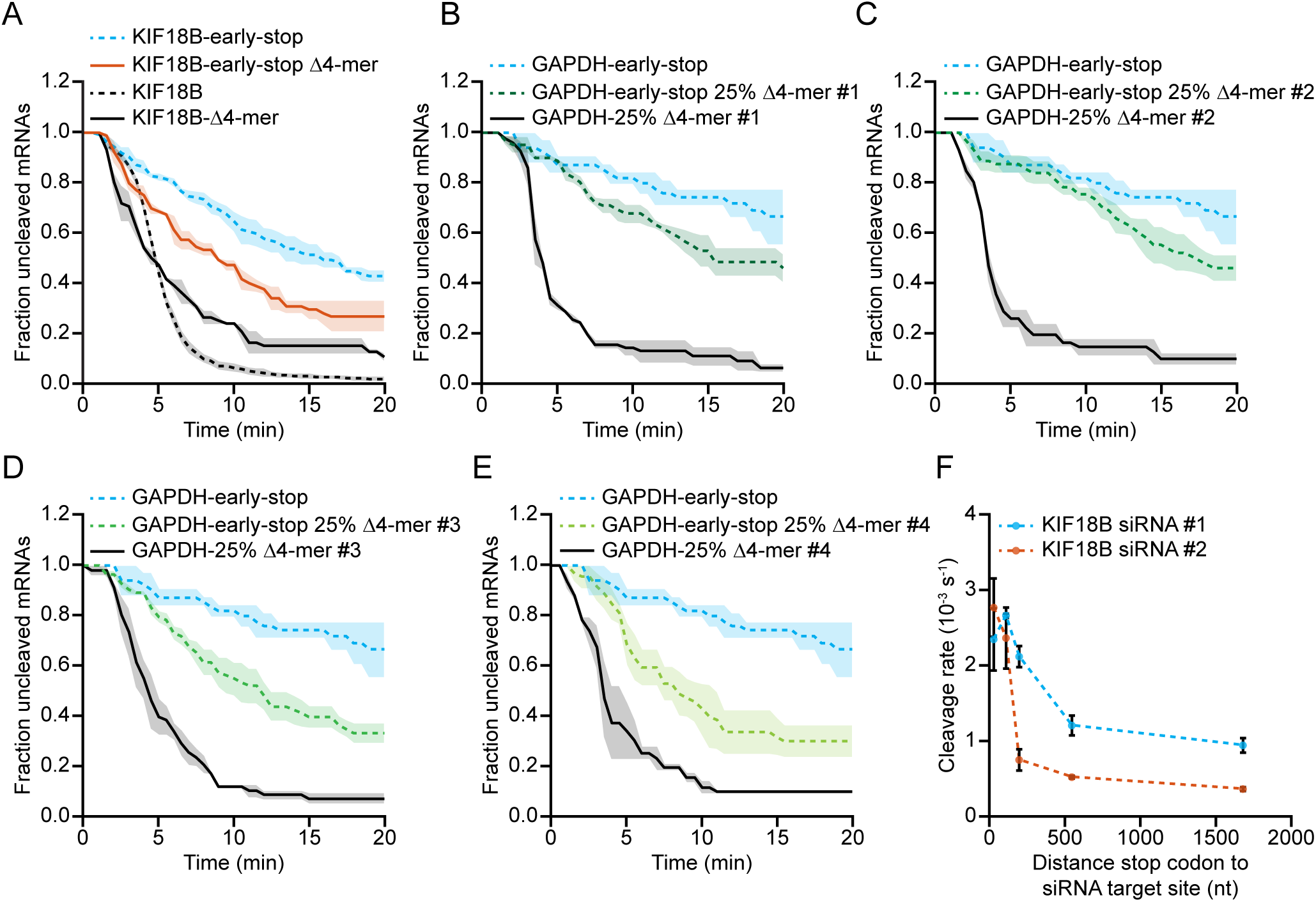
Multiple weak intramolecular mRNA interactions together result in potent AGO2 target site masking - related to Figure 5. (A-E) SunTag/PP7 cells expressing the indicated reporters were transfected with (A) 10 nM KIF18B siRNA #1 or (B-E) 10 nM GAPDH siRNA #3. The time from first detection of translation until separation of GFP and mCherry foci (i.e. mRNA cleavage) is shown. Solid lines and corresponding shaded regions represent mean ± SEM. Dotted lines indicate that the data is replotted from an earlier figure panel for comparison. (F) Cleavage rates for the ‘luciferase’ reporters with indicated siRNA target sites and with different distances between the stop codon and the siRNA target site are shown. Each dot and error bar indicate the mean ± SEM. Dotted lines are only for visualization. Number of measurements for each experiment is listed in Table S1.

**Supplemental figure S5.**
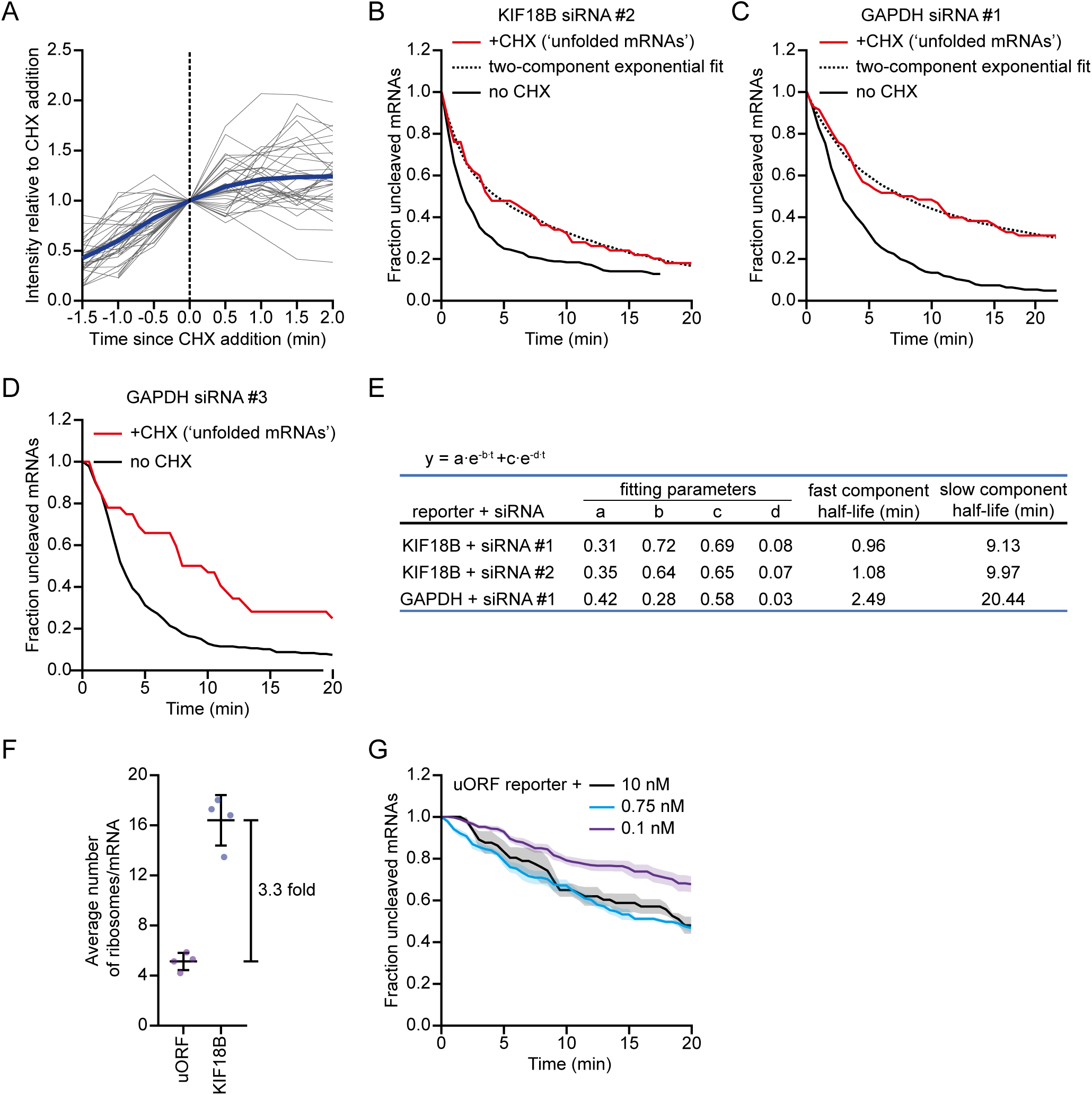
Structural dynamics of RNA folding - related to Figure 6. (A) Relative GFP fluorescence intensities were measured before and after the addition of CHX in SunTag/PP7 cells expressing the KIF18B reporter. Intensity-time traces were aligned at the moment of CHX addition. GFP fluorescence intensities were normalized to the GFP fluorescence intensities at the moment of CHX addition. The thick blue line represents the average intensity of all traces, thin grey lines represent intensity traces of multiple single mRNAs. (B-D) SunTag/PP7 cells expressing the indicated reporters were transfected with 10 nM of the indicated siRNA and treated with CHX, where indicated. The CHX cleavage curves (red lines) only include mRNAs for which translation started between 2.5-5.0 min (B), 2.0-4.5 min (C), or 2.0-5.0 min (D) before CHX addition (see methods). Dotted lines represent optimal fit with a two-component exponential decay distribution. The no CHX cleavage curve is re-normalized and plotted from 2.5 min (B) or 2.0 min (C-D) after the start of translation. (E) Fitting parameters and corresponding half-lives of the two-component exponential fits from Figures 6A and S5B-C. (F) Average number of ribosomes per mRNA molecule for the KIF18B-uORF and KIF18B reporters. Each dot represents an independent experiment and lines with error bars indicate the mean ± SEM. (G) SunTag/PP7 cells expressing the KIF18B-uORF reporter were transfected with either 10 nM, 0.75 nM or 0.1 nM of KIF18B siRNA #1 and treated with CHX. The time from CHX addition until mRNA cleavage is shown. Solid lines and corresponding shaded regions represent mean ± SEM. Number of measurements for each experiment is listed in Table S1.

## Supplemental items

**Table S1.** Related to all figures. Number of repeats, cells, and mRNAs per experiment. Overview of the number of repeats per experiment, the number of cells, and the number of mRNAs analyzed per experiment. Some datasets are used in multiple analyses, which is indicated in the last column (referred to as ‘also plotted in Figure’). Replotting of the same analysis is indicated in the figure legends.

| Overview | condition | repeats | cells | # of events/molecules | also plotted in Figure |
| --- | --- | --- | --- | --- | --- |
| 1E | KIF18B no siRNA | 3 | 21 | 110 |  |
|  | KIF18B; KIF18B siRNA #1 | 10 | 82 | 732 | 1F, 2B, 2C, 3A, 6C, S2B, S3B, S4A |
|  | KIF18B-siRNA-resistant; KIF18B siRNA #1 | 2 | 22 | 85 |  |
| 1F | KIF18B; KIF18B siRNA #1 | 10 | 82 | 732 | 1E, 2B, 2C, 3A, 6C, S2B, S3B, S4A |
|  | KIF18B; AGO2 knockdown + KIF18B siRNA #1 | 4 | 45 | 305 |  |
|  | KIF18B; AGO2 knockdown + AGO2 overexpression + KIF18B siRNA #1 | 2 | 24 | 124 |  |
| 2B | KIF18B; KIF18B siRNA #1 | 10 | 82 | 732 | 1E, 1F, 2C, 3A, 6C, S2B, S3B, S4A |
|  | spacer-KIF18B; KIF18B siRNA #1 | 2 | 9 | 48 |  |
| 2C | KIF18B; KIF18B siRNA #1 | 10 | 82 | 732 | 1E, 1F, 2B, 3A, 6C, S2B, S3B, S4A |
|  | KIF18B-early-stop (1050 nt up. of siRNA site); KIF18B siRNA #1 | 3 | 26 | 150 | S3B, S4A |
|  | KIF18B; KIF18B siRNA #1 + CHX | 3 | 39 | 178 | 6E |
|  | KIF18B; KIF18B siRNA #1 + Eme | 3 | 11 | 38 |  |
| 3A | KIF18B; 10 nM KIF18B siRNA #1 | 10 | 82 | 732 | 1E, 1F, 2B, 2C, 6C, S2B, S3B, S4A |
| 3C | KIF18B; 0.75 nM KIF18B siRNA #1 | 7 | 161 | 530 | 6C |
|  | KIF18B; 0.1 nM KIF18B siRNA #1 | 12 | 106 | 692 | 6C |
| 3E | KIF18B; 10 nM KIF18B siRNA #1 | 2 | 23 | 86 |  |
|  | EMI1-5'UTR-KIF18B; 10 nM KIF18B siRNA #1 | 5 | 28 | 71 |  |
|  | EMI1-5'UTR-KIF18B; no siRNA | 2 | 20 | 51 |  |
| 4B | BFP-Luc-GAPDH siRNA site #3-early-stop (1677 nt up. of siRNA site); GAPDH siRNA #3 | 3 | 18 | 50 |  |
|  | BFP-Luc-GAPDH siRNA site #3-early-stop (27 nt up. of siRNA site); GAPDH siRNA #3 | 1 | 10 | 37 |  |
|  | BFP-Luc-GAPDH siRNA site #3; GAPDH siRNA #3 | 1 | 9 | 41 |  |
| 4C | BFP-Luc-KIF18B siRNA site #1-early-stop (1677 nt up. of siRNA site); KIF18B siRNA #1 | 2 | 13 | 38 | 5I |
|  | BFP-Luc-KIF18B siRNA site #1-early-stop (110 nt up. of siRNA site); KIF18B siRNA #1 | 2 | 15 | 59 | 5I |
|  | BFP-Luc-KIF18B siRNA site #1-early-stop (27 nt up. of siRNA site); KIF18B siRNA #1 | 2 | 20 | 65 | 5I |
|  | BFP-Luc-KIF18B siRNA site #1; KIF18B siRNA #1 | 2 | 17 | 58 |  |
| 4D | BFP-Luc-KIF18B siRNA site #2-early-stop (1677 nt up. of siRNA site); KIF18B siRNA #2 | 2 | 18 | 43 | 5H |
|  | BFP-Luc-KIF18B siRNA site #2-early-stop (110 nt up. of siRNA site); KIF18B siRNA #2 | 2 | 14 | 54 | 5H |
|  | BFP-Luc-KIF18B siRNA site #2-early-stop (27 nt up. of siRNA site); KIF18B siRNA #2 | 2 | 17 | 62 | 5H |
|  | BFP-Luc-KIF18B siRNA site #2; KIF18B siRNA #2 | 2 | 13 | 63 |  |
| 4H | short oligo target (RT) | 3 | n.a | 792 |  |
|  | full length KIF18B target (RT) | 3 | n.a | 762 |  |
|  | full length KIF18B target siRNA resistant (RT) | 3 | n.a | 1591 |  |
| 4I | short oligo target (37C) | 3 | n.a | 447 |  |
|  | full length KIF18B target (37C) | 3 | n.a | 756 |  |
|  | full length KIF18B target siRNA resistant (37C) | 3 | n.a | 919 |  |
| 5A | GAPDH-early-stop (518 nt up of siRNA site); GAPDH siRNA #3 | 2 | 27 | 60 | 5B, 5F, S2G, S4B, S4C, S4D, S4E |
|  | GAPDH-mfold mutant; GAPDH siRNA #3 | 3 | 23 | 98 |  |
|  | GAPDH-mfold mutant-early-stop (518 nt up of siRNA site); GAPDH siRNA #3 | 3 | 20 | 49 |  |
| 5B | GAPDH-early STOP (518 nt up of siRNA site); GAPDH siRNA #3 | 2 | 27 | 60 | 5A, 5F, S2G, S4B, S4C, S4D, S4E |
|  | GAPDH-Δ4 mer; GAPDH siRNA #3 | 3 | 21 | 119 |  |
|  | GAPDH-early-stop (518 nt up of siRNA site)-Δ4-mer; GAPDH siRNA #3 | 10 | 60 | 214 | 5F |
| 5F | GAPDH-early-stop (518 nt up of siRNA site); GAPDH siRNA #3 | 2 | 27 | 60 | 5A, 5B, S2G, S4B, S4C, S4D, S4E |
|  | GAPDH-early-stop (518 nt up of siRNA site) 25% Δ4mer #1; GAPDH siRNA #3 | 3 | 26 | 61 | S4B |
|  | GAPDH-early-stop (518 nt up of siRNA site) 25% Δ4mer #2; GAPDH siRNA #3 | 4 | 26 | 67 | S4C |
|  | GAPDH-early-stop (518 nt up of siRNA site) 25% Δ4mer #3; GAPDH siRNA #3 | 4 | 32 | 80 | S4D |
|  | GAPDH-early-stop (518 nt up of siRNA site) 25% Δ4mer #4; GAPDH siRNA #3 | 3 | 23 | 62 | S4E |
|  | GAPDH-early-stop (518 nt up of siRNA site) 100% Δ4-mer; GAPDH siRNA #3 | 10 | 60 | 214 | 5B |
| 5H | BFP-Luc-KIF18B siRNA site #2-early-stop (1677 nt up. of siRNA site); KIF18B siRNA #2 | 2 | 18 | 43 | 4D |
|  | BFP-Luc-KIF18B siRNA site #2-early-stop (542 nt up. of siRNA site); KIF18B siRNA #2 | 2 | 15 | 58 |  |
|  | BFP-Luc-KIF18B siRNA site #2-early-stop (194 nt up. of siRNA site); KIF18B siRNA #2 | 2 | 18 | 58 |  |
|  | BFP-Luc-KIF18B siRNA site #2-early-stop (110 nt up. of siRNA site); KIF18B siRNA #2 | 2 | 14 | 54 | 4D |
|  | BFP-Luc-KIF18B siRNA site #2-early-stop (27 nt up. of siRNA site); KIF18B siRNA #2 | 2 | 17 | 62 | 4D |
| 5I | BFP-Luc-KIF18B siRNA site #1-early-stop (1677 nt up. of siRNA site); KIF18B siRNA #1 | 2 | 13 | 38 | 4C |
|  | BFP-Luc-KIF18B siRNA site #1-early-stop (542 nt up. of siRNA site); KIF18B siRNA #1 | 3 | 18 | 58 |  |
|  | BFP-Luc-KIF18B siRNA site #1-early-stop (194 nt up. of siRNA site); KIF18B siRNA #1 | 2 | 7 | 43 |  |
|  | BFP-Luc-KIF18B siRNA site #1-early-stop (110 nt up. of siRNA site); KIF18B siRNA #1 | 2 | 15 | 59 | 4C |
|  | BFP-Luc-KIF18B siRNA site #1-early-stop (27 nt up. of siRNA site); KIF18B siRNA #1 | 2 | 20 | 65 | 4C |
| 6A | KIF18B; KIF18B siRNA #1 + CHX ('unfolded mRNAs') | 3 | 22 | 27 |  |
|  | KIF18B; KIF18B siRNA #1 (re-plotted from 3.5 min) | 10 | 80 | 587 |  |
| 6C | KIF18B; 10 nM KIF18B siRNA #1 | 10 | 82 | 732 | 1E, 1F, 2B, 2C, 3A, S2B, S3B, S4A |
|  | KIF18B; 0.75 nM KIF18B siRNA #1 | 7 | 161 | 530 | 3C |
|  | KIF18B; 0.1 nM KIF18B siRNA #1 | 12 | 106 | 692 | 3C |
|  | uORF-KIF18B; 10 nM KIF18B siRNA #1 | 3 | 36 | 332 |  |
|  | uORF-KIF18B; 0.75 nM KIF18B siRNA #1 | 6 | 158 | 483 |  |
|  | uORF-KIF18B; 0.1 nM KIF18B siRNA #1 | 10 | 109 | 594 |  |
| 6E | KIF18B; 10 nM KIF18B siRNA #1 + CHX | 3 | 39 | 134 | 2C |
|  | KIF18B; 0.75 nM KIF18B siRNA #1 + CHX | 6 | 113 | 196 |  |
|  | KIF18B; 0.1 nM KIF18B siRNA #1 + CHX | 9 | 96 | 243 |  |
| S1C | KIF18B; no siRNA | 1 | 23 | 1319 |  |
|  | KIF18B; KIF18B siRNA #1 | 1 | 24 | 577 |  |
| S1D | KIF18B; no siRNA | 1 | 23 | 4419 |  |
|  | KIF18B; KIF18B siRNA #1 | 1 | 24 | 3920 |  |
| S1E | KIF18B; no siRNA | 1 | 21 | 574 |  |
|  | KIF18B; KIF18B siRNA #1 | 1 | 19 | 405 |  |
| S1F | KIF18B; no siRNA | 1 | 21 | 721 |  |
|  | KIF18B; KIF18B siRNA #1 | 1 | 19 | 765 |  |
| S1I | KIF18B; no siRNA | 1 | 16 | n.a. |  |
|  | KIF18B; KIF18B siRNA #1 | 1 | 19 | n.a. |  |
| S1J | KIF18B; no siRNA | 3 | 69 | 224 |  |
|  | KIF18B; KIF18B siRNA #1 | 3 | 141 | 809 |  |
| S2A | KIF18B-early-stop (1050 nt up. of siRNA site) | 5 | 25 | 59 |  |
|  | KIF18B | 3 | 51 | 79 |  |
|  | GAPDH | 3 | 43 | 125 |  |
| S2B | KIF18B; KIF18B siRNA #1 | 10 | 82 | 732 | 1E, 1F, 2B, 2C, 3A, 6C, S3B, S4A |
|  | KIF18B; no siRNA +CHX | 2 | 19 | 70 |  |
|  | KIF18B-early-stop (1050 nt up. of siRNA site); no siRNA | 2 | 20 | 58 |  |
| S2C | KIF18B; KIF18B siRNA #2 | 7 | 63 | 436 | S3C |
|  | KIF18B; KIF18B siRNA #2 + CHX | 3 | 65 | 142 |  |
|  | KIF18B-early-stop (826 nt up. of siRNA site); KIF18B siRNA #2 | 3 | 23 | 121 | S3C |
| S2D | KIF18B; KIF18B siRNA #3 | 9 | 124 | 569 |  |
|  | KIF18B; KIF18B siRNA #3 + CHX | 3 | 47 | 101 |  |
|  | KIF18B-early-stop (1355 nt up. of siRNA site); KIF18B siRNA #3 | 3 | 22 | 130 |  |
| S2E | GAPDH; GAPDH siRNA #1 | 6 | 65 | 290 |  |
|  | GAPDH; GAPDH siRNA #1 + CHX | 5 | 48 | 92 |  |
|  | GAPDH-early-stop (235 nt up. of siRNA site); GAPDH siRNA #1 | 3 | 14 | 57 |  |
| S2F | GAPDH; GAPDH siRNA #2 | 3 | 19 | 139 |  |
|  | GAPDH; GAPDH siRNA #2 + CHX | 3 | 18 | 51 |  |
|  | GAPDH-early-stop (711 nt up. of siRNA site); GAPDH siRNA #2 | 3 | 17 | 72 |  |
| S2G | GAPDH; GAPDH siRNA #3 | 8 | 72 | 391 |  |
|  | GAPDH; GAPDH siRNA #3 + CHX | 3 | 32 | 70 |  |
|  | GAPDH-early-stop (518 nt up. of siRNA site); GAPDH siRNA #3 | 2 | 27 | 60 | 5A, 5B, 5F, S4B, S4C, S4D, S4E |
| S2H | TPI; TPI siRNA #1 | 5 | 81 | 249 |  |
|  | TPI; TPI siRNA #1 + CHX | 4 | 45 | 75 |  |
| S2I | TPI; TPI siRNA #2 | 6 | 95 | 216 |  |
|  | TPI; TPI siRNA #2 + CHX | 4 | 48 | 69 |  |
| S2J | KIF18B (control) | 1 | 22 | 7127 |  |
|  | KIF18B-early-stop (stop) | 1 | 25 | 4995 |  |
| S2K | KIF18B-extended; KIF18B siRNA #1 | 4 | 20 | 87 |  |
|  | KIF18B-extended-early-stop (1050 nt up. of siRNA site); KIF18B siRNA #1 | 3 | 33 | 121 |  |
|  | GAPDH-extended; GAPDH siRNA #3 | 3 | 19 | 87 |  |
|  | GAPDH-extended; GAPDH siRNA #3 + CHX | 3 | 15 | 28 |  |
| S3B | KIF18B; KIF18B siRNA #1 | 10 | 82 | 732 | 1E, 1F, 2B, 2C, 3A, 6C, S2B, S4A |
|  | KIF18B-early-stop (1050 nt up. of siRNA site); KIF18B siRNA #1 | 3 | 26 | 150 | 2C, S4A |
|  | KIF18B-swap; KIF18B siRNA #1 | 5 | 32 | 231 |  |
|  | KIF18B-swap-early-stop (1050 nt up of siRNA site); KIF18B siRNA #1 | 4 | 28 | 147 |  |
| S3C | KIF18B; KIF18B siRNA #2 | 7 | 63 | 436 | S2C |
|  | KIF18B-early-stop (826 nt up. of siRNA site); KIF18B siRNA #2 | 3 | 23 | 121 | S2C |
|  | KIF18B-swap; KIF18B siRNA #2 | 4 | 21 | 180 |  |
|  | KIF18B-swap-early-stop (826 nt up. of siRNA site); KIF18B siRNA #2 | 3 | 16 | 62 |  |
| S3E | GAPDH-KIF18B-siRNA site #1; KIF18B siRNA #1 | 4 | 13 | 64 |  |
|  | GAPDH-KIF18B-siRNA site #1-early-stop(518 nt up. of siRNA site)/CHX; KIF18B siRNA #1 | 2 | 9 | 40 |  |
| S3F | GAPDH-KIF18B-siRNA site #2; KIF18B siRNA #2 | 4 | 18 | 148 |  |
|  | GAPDH-KIF18B-siRNA site #2-early-stop(518 nt up. of siRNA site)/CHX; KIF18B siRNA #2 | 4 | 17 | 61 |  |
| S4A | KIF18B-early-stop (1050 nt up of siRNA site); KIF18B siRNA #1 | 3 | 26 | 150 | 2C, S3B |
| | KIF18B-early-stop (1050 nt up of siRNA site) Δ4 mer; KIF18B siRNA #1 | 3 | 10 | 34 | |
|  | KIF18B; KIF18B siRNA #1 | 10 | 82 | 732 | 1E, 1F, 2B, 2C, 3A, 6C, S2B, S3B |
| | KIF18B-Δ4 mer; KIF18B siRNA #1 | 2 | 13 | 55 | |
| S4B | GAPDH-early-stop (518 nt up of siRNA site); GAPDH siRNA #3 | 2 | 27 | 60 | 5A, 5B, 5F, S2G, S4C, S4D, S4E |
| | GAPDH-early-stop (518 nt up of siRNA site) 25% Δ4 mer #1; GAPDH siRNA #3 | 3 | 26 | 61 | 5F |
| | GAPDH 25% Δ4 mer #1; GAPDH siRNA #3 | 3 | 24 | 69 | |
| S4C | GAPDH-early-stop (518 nt up of siRNA site); GAPDH siRNA #3 | 2 | 27 | 60 | 5A, 5B, 5F, S2G, S4B, S4D, S4E |
| | GAPDH-early-stop (518 nt up of siRNA site) 25% Δ4 mer #2; GAPDH siRNA #3 | 4 | 26 | 67 | 5F |
| | GAPDH 25% Δ4 mer #2; GAPDH siRNA #3 | 2 | 13 | 44 | |
| S4D | GAPDH-early-stop (518 nt up of siRNA site); GAPDH siRNA #3 | 2 | 27 | 60 | 5A, 5B, 5F, S2G, S4B, S4C, S4E |
| | GAPDH-early-stop (518 nt up of siRNA site) 25% Δ4 mer #3; GAPDH siRNA #3 | 4 | 32 | 80 | 5F |
| | GAPDH 25% Δ4 mer #3; GAPDH siRNA #3 | 3 | 23 | 59 | |
| S4E | GAPDH-early-stop (518 nt up of siRNA site); GAPDH siRNA #3 | 2 | 27 | 60 | 5A, 5B, 5F, S2G, S4B, S4C, S4D |
| | GAPDH-early-stop (518 nt up of siRNA site) 25% Δ4 mer #4; GAPDH siRNA #3 | 3 | 23 | 62 | 5F |
| | GAPDH 25% Δ4 mer #4; GAPDH siRNA #3 | 2 | 14 | 50 | |
| S5A | KIF18B; CHX | 3 | 10 | 36 |  |
| S5B | KIF18B; KIF18B siRNA #2 (re-plotted from 2.5 min) | 7 | 61 | 242 |  |
|  | KIF18B; KIF18B siRNA #2 + CHX ('unfolded mRNAs') | 3 | 38 | 50 |  |
| S5C | GAPDH; GAPDH siRNA #1 (re-plotted from 2.0 min) | 5 | 56 | 224 |  |
|  | GAPDH; GAPDH siRNA #1 + CHX ('unfolded mRNAs') | 4 | 34 | 58 |  |
| S5D | GAPDH; GAPDH siRNA #3 (re-plotted from 2.0 min) | 8 | 69 | 360 |  |
|  | GAPDH; GAPDH siRNA #3 + CHX ('unfolded mRNAs') | 3 | 20 | 32 |  |
| S5F | KIF18B-uORF | 4 | >4 | 218 |  |
|  | KIF18B | 4 | >4 | 350 |  |
| S5G | uORF-KIF18B; 10 nM KIF18B siRNA #1 + CHX | 3 | 29 | 65 |  |
|  | uORF-KIF18B; 0.75 nM KIF18B siRNA #1 + CHX | 5 | 88 | 145 |  |
|  | uORF-KIF18B; 0.1 nM KIF18B siRNA #1 + CHX | 9 | 95 | 22 |  |

## Methods

### U2OS and HEK293T cell culture

Human U2OS cells and HEK293T cells (ATCC) were grown in DMEM (4.5g/L glucose, Gibco) containing 5% fetal bovine serum (Sigma-Aldrich) and 1% penicillin/streptomycin (Gibco). Cells were grown at 37°C and with 5% CO2. Where indicated, Cycloheximide (CHX) (ThermoFisher) was used at a final concentration of 200 µg/ml, and Emetine (Eme) (Sigma-Aldrich) was used at a final concentration of 100 µg/ml.

### Cells, Plasmids, Transfections and Lentiviral infections

#### Cells

Live-cell imaging experiments were performed using U2OS cells, stably expressing TetR, scFv-sfGFP and PCP-mCherry-CAAX (referred to as SunTag/PP7 cells) (Yan et al., 2016) as well as the reporter of interest. The smFISH imaging experiments were performed in a monoclonal cell line, stably expressing TetR, scFv-sfGFP, PCP-Halo-CAAX and the 24xGCN4-KIF18B-24xPP7 reporter. Northern blot experiments were performed using two monoclonal cell lines, both expressing TetR, scFv-sfGFP, PCP-Halo-CAAX and in addition either the 24xGCN4-KIF18B-24xPP7 reporter or the 24xGCN4-KIF18B-early-stop-PP7 reporter.

#### Plasmids

The complete list and sequence of all plasmids used in this study is provided in Table S1 and Document S1.

For reporters where a stop codon was introduced upstream of the siRNA binding site, the distance between the stop codon and the siRNA binding site was as follows:

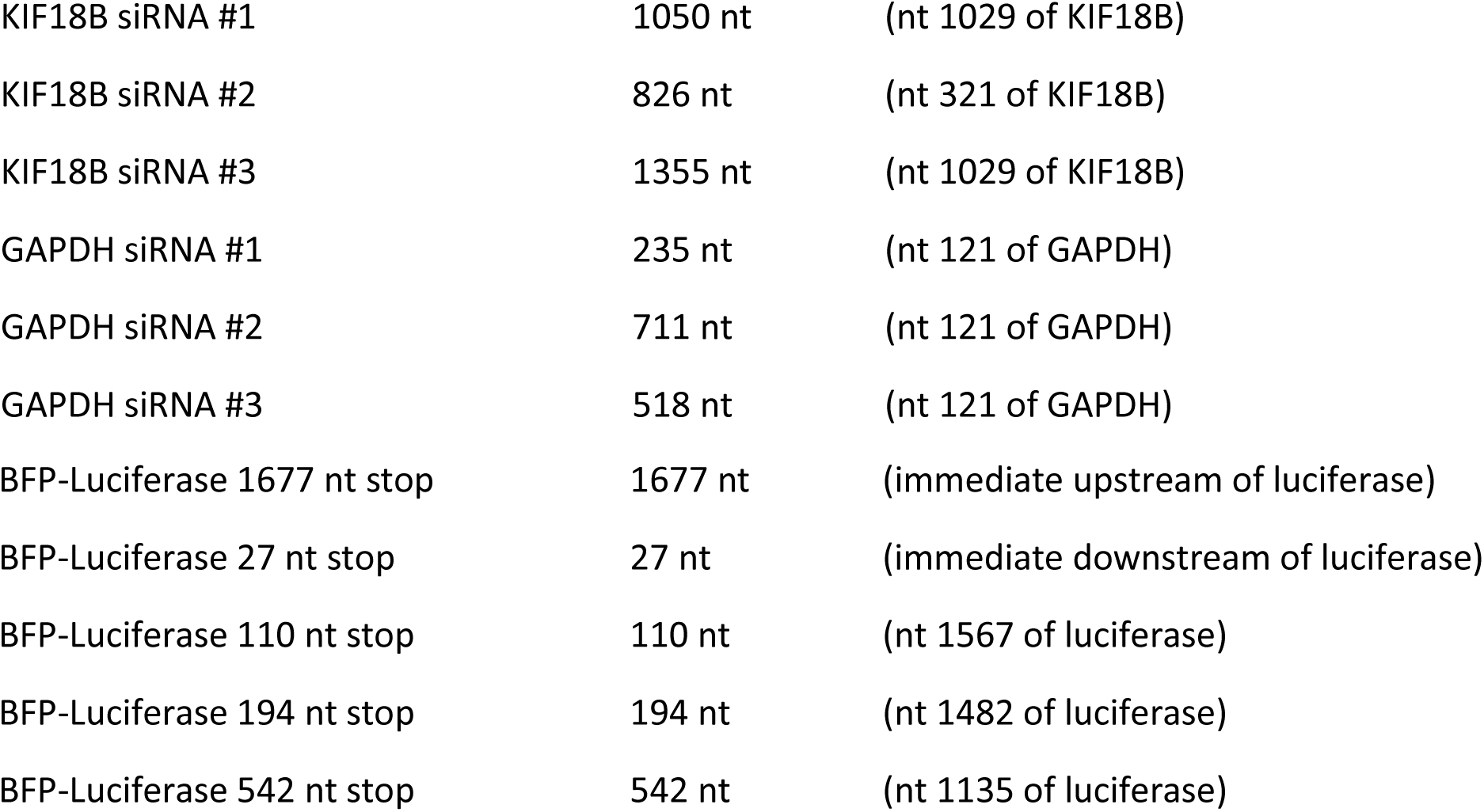

Mutant reporters, that were designed to reduce mRNA structures using the mfold web server (mfold mutants) (Zuker, 2003) and Δ4-mer mutant reporters, were ordered from Genewiz. To design the mfold mutants we examined the 32 most stably folded structures that were generated by the mfold web server in BFP-GAPDH in which the GAPDH siRNA3 binding site was part of an RNA duplex. To remove possible structures caused by base pairing between the siRNA binding site and the surrounding sequences, we introduced single nucleotide substitutions in the sequences predicted to base pair with the siRNA binding site. A total of 19 point mutations were required to remove all possible interactions between the siRNA binding site and the surrounding sequences that were present in the 32 most stably folded structures predicted by mfold. When designing the mfold mutant reporter, the GC content was kept within similar range as the parental reporter. 10/19 mutations were made in the BFP sequence, and 9/19 mutations were made in the GAPDH sequence, 7 of which were upstream and 2 of which were downstream of the siRNA binding site. In 11/19 mutations, altering the nucleotide sequence changed the encoded amino acid. To assess cleavage rates in the absence of ribosomes translating the siRNA binding site, a stop codon was introduced 518 nt upstream of the siRNA binding site. 9 mutations (all in the GAPDH sequence) were downstream of this stop codon.

To design the Δ4-mer mutants in BFP-GAPDH and KIF18B, stretches of 4 consecutive nucleotides (4-mer) which had complete sequence complementarity to a 4-mer sequence in the siRNA binding site (GAPDH siRNA #3 and KIF18B siRNA #1, respectively) were identified. For these 4-mers, a single nucleotide substitution was introduced to disrupt complementarity with the siRNA binding site. In BFP-GAPDH 155 point mutations were introduced, 50 of which were in BFP and 105 in GAPDH. To assess cleavage rates in the absence of ribosomes translating the siRNA binding site, a stop codon was introduced 518 nt upstream of the siRNA binding site. 88 point mutations were downstream of this early-stop codon. If possible, the nucleotide sequence was altered without changing the amino acid sequence. However, in 3/155 mutations the altered nucleotide sequence resulted in a change in the encoded amino acid. Note that for GAPDH siRNA #3 sequence, nucleotides 6-9 and nucleotides 15-18 are fully complementary. This complementarity was not disrupted, as this would change the GAPDH siRNA #3 sequence. In addition, due to a design error 1/20 4-mers, which has complementarity to nt 7-10 of the siRNA binding site, was not disrupted in this reporter, resulting in the presence of seven positions in the construct where 4 consecutive nucleotides are complementary to the GAPDH siRNA #3 sequence. To design the four 25% Δ4-mer mutants in BFP-GAPDH, 25% of the point mutations were made, resulting in a total of 39 point mutations compared to the WT BFP-GAPDH reporter. In order to make four reporters with a non-overlapping set of point mutations, different point mutations were introduced in each reporter. In the 25% Δ4-mer reporter #1, point mutations 1, 5, 9 etc. were made, in reporter #2, point mutations 2, 6, 10 etc. were made, in reporter #3, point mutations 3, 7, 11 etc. were made, and in reporter #4, point mutations 4, 8, 12 etc. were made. In total 5 point mutations were shared between the mfold BFP-GAPDH reporter and the Δ4-mer BFP-GAPDH reporter. In KIF18B 211 point mutations were introduced. To assess cleavage rates in the absence of ribosomes translating the siRNA binding site, a stop codon was introduced 1050 nucleotides upstream of the siRNA binding site. Here, 118 point mutations were downstream of this early-stop codon. In 6/211 mutations, altering the nucleotide sequence resulted in a change in the encoded amino acid.

To create the KIF18B reporter in which the binding site for KIF18B siRNA #1 was disrupted (siRNA-resistant reporter), we introduced 5 mutations in the siRNA binding site. The sequence of the siRNA #1 binding site in this reporter is GCgACtGTgATaAAAAGtC compared to GCCACAGTCATCAAAAGCC in the KIF18B reporter.

#### Plasmid transfections for stable integration

Cells were plated one day prior to transfection in a 6 cm dish (Greiner Bio-one). A transfection mix, containing 100 µl OptiMEM (Sigma-Aldrich), 2 µl FUGENE 6 (Promega), and ∼1 µg of DNA, was added to the cells in a total volume of 1 ml cell culture medium per dish. Selection for stable integration was initiated 24h after transfection, using 0.4 mg/ml Zeocin (Invitrogen), and continued for at least 10 days. To generate monoclonal cell lines, single cells from the polyclonal cell line were sorted into 96-well plates (Greiner Bio-one) by FACS, and grown for 14 days. Individual clones were inspected by microscopy and clones in which a high percentage of cells expressed the transgene were selected for further use. For generating stable monoclonal cell lines expressing reporter mRNA, clones were additionally screened for the number of mRNAs expressed per cell. Clones expressing ∼10-50 mRNAs per cell were selected.

#### siRNA transfections

The complete list and sequence of all siRNAs used in this study is provided in Document S1. siRNAs were custom designed and ordered from Dharmacon, except KIF18B siRNA #1 (AM16708, 251223; ThermoFisher) and GAPDH siRNA #1 (4390849, ThermoFisher). siRNAs were reverse-transfected at a final concentration of 10 nM (unless stated otherwise) using RNAiMAX (Invitrogen) according to the manufacturer’s guidelines. KIF18B siRNA #3 was transfected at a final concentration of 50 nM, as it showed weak target repression at 10 nM. For all microscopy experiments, cells were seeded at a confluency of ∼40-50% in 96-well glass-bottom imaging plates (Matriplate, Brooks) in a final volume of 200 µl and imaged 16-24 hr after transfection. For northern blot experiments, cells were seeded at a confluency of ∼40-50% in a 6 cm plate (Greiner Bio-one) in a final volume of 3 ml and harvested 16-24 hr after transfection. For qPCR experiments, cells were seeded in a 24-well plate (Greiner Bio-one) and harvested 16-24 hr after transfection.

#### Lentivirus production and infection

For lentivirus production, HEK293T cells were plated in a 6-well plate (Greiner Bio-one) at 30% confluency, and transfected 24 hr after plating with a mixture of 50 µl OptiMEM (Sigma-Aldrich), 10 µl Polyethylenimine (PEI) (Polysciences Inc) (1mg/ml), 0.4 µg pMD2.g, 0.6 µg psPAX2, and 1 µg of lentiviral vector. The medium was replaced with 2 ml fresh culture medium 24 hr after transfection, and 72 hr after transfection, viral supernatant was collected. For lentiviral infections, cells were seeded in a 6-well plate (Greiner Bio-one) at 70% confluency. Viral supernatant was added to the cells along with Polybrene (10 µg/ml) (Santa Cruz Biotechnology Inc) and the cells were spun at 2000 rpm for 90 min at 37°C (spin infection). After the spin infection, the culture medium was replaced with fresh medium, and cells were incubated for at least 48 hr before further analysis.

#### CRISPRi-mediated knockdown of endogenous AGO2

To knockdown endogenous AGO2 we made use of CRISPRi, since we reasoned that siRNA-mediated approaches would not be very efficient as they rely on the presence of AGO2. For CRISPRi-mediated knockdown of AGO2 we expressed dCAS9-BFP-KRAB in cells stably expressing TetR, scFv-sfGFP, PCP-Halo-CAAX and 24xGCN4-KIF18B-24xPP7. The 30% highest BFP expressing cells were isolated by FACS sorting for further use. A sgRNA targeting AGO2 (sequence: GCGCGTCGGGTAAACCTGTT) was expressed in cells together with BFP through lentiviral transduction. The BFP signal associated with the sgRNA was much higher than the BFP associated with dCAS9-BFP-KRAB, and thus sgRNA positive cells could be identified in dCAS9-BFP-KRAB-expressing cells. qPCR and imaging were performed 4 to 5 days after infection with the sgRNA. In experiments where cleavage was measured after AGO2-knockdown in combination with expression of an exogenous AGO2 rescue construct (insensitive to the sgRNAs targeting endogenous AGO2), cells were infected with an AGO2 expression construct 10 to 11 days before imaging. As AGO2 rescue construct, we used pLJM1-FH-AGO2-WT, which was a gift from Joshua Mendell (Addgene plasmid #91978; http://n2t.net/addgene:91978) (Golden et al., 2017). Cells expressing exogenous AGO2 were selected with puromycin (2 µg/ml) (ThermoFisher). Infection with exogenous AGO2 was followed by a second infection 4 to 5 days before imaging with the sgRNA targeting AGO2 to knockdown endogenous AGO2.

### smFISH

Single-molecule Fluorescence In Situ Hybridization (smFISH) was performed as described previously (Lyubimova et al., 2013; Ozata et al., 2019; Raj et al., 2008). Five oligonucleotide probes against the PP7-array and 48 probes against the SunTag-array were designed, using the website www.biosearchtech.com (the complete list and sequence of smFISH probes used in this study is provided in Document S1). Probes were synthesized with a 3’ amine modification. Probes were then coupled to either a Cy5 or an Alexa 594 fluorescent dye (Cy5 succinimidyl ester (GE Healthcare) or Alexa Fluor 594 Fluorcarboxylic acid succinimidyl ester (Molecular probes/Invitrogen), respectively) as described previously (Lyubimova et al., 2013), and HPLC purified (ELLA Biotech GmbH). Purified probes were dissolved in 50 µl TE and used at a final dilution of 1:2000. For hybridization, cells were plated in a 96-wells glass bottom dish (Matriplate, Brooks) 16-24 hr before fixation. Doxycycline (1 µg/ml) (Sigma-Aldrich) was added 40 to 90 min before fixation (as indicated). Cells were fixed in PBS with 4% paraformaldehyde (Electron Microscopy Science) for 15 minutes at Room Temperature (RT), washed twice with PBS and incubated for 30 min in 100% ethanol at 4°C. After fixation, cells were washed twice in hybridization buffer with 10% formamide (ThermoFisher) at RT, followed by overnight incubation with the probes in hybridization buffer at 37°C. Following overnight incubation, samples were washed 3x for 1 hour in wash buffer at 37°C. DAPI (Sigma-Aldrich) was added to the final wash step in order to stain the nuclei. Shortly before imaging, samples were placed in anti-bleach buffer (Lyubimova et al., 2013; Raj et al., 2008) to reduce fluorescence bleaching.

### *In vitro* AGO2 binding assay

#### Expression and purification of TMR-HaloTag-AGO2-siRNA

His6-Flag-TEV-Halo-tagged human AGO2 protein was expressed in Sf9 cells using a baculovirus system (Invitrogen). 750 ml of Sf9 cells at 1.7 × 10^6^ cells/ml were infected for 60 hours at 27°C. Infected cells were harvested by centrifugation and resuspended in 30 ml of Lysis Buffer (50 mM NaH2PO4, pH 8, 300 mM NaCl, 5% glycerol, 0.5 mM TCEP). Resuspended cells were lysed by passing twice through an M-110P lab homogenizer (Microfluidics). The resulting total cell lysate was clarified by centrifugation (30,000 × *g* for 25 min) and the soluble fraction was applied to 1.5 ml (packed) Ni-NTA resin (Qiagen) and incubated at 4°C for 1.5 h in 50 ml conical tubes. Resin was pelleted by brief centrifugation and the supernatant solution was discarded. The resin was washed with 50 ml ice cold Nickel Wash Buffer (300 mM NaCl, 15 mM imidazole, 0.5 mM TCEP, 50 mM Tris, pH 8). Centrifugation/wash steps were repeated a total of three times. Co-purifying cellular RNAs were degraded by incubating with 100 units of micrococcal nuclease (Clontech) on-resin in ∼15 ml of Nickel Wash Buffer supplemented with 5 mM CaCl2 at room temperature for 45 minutes. The nuclease-treated resin was washed three times again with Nickel Wash Buffer and then eluted in four column volumes of Nickel Elution Buffer (300 mM NaCl, 300 mM imidazole, 0.5 mM TCEP, 50 mM Tris, pH 8). Eluted AGO2 was incubated with a synthetic siRNA and 150 µg of TEV protease during an overnight dialysis against 1–2 liters of Dialysis Buffer (300 mM NaCl, 0.5 mM TCEP, 50 mM Tris, pH 8) at 4°C. The sequence of the siRNA is provided in Document S1. Please note that the first nucleotide is a U instead of a G (as in the original KIF18B siRNA #1 sequence) to improve siRNA loading in AGO2, which does not affect AGO2-target binding (Frank et al., 2010). AGO2 molecules loaded with the siRNA were isolated using an immobilized capture oligo nucleotide with complementarity to the siRNA, and then eluted by adding competitor DNA with more extensive complementarity to the capture oligo via the Arpon method (Flores-Jasso et al., 2013). Sequences of the capture oligonucleotide and competitor DNA are provided in Document S1. Loaded AGO2 proteins were further purified by size exclusion chromatography using a Superdex Increase 10/300 column (GE Healthcare Life Sciences) equilibrated in 1 M NaCl, 50 mM Tris pH 8, 0.5 mM TCEP. Purified Halo-AGO2-siRNA complex was incubated with Halo-TMR ligand (Promega) and dialyzed against 2 L of 1xPBS (137 mM NaCl, KCl 2.7 mM, 10 mM Na2HPO4, 1.8 mM KH2PO4), concentrated to ∼2 mg/ml, aliquoted, flash frozen with liquid N2, and stored at −80°C.

#### Target RNA synthesis and purification

A short RNA oligonucleotide (sequence provided in Document S1) was ordered from IBA Lifesciences, labeled with a Cy5 dye (Sigma-Aldrich), as described previously (Joo and Ha, 2012), and purified using ethanol precipitation. The labeled oligonucleotide was subsequently ligated to a U30-mer with biotin using T4 RNA ligase II (NEB) and a DNA splint (sequence provided in Document S1).

Full length mRNA targets (KIF18B sequence or KIF18B sequence with a mutated siRNA target site) were *in vitro* transcribed using the HiScribe™ T7 High Yield RNA Synthesis Kit (NEB), and purified using phenol-chloroform extraction and ethanol precipitation. The complete sequence of the full length mRNA targets is provided in Document S1. The full length mRNA targets were ligated to a 22 nt Cy5 labeled and biotinylated RNA oligonucleotide using a 40 nt DNA strand as a splint. The sequences of the oligonucleotide and DNA splint are provided in Document S1. After ligation with T4 RNA ligase II (NEB), the ligated constructs were purified from an agarose gel using a Zymo Gel RNA recovery kit (Baseclear).

#### Sample preparation

Quartz slides were prepared as described previously (Chandradoss et al., 2014). Briefly, quartz slides and coverslips were treated with KOH after which slides were treated with piranha followed with (3-Aminopropyl)triethoxysilane (*APTES*) (Sigma-Aldrich). Slides were subsequently PEGylated with mPEG-SVA (MW 5.000) (Laysan) under a humid atmosphere overnight. Before experiments, an additional round of PEGylation took place with (MS)PEG-4 (ThermoFisher). Quartz slides and coverslips were assembled with double-sided scotch tape after which the chambers were sealed with epoxy glue.

For the single molecule imaging experiments, slides were incubated with T50 and 1% Tween-20 for > 10 min to further passivate the chambers (Pan et al., 2015). Chambers were subsequently rinsed with T50 and streptavidin (0.1 mg/ml) (ThermoFisher) was introduced inside the chamber for 1 min and rinsed out by T50. The RNA sample was then introduced inside the chamber at a concentration of 100 pM. After 1 minute incubation, unbound RNAs were flushed out with T50. Tubing was attached to the outlet of the microfluidic chambers through epoxy glue and an injection needle was attached to the other side of the tubing. Imaging buffer (150 mM NaCl, 2 mM MgCl2, 50 mM Tris (pH 8.0), 1 mM Trolox, 0.8% glucose, 0.1 mg/ml glucose oxidase (Sigma-Aldrich), and 17 μg/ml catalase (ThermoFisher)) was then introduced inside the chamber.

### Microscopy

All in vivo imaging experiments were performed using a Nikon TI inverted microscope with perfect focus system equipped with a Yokagawa CSU-X1 spinning disc, a 100x 1.49 NA objective and an iXon Ultra 897 EM-CCD camera (Andor) using Micro-Manager software (Edelstein et al., 2010) or NIS elements software (Nikon). All live-cell imaging experiments were performed at 37°C, while smFISH experiments were imaged at RT.

#### Live-cell imaging of mRNA cleavage

15 to 30 minutes before imaging, cell culture medium was replaced with pre-warmed CO2-independent Leibovitz’s-15 medium (Gibco) containing 5% fetal bovine serum (Sigma-Aldrich) and 1% penicillin/streptomycin (Gibco). Transcription of the reporters was induced by addition of doxycycline (1 µg/ml) (Sigma-Aldrich) to the cell culture medium. During the experiments, cells were maintained at a constant temperature of 37°C. Cells were selected for imaging based on the levels of mature protein (an indication of reporter expression) and a small number of mRNAs at the start of imaging (Ruijtenberg et al., 2018). For CRISPRi experiments, cells were additionally selected based on the presence of BFP. Camera exposure times of 500 ms were used for both GFP (488 laser) and mCherry (561 laser), and images were acquired every 30 s for 45 minutes, unless stated otherwise. Since mRNAs are tethered to the plasma membrane, we focused the objective slightly above the plasma membrane to focus on both mRNAs and translation sites, and single Z-plane images were acquired.

#### Imaging smFISH samples

Images for all 3 colors (DAPI, Cy5 and Alexa 594) were acquired with a camera exposure time of 50 ms. Z-stacks were acquired for all 3 colors with an inter-slice distance of 0.5 µm each.

#### In vitro imaging of AGO2-target interactions

*In vitro* imaging experiments were performed on a custom built inverted microscope (IX73, Olympus) using prism-based total internal reflection. The Halo-TMR was excited with a 532nm diode laser (Compass 215M/50mW, Coherent), and Cy5 was excited with a 637 nm diode laser (OBIS 637 nm LX 140 mW). A 60x water immersion objective (UPLSAPO60XW, Olympus) was used for imaging. A 532 nm notch filter (NF03-532E-25, Semrock) and a 633 nm notch filter (NF03-633E-25, Semrock) were used to block the scattered light. A dichroic mirror (635 dcxr, Chroma) separates the fluorescence signal into separate channels, and the light is projected onto an EM-CCD camera (iXon Ultra, DU-897U-CS0-#BV, Andor Technology). The *in vitro* experiments were either performed at RT (20°C) or at 37°C through the use of a custom built heating elements and custom written Labview code.

For the co-localization experiments, a short movie was taken with the 637 nm laser (5 mW) as a reference for the position of the RNAs of interest (referred to as reference movie), since the RNA molecules are labeled with Cy5. Subsequently, both green (25 mW) and red (5 mW) lasers were turned on, and movies were taken for 3500 frames at an exposure time of 0.1 s (referred to as measurement movies). After 200 frames, 1 nM Halo-TMR AGO2 complexes together with imaging buffer (see section ‘*Sample preparation’*) were introduced in the microfluidic chamber.

### Northern blot

Northern blots were performed using NorthernMax-Gly kit from ThermoFisher according to the protocol supplied by the manufacturer. In short, cells stably expressing the TetR, scFv-sfGFP, PCP-Halo-CAAX, and 24xGCN4-KIF18B-24xPP7 or 24xGCN4-KIF18B-early-stop-24xPP7 were incubated for 90 min with doxycycline (1 µg/ml) (Sigma-Aldrich) to induce expression of the mRNA reporter and RNA was extracted using TRIsure (Bioline). RNA mixed 1:1 with Glyoxal Load Dye was incubated for 30 min at 50°C to denature RNA before loading 10 µg of RNA onto a 0.8% agarose gel. After running the gel, rRNA (18S and 28S) bands were visualized using UV and signal intensities were quantified to ensure that RNA samples were loaded equally and RNA was intact. RNA transfer from the agarose gel to a positively charged nylon membrane was performed for 2 hours at RT, followed by RNA crosslinking to the membrane using UV light (120mJ/cm^2^ at 254 nm) for 1 min. After prehybridization at 68°C for 1 hour, the membrane was incubated with a DIG-labeled RNA probe targeting the BGH sequence present in the 3’ UTR of the mRNA reporter (the complete sequence of the probe is provided in Document S1), and hybridization was performed overnight at 68°C. The membrane was washed 3x, and incubated with the anti-DIG antibody-AP (Sigma-Aldrich) for 16 to 40 hours at 4°C. The membrane was washed 9x (6x in PBS-T and 3x in AP buffer) and incubated with a few drops of CDP-star for 5 minutes at RT. The film was exposed and developed for 2 to 10 minutes, using an Amersham Imager 600 (GE).

### Quantitative RT-PCR (qPCR)

The complete list and sequence of primers for RT-PCR used in this study is provided in Document S1. To determine the siRNA knockdown efficiency of endogenous KIF18B by qPCR, siRNA treated cells were harvested 24 hr after transfection and RNA was isolated. To measure knockdown levels of endogenous AGO2 by CRIPSRi, cells expressing the dCAS9-BFP-KRAB were infected with sgRNAs targeting AGO2 and harvested 4-5 days later to isolate RNA. RNA was isolated using TRIsure (Bio-line), according to manufacturer’s guidelines. Next, cDNA was generated using Bioscript reverse transcriptase (Bioline) and random hexamer primers. qPCRs were performed using SYBR-Green Supermix (Bio-Rad) on a Bio-Rad Real-time PCR machines (CFX Connect Real-Time PCR Detection System). RNA levels were normalized to the levels of GAPDH mRNA.

### Quantifying translation dynamics

#### Number of ribosomes per mRNA molecule

We determined the number of ribosomes per mRNA molecule (ribosome occupancy) for the uORF-KIF18B and KIF18B reporters. First, we quantified the ribosome occupancy on the uORF-KIF18B reporter. Next, we determined the fold difference in ribosome occupancy between the uORF-KIF18B and KIF18B reporter and used the ribosome occupancy on the uORF-KIF18B reporter to determine the ribosome occupancy on the KIF18B reporter. To determine the ribosome occupancy on a uORF-KIF18B reporter mRNA, we divided the GFP intensity of a translation site by the average intensity of a single mature protein (as the intensity of a single mature protein is equal to the fluorescence intensity associated with a single ribosome that translated the entire SunTag). Next, we multiplied the normalized GFP intensity of a translation site with a correction factor of 1.32 (Yan et al., 2016) to account for those ribosomes that have not completely translated the SunTag yet and will thus have a lower intensity compared to a single mature protein. To determine the GFP fluorescence intensity of single translation sites and single mature proteins, cells containing the uORF-KIF18B reporter were imaged using very short (20 ms) exposure time and maximal laser power (instead of 500 ms exposure time with low laser power, which was used for the majority of experiments), since at long exposure times single mature proteins cannot be visualized due to motion blurring. Next, the GFP fluorescence intensity of single mature proteins and translation sites was measured in FIJI in a circular region of interest (ROI) with a width and height of 6 pixels (0.81 μm). For each measurement, local background fluorescence intensity was measured using a second ROI with the same dimensions at a location directly next to the spot of interest. Finally, the mean background fluorescence intensity was subtracted from the mean spot intensity to obtain the fluorescence intensity of single mature proteins and translation sites.

To obtain the fold difference in ribosome occupancy between the uORF-KIF18B and KIF18B reporters, we divided the average GFP fluorescence intensity of translation sites of the KIF18B reporter by the average GFP fluorescence intensity of translation sites of the uORF-KIF18B reporter. To determine the GFP fluorescence intensity of single translation sites, cells containing either the uORF-KIF18B reporter or the KIF18B reporter were imaged using 500 ms exposure time and low laser power (as we did not image single mature proteins here, the standard imaging conditions could be used). After this, the GFP fluorescence intensity of single translation sites was measured in FIJI in a circular ROI with a width and height of 10 pixels (1.35 μm). For each measurement, local background fluorescence intensity was measured using a second ROI with the same dimensions at a location directly next to the spot of interest. Next, the mean background fluorescence intensity was subtracted from the mean spot intensity to obtain the fluorescence intensity of single translation sites. Finally, the average GFP fluorescence intensity of translation sites of the KIF18B reporter was divided by the average GFP fluorescence intensity of translation sites of the uORF-KIF18B reporter and the previously determined ribosome occupancy for the uORF-KIF18B reporter was used to calculate the average ribosome occupancy for the KIF18B reporter.

#### Arrival time of the first ribosome at the siRNA binding site

For many reporter-siRNA pairs we observed that the probability that an mRNA is cleaved was not constant over time. Rather, we found a peak in the cleavage probability which occurred several minutes after the start of translation (represented by a strong decrease in the fraction of uncleaved mRNAs during this time window). This led us to hypothesize that the increased probability for mRNA cleavage may coincide with the time at which the first ribosome reaches the siRNA target site. To test this hypothesis, we wanted to precisely determine the time at which the first ribosome arrives at the siRNA target site for different reporter mRNAs. The time at which the first ribosome arrives at the siRNA target site could be variable between mRNAs as different ribosomes might move at different speeds (Yan et al., 2016). Therefore, we wanted to determine not only the average time at which the first ribosome arrives at the siRNA target site but also the distribution of the arrival times of the first ribosome (this will be referred to as the *first ribosome arrival time distribution*). To determine the *first ribosome arrival time distribution* at the siRNA target site, we first determined the arrival time distribution at the stop codon and used these values to extrapolate the *ribosome arrival time distribution* at the siRNA target site (which is located upstream of the stop codon). We assumed that translation elongation speed is constant along the mRNA, and thus that the first ribosome arrival time distribution at the stop codon can be extrapolated to the siRNA target site (which is located upstream of the stop codon) based on the difference in length of the mRNA sequence that is translated.

We calculated the *first ribosome arrival time distribution* at the stop codon by measuring the fluorescence intensity build-up over time of newly translated mRNAs. The fluorescence intensity build-up on a single mRNA can be divided into two phases, which we will refer to as the *increasing* phase and the *plateau* phase, respectively. In the first phase, the fluorescence intensity increases, because an increasing number of ribosomes translates the mRNA and synthesizes SunTag peptides that are rapidly fluorescently labeled by the SunTag antibody. The fluorescence intensity of the translation site increases until the first ribosome reaches the stop codon, and remains constant afterwards (which is the start of the *plateau* phase). The fluorescence intensity remains constant after the first ribosome reaches the stop codon, because at that point translation of the mRNA is in steady state: new ribosomes will initiate translation (which will cause an increase in fluorescence intensity), while at the same time ribosomes will terminate translation when they reach the stop codon (which will lead to a decrease in fluorescence intensity as the nascent polypeptide chain with the SunTag peptides is released from the ribosome and diffuses away from the mRNA). Therefore, on average, a constant number of ribosomes will translate the mRNA. Since the transition between the two phases occurs when the first ribosome reaches the stop codon, the time of the transition point reports on the arrival time of the first ribosome at the stop codon. Thus, to determine the distribution of arrival times of the first ribosome at the stop codon, we have to (1) measure fluorescence intensity build-up on single mRNAs and (2) determine for each mRNA the transition time point between the *increasing* and *plateau* phases.

To measure the fluorescence intensities over time on single mRNAs, we used a Matlab-based software package called ‘TransTrack’ that we have recently developed (Boersma et al., 2019). Using TransTrack, we tracked single mRNAs from the start of translation, measured the GFP fluorescence intensity of each translation site and performed local background subtraction. All traces were manually curated and incorrect traces were discarded. In order to determine the transition time point between the *increasing* phase and *plateau* phase, we first determined the slope of each fluorescence intensity trace at all time points by calculating the derivative of the intensity trace. The slope of the fluorescence intensity trace also contains two phases that represent the *increasing* and *plateau* phase: (1) a phase in which the derivative is positive (which represents *the increasing* phase); (2) a phase in which the derivative is around zero (which represents the *plateau* phase).

To find the most likely transition time point between the *increasing* phase and *plateau* phase, we calculated a probability score for each hypothetical transition time point and selected the transition time point with the highest probability. To calculate a probability score, we assessed the probability of observing the derivative intensity trace for each hypothetical transition time point: we expect to observe only positive derivative values before the hypothetical transition time point (as this is during the *increasing* phase), while we expect to observe both positive and negative derivative values after the hypothetical transition time point (as this is during the *plateau* phase). To calculate the probability score we first determined an expected distribution of the derivative during the *increasing* phase and an expected distribution of the derivative during the *plateau* phase. To obtain an expected *increasing* phase slope distribution, we only took the slopes from all traces at early time points (< 3.5 minutes after the start of translation) to ensure that only values were used that represented mRNAs for which the first ribosome had not yet reached the stop codon. Next, we obtained an expected *plateau* phase slope distribution by only taking the slopes from all traces at late time points, to ensure that on all mRNAs the first ribosome has already reached the stop codon. Next, we fit the *increasing* phase slope and the *plateau* phase slope distributions to a normal distribution with fitting parameters *μ* and *σ*, which resulted in good fits.

Next, we used both the *increasing* and *plateau* phase slope distributions to calculate the probability score for the slopes observed before the hypothetical transition time point (using the expected distribution for the *increasing* phase) and the slopes observed after the hypothetical transition time point (using the expected distribution for the *plateau* phase). A total probability score was calculated by adding the probability scores before and after the hypothetical transition time point together. The total probability score was iteratively calculated for all hypothetical transition time points along the fluorescence intensity trace and the transition time point with the highest probability was selected. Finally, a threshold probability score was set to exclude traces that did not result in a good fit (10-20% of all traces). In this way we determined for all GFP fluorescence intensity traces the most likely transition point between the *increasing* phase and *plateau* phase, which marks the arrival of the first ribosome at the stop codon. From the arrival time of the first ribosome at the stop codon on all mRNAs, we generated the ‘first ribosome arrival time’ distribution and we fit the distribution to a gamma distribution with fitting parameters *k* (shape) and *θ* (scale). We used a gamma distribution as it fit the data well and only contains non-negative values (in contrast to a normal distribution).

The analysis described above was performed for three different reporters: KIF18B, KIF18B-early-stop, and GAPDH. This resulted in three gamma distributions for the arrival time of the first ribosome at the stop codon for the respective reporters. Next, the arrival distribution at the different siRNA target sites was determined by scaling the shape parameter of these distributions linearly to the effective mRNA length while keeping the scale parameter of these distributions the same. The effective mRNA length was calculated by determining the distance between the start codon and either the stop codon or the siRNA target site and subtracting a length of 753 nt, which corrects for the fact that the first ribosome is not detectable until approximately 8 SunTag peptides (which have a length of 753 nt) have been translated. For siRNAs targeting the KIF18B reporter (siRNA #1, #2, and #3) the average ribosome arrival time at the stop codon of both the KIF18B as well as the KIF18B-early-stop was used to extrapolate the ribosome arrival time at the siRNA target sites, an average of both reporters was used to increase the reliability. For siRNAs targeting the GAPDH reporter (siRNA #1, #2, and #3) the average ribosome arrival time at the stop codon of the GAPDH reporter was used to extrapolate to the siRNA target sites. For siRNAs targeting the TPI1 reporter (siRNA #1, and #2) the average ribosome arrival time at the stop codon of all three reporters (KIF18B, KIF18B-early-stop, and GAPDH) was used to extrapolate to the siRNA target sites, and the average of the three reporters was used to increase the reliability.

#### Effects of siRNAs on translation efficiency

To determine the effects of siRNAs on translation dynamics, we analyzed the GFP fluorescence intensity on the KIF18B reporter in the presence or absence of KIF18B siRNA #1 (figure S1I and S1J). The GFP fluorescence intensity on newly transcribed mRNAs increases from the start of translation as multiple ribosomes sequentially initiate translation on an mRNA molecule and sequentially synthesize all 24 SunTag peptides that are rapidly fluorescently labeled with the SunTag antibody. The slope of the increasing fluorescence intensity of single translation sites depends both on the translation initiation and elongation rate. The higher the translation initiation rate, the more ribosomes will simultaneously synthesize SunTag peptides, and thus the faster the increase in the GFP fluorescence will be. Similarly, the higher the translation elongation rate, the faster ribosomes synthesize SunTag peptides. Therefore, by comparing the slope of the fluorescence intensity increase in the presence and absence of siRNA, we can assess whether the siRNA affects the translation initiation and/or elongation rate. To obtain the slope of the fluorescence intensity increase, we tracked single mRNAs in the presence and absence of siRNA, measured the GFP fluorescence intensity, and performed local background subtraction using TransTrack (Boersma et al., 2019). All traces were manually curated. Next, the slope of the fluorescence intensity increase was calculated by determining the derivative of the GFP fluorescence intensity traces.

To compare the slope of the fluorescence intensity in the presence and absence of siRNA two additional considerations have to be taken in account. First, the fluorescence intensity increase on a single mRNA is not constant over time, but rather contains three phases with distinct slopes. At the start of translation the slope of the fluorescence intensity increases, because at first only a single ribosome is synthesizing SunTag peptides. However, at later time points multiple ribosomes are simultaneously synthesizing SunTag peptides, resulting in a larger increase of the fluorescence intensity per time unit. The maximal slope of the fluorescence intensity is achieved when the first ribosome reaches the end of the SunTag array, because from this time point on the maximal number of ribosomes is simultaneously synthesizing SunTag peptides. Thus, the first phase is defined as the time from translation initiation by the first ribosome, until the time when the first ribosome reaches the end of the SunTag array. In the second phase, the fluorescence intensity of the translation site will increase at a constant rate (i.e. the maximal rate) until the first ribosome reaches the stop codon. The third phase starts once the first ribosome has reached the stop codon, because from that time point on the increase in the fluorescence intensity by newly initiating ribosomes is offset by a decrease in the fluorescence intensity due to terminating ribosomes. This results in a net slope of zero. To compare the slope of the fluorescence intensities in the presence and absence of siRNA, we focused on the second phase where the fluorescence intensity increases linearly with a maximum slope. Since the slope during the second phase remains constant, this allowed us to average multiple time points to create a larger dataset,. A time window between 1.5 and 4 minutes after the start of translation was selected based on visual inspection of the fluorescence intensity graph (as during this time window a linear increase was observed).

The second consideration to take into account when comparing the slopes of the fluorescence intensity traces in the presence and absence of siRNA, is that in the presence of siRNA it is unknown at which timepoint mRNAs have an AGO2/siRNA complex bound, as the AGO2/siRNA complex is not fluorescently labeled itself. We would not expect an effect on the translation dynamics for mRNAs that don’t have an AGO2/siRNA complex bound. Therefore, we only examined translation efficiency immediately before cleavage was observed, since AGO2/siRNA complex must have been bound to the mRNA at the moment of cleavage. To determine the translation efficiency immediately before cleavage, we only calculated the slope of the GFP intensity curve between the two time points preceding the moment of cleavage, and grouped mRNAs together that were cleaved in the time window between 1.5 and 4 minutes after the start of translation (see section *’determining the moment of mRNA cleavage‘* for a description on the quantification of the cleavage time) to average the slope of multiple mRNA molecules. The average slope of the fluorescence intensity increase in siRNA treated cells and control cells was compared using a paired two-tailed t-test (Figure S1K).

#### Time between CHX treatment and inhibition of ribosome elongation

To determine how long after addition of CHX ribosomes are completely immobilized, we analyzed GFP intensity build-up on mRNAs on which translation started 1 to 3 minutes before the addition of CHX. In the absence of CHX we expect a constant increase in GFP intensity during this phase (See *‘Arrival time of the first ribosome at the siRNA site’)*. However, when CHX is active, translation elongation will be inhibited, so GFP intensity will no longer increase. To determine the GFP fluorescence intensity of translation sites, the mean GFP intensity of translation sites was measured in FIJI in a circular region of interest (ROI) with a width and height of 9 pixels (1.09 μm). For each spot, local background fluorescence intensity was measured using the same ROI at multiple locations directly next to the spot of interest, and the mean background fluorescence intensity was subtracted from the mean spot intensity. GFP intensity was measured from the first frame of GFP appearance until 3-5 minutes after CHX addition. Intensity traces were aligned at the moment of CHX addition and the moment at which GFP no longer increased over time was determined.

### Analysis of mRNA cleavage

#### Scoring mRNA cleavage

Two types of mRNA cleavage events could be observed in our live-cell imaging assay: (1) cleavage events where GFP and mCherry foci separated completely, which represents mRNAs in which all ribosomes were present on the 5’ cleavage fragment (for example, because the first translating ribosome had not yet passed the cleavage site) and (2) cleavage events where one or more ribosomes were present on the 3’ cleavage fragment upon cleavage, resulting in GFP foci that separated from dual-color GFP/mCherry foci. In the latter case, at least one ribosome must have already passed the cleavage site at the moment of cleavage.

For mRNAs that did not show cleavage, we tracked the total time that the translation signal associated with the mRNA could be observed. A track was ended either when: (1) translation of the mRNA could no longer be observed, (2) the end of the time lapse movie was reached, (3) when the mRNA moved out of the field of view, (4) when a mRNA would cross paths with a fluorescent lysosome or another mRNA, or (5) when the mRNA detached from the plasma membrane.

#### Determining the moment of mRNA cleavage

To quantify the precise moment of cleavage of reporter mRNA molecules relative to the start of translation, we determined the first time point at which we observed GFP signal for each mRNA, as well as the moment where we observed spatial separation of the GFP and mCherry foci (which was scored as cleavage). If no cleavage was observed for an mRNA during the course of the experiment, we tracked the total time that the translation signal on the mRNA could be observed (see section ‘*scoring mRNA cleavage’*). To ensure that we had observed the first round of translation on each mRNA molecule, we excluded mRNAs that (1) were already present in the field of view at the start of the movie, (2) mRNAs that were already associated with GFP fluorescence when they first appeared in the field of view, or (3) mRNAs that were never associated with GFP fluorescence. We plotted the fraction of uncleaved mRNAs using a Kaplan-Meier curve (which we refer to as a cleavage curve), which takes into account both the track length of the cleaved and uncleaved mRNAs.

#### Distinguishing mRNA cleavage from translation termination

In our assay, spatial separation of GFP and mCherry foci was scored as mRNA cleavage. However, separation of GFP foci from mRNAs (mCherry foci) could also represent translation termination of a ribosome and release of the (GFP-positive) nascent polypeptide. A key difference between these two processes is that for translation termination the GFP foci that are separating from the mRNA represent a single ribosome, while for mRNA cleavage the GFP foci represent the 5’ mRNA cleavage fragment, which generally contains multiple ribosomes and thus is associated with a higher GFP fluorescence intensity. To demonstrate that separation of GFP and mCherry foci represented mRNA cleavage rather than translation termination, we quantified the intensity of the GFP foci that separated from the mCherry foci. We compared the fluorescence intensity of the GFP foci to the intensity of a single mature protein (which is similar to the fluorescence intensity that is associated with a single ribosome at the moment of translation termination) to establish whether GFP foci after cleavage represented a single ribosome (as expected for translation termination) or multiple ribosomes (as expected for mRNA cleavage). To visualize single mature proteins, we imaged with 100% laser power and 20 ms exposure. Next, to determine the fluorescence intensity of GFP foci that separated from mCherry foci and single mature proteins, GFP intensities were measured in FIJI in an ROI with a width and height of 9 pixels (1.09 μm). For each GFP spot (both GFP foci after separation from mCherry foci and single mature proteins), local background fluorescence intensity was measured using a second ROI at multiple locations directly next to the spot of interest, and the mean background fluorescence intensity was subtracted from the mean spot intensity. Mean intensities of single mature proteins were then compared to GFP foci that separated from mCherry foci.

#### Analysis of mRNA cleavage with the EMI1-KIF18B reporter

To analyze mRNA cleavage on mRNAs translated by a single ribosome, we introduced a 5’ UTR sequence of the EMI1 gene in our reporter. This isoform of the 5’ UTR of EMI1 has been shown to act highly repressive on translation initiation, frequently limiting the number of ribosomes per mRNA molecule to one ribosome (Tanenbaum et al., 2015; Yan et al., 2016). To ensure that exclusively translation events with single ribosomes were analyzed, translation events were selected on two criteria: (1) the GFP intensity correlated with the intensity of a single ribosome and (2) the time between GFP appearance and disappearance was <7.5 minutes, as longer events are more likely to represent either translation events by multiple ribosomes or stalled ribosomes. Because translation termination of a single ribosome cannot be readily distinguished from mRNA cleavage, as they both result in the separation of a single ribosome/SunTag array from the mRNA, we used mRNA disappearance (i.e. decay of the 3’ cleavage fragment) as a read-out for mRNA cleavage (Hoek et al., 2019). Therefore, we determined the time from GFP appearance to mRNA disappearance for each mRNA molecule.

#### Quantification of mRNA cleavage rates

From the Kaplan-Meier cleavage curves (referred to as cleavage curves) we quantified the mRNA cleavage rates for different reporters in the presence or absence of ribosomes translating the siRNA binding site (Figures 2D, 5E, 6F-G, S3G, and S4G). To determine the cleavage rate in the absence of ribosomes translating the siRNA binding site (e.g. experiments in which CHX was added to cells, or in which a stop codon was introduced upstream of the AGO2 binding site), we fit the cleavage curve (of each individual replicate experiment) with a single exponential function. A plateau was included in the fit, as we frequently observed that a small fraction of mRNAs was resistant to cleavage (e.g. see Figure 5B and Equation 1).

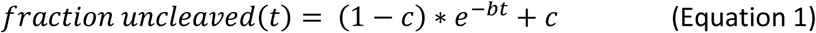

Here, *b* is the cleavage rate, *c* is the plateau level, and *t* is the time in seconds after the first detection of GFP on an mRNA molecule (i.e. the start of translation). We fixed the plateau level for all siRNAs at the plateau level observed for GAPDH siRNA #3 in the GAPDH Δ4-mer reporter (∼15%) (see Figure 5B).

To determine the cleavage rate in the presence of ribosomes translating the siRNA target site, we could not fit the cleavage curve with a single exponential function, because the cleavage curve is more complex. Initially the cleavage curve decreases slowly, followed by a faster decrease from the moment the first ribosome arrives at the siRNA target site, and a final slower decrease of the last 10-20% of mRNAs. Therefore, instead of fitting with a single exponential function, we determined the cleavage rate at individual time points using Equation 2.

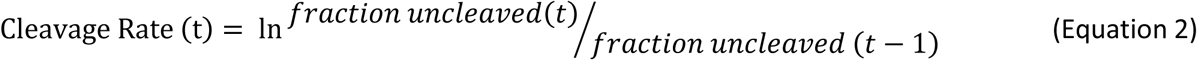

Using Equation 2, the cleavage rate can be computed at each time point and we averaged the cleavage rate over a specific time window (see below) to determine the cleavage rate in the presence of ribosomes translating the siRNA target site. The start of this time window was set at the time point when on 50% of the mRNAs the first ribosome has reached the siRNA target site. This time point was determined for each individual siRNA and reporter based on the ribosome arrival distribution (see section ‘*Arrival time of the first ribosome at the siRNA binding site*’). The end of the abovementioned time window was set at the time point when the fraction of uncleaved mRNAs reached 20% (or at the end of the cleavage curve in case the 20% is never reached), because we observed that in most experiments the last 10-20% of mRNAs is cleaved at a lower rate, for reasons that are currently unclear. Statistical significance of differences between cleavage rates were tested using a two-tailed Student’s t-test.

#### Analysis of ‘late’ cleavage events in CHX-treated cells

We used different datasets in which cleavage was scored after CHX treatment to assess the mRNA refolding rate (Figure 6A, and S5B-D). We found that CHX addition reduces the cleavage rate, likely due to refolding of the mRNA and masking of the target site. However, a delay is expected between CHX addition and the decrease in the cleavage rate, which represents the time needed for mRNA re-folding Thus, the transition time from a fast to a slow cleavage rate reports on the mRNA refolding rate. To assess the mRNA refolding rate, we therefore analyzed the cleavage rates for CHX treated cells in more detail. In our initial analyses all translating mRNAs were included. However, in the current analysis we wanted to only include mRNAs on which the first ribosome had recently translated (and unfolded) the siRNA target site to omit mRNAs on which the first ribosome had not reached the siRNA target site yet (as the target site had not been unmasked on these mRNAs yet), and mRNAs on which the first ribosome has passed the siRNA target site long ago (as these are likely enriched in the small subset of mRNAs that appear resistant to siRNA-mediated cleavage). To determine the time window during which the first ribosome recently translated the siRNA target site we used the first ribosome arrival distribution at the siRNA target site (see section *‘Arrival time of the first ribosome at the siRNA target site’*) and we selected for each reporter the time points during which on 85-90% of the mRNAs the first ribosome had arrived at the siRNA target site. For KIF18B siRNA #1 this included mRNAs on which translation had started between 3.5 and 6.5 minutes before CHX addition. For KIF18B siRNA #2 this included mRNAs on which translation had started between 2.5 and 5.0 minutes before CHX addition. For GAPDH siRNA #1 this included mRNAs on which translation had started between 2.0 and 4.5 minutes before CHX addition. For GAPDH siRNA #3 this included mRNAs on which translation had started between 2.0 and 5.0 minutes before CHX addition. Next, we fit the resulting cleavage curves with a double exponential function to assess the transition time from a fast to a slow cleavage rate. For 1/4 reporters (GAPDH reporter with GAPDH siRNA #3) the data did not fit well with a double exponential function and was omitted from further analysis.

### Analysis of the fraction of mRNAs with ribosomes on the 3’ cleavage fragment

#### Detecting ribosomes on the 3’ cleavage fragment

To distinguish between the *release model* (ribosomes collide into AGO2, stimulating release of cleavage fragments) and the *binding model* (ribosomes unmask the siRNA target site, stimulating the binding of AGO2 to the target site), we quantified the fraction of cleavage events where one or more ribosomes were present on the 3’ cleavage fragment. The *binding* model predicts that one or more ribosomes should be present on most 3’ cleavage fragments, while the *release* model predicts that ribosomes should be mostly absent from the 3’ cleavage fragment. Since the fluorescence intensity of a single ribosome is weak and the signal is only present on the 3’ cleavage fragment for a very short amount of time (as the length between the siRNA target site, where the mRNA is cleaved, and the stop codon is relatively short for most reporters), we modified our assay in two ways to allow efficient detection of ribosomes on the 3’ cleavage fragment.

The first modification was made to the reporters. We made two new reporter constructs where a linker sequence was added between the siRNA binding site and the stop codon to increase the time a ribosome remains on the 3’ cleavage fragment. The linker sequence facilitates the detection of the ribosome on the 3’ cleavage fragment as it takes longer for the ribosome to reach the stop codon after passing the siRNA target site. For the KIF18B reporter, we inserted the sequence of the fluorescent protein mRuby (714 nt) as a linker; for the GAPDH reporter, we inserted the KIF18B sequence (2568 nt) as a linker (the KIF18B sequence could not be used as a linker in the KIF18B reporter as this would duplicate the siRNA target site). Introduction of these linker sequences in the KIF18B and GAPDH reporters did not substantially alter cleavage kinetics (Figure S2K).

The second modification to the assay was made to the image acquisition settings. We acquired images every 15 s instead of every 30 s to more efficiently detect GFP signals only present for a short amount of time on the 3’ cleavage fragment, and we imaged with high laser power to detect weak signals associated with only a single ribosome. With these two modifications we quantified for each mRNA the time of cleavage relative to the start of translation (as described in the section ‘*Determining the moment of mRNA cleavage’*) and scored for each cleavage event if there was a ribosome present on the 3’ cleavage fragment (i.e. whether a GFP signal was associated with the 3’ fragment after cleavage). Finally, we grouped mRNAs together that were cleaved at the same time point relative to the start of translation and calculated for each time point the fraction of cleavage events with a ribosome on the 3’ cleavage fragment.

#### Normalization of the fraction of mRNAs with ribosomes on the 3’ cleavage fragment

To distinguish between the *release* and *binding* models, we performed two processing steps on the fraction of cleavage events with a ribosome on the 3’ cleavage fragment.

First, we selected cleavage events that occurred around the time when the first ribosome reached the siRNA target site. Cleavage events that occurred before the first ribosome reached the siRNA target site (early cleavage events) or cleavage events that occurred when a second or subsequent ribosome reached the siRNA target site (late cleavage events) are not informative to distinguish between abovementioned models, because in both models **none** of the early cleavage events are predicted to have a ribosome present on the 3’ cleavage fragment (as the first ribosome hasn’t reached the siRNA target site yet), while **all** of the late cleavage events will have a ribosome present on the 3’ cleavage fragment (as the first ribosome has already passed the siRNA target site). To select cleavage events that occurred around the time of the first ribosome arrival at the cleavage site, we used the first ribosome arrival distribution (see section *‘Arrival time of the first ribosome at the siRNA target site’*) to select the three time points when the first ribosome was most likely to reach the siRNA target site.

As reporter mRNAs can also be cleaved in the absence of ribosomes translating the siRNA target site (e.g. + CHX experiments), and ribosome-independent cleavage may even occur when a ribosome is close to the cleavage site, it is likely that not all cleavage events that occur in the time period when ribosomes arrive at the cleavage site are ribosome-dependent cleavage events (even in the absence of ribosomes translating the siRNA target site (CHX experiments) cleavage is observed during the same time window). Ribosome-independent cleavage events are not informative to distinguish between the models, because in all models the same fraction of ribosome-independent cleavage events will have a ribosome present on the 3’ cleavage fragment. Therefore, during the second processing step we wanted to correct for ribosome-independent cleavage by calculating the fraction of ribosome-dependent cleavage events for each time point, and assessing which fraction of ribosome-dependent cleavage events contained a ribosome on the 3’ cleavage fragment. To estimate the fraction of ribosome-dependent cleavage events with a ribosome on the 3’ cleavage fragment at each time point relative to the start of translation, we used Equation 3.

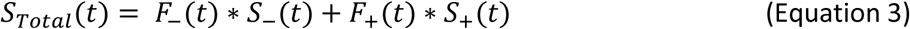

Here, *t* is the time relative to moment of GFP appearance (i.e. the start of translation), *S_Total_* is the fraction of all cleavage events with a ribosome on the 3’ cleavage fragment, *F_−_* is the fraction of ribosome-independent cleavage events, *F*_+_ is the fraction of ribosome-dependent cleavage events, and *S*_−_ the fraction of ribosome-independent cleavage events with a ribosome on the 3’ cleavage fragment. *S*_+_ is the fraction of ribosome-dependent cleavage events with a ribosome on the 3’ cleavage fragment, and we calculated *S*_+_for the three time points when the first ribosome was most likely to reach the siRNA target site. To calculate *S*_+_ with Equation 3, we first determined the other four parameters. As we have experimentally determined *S_Total_* (the fraction of cleavage events with a ribosome on the 3’ cleavage fragment), we only need to calculate *F*_−_, *F*_+_, and *S*_−_. To calculate the fraction of cleavage events that were either ribosome-independent (*F*_−_) or ribosome-dependent (*F*_+_) at each time point, we first calculated the rate of ribosome-independent cleavage using the data sets where cells were treated with CHX (+CHX datasets; as ribosomes are not *translating* the siRNA target site in the presence of CHX). We fit the +CHX datasets with a single exponential function, which was used as the rate of ribosome-independent cleavage. Next, to calculate the fraction of ribosome-dependent cleavage events, we subtracted the exponential function from the cleavage curve obtained in the absence of CHX. Finally, to calculate the fraction of ribosome-independent cleavage events with a ribosome on the 3’ cleavage fragment (*S*_−_), we used the first ribosome arrival distribution at the siRNA target site. We expect that the fraction of ribosome-independent cleavage events with a ribosome on the 3’ cleavage fragment for each time point is directly related to the ribosome arrival distribution (if at a given time point in 30% of the mRNAs the first ribosome has passed the siRNA target site, we would expect 30% of the ribosome-independent cleavage events to have a ribosome on the 3’ cleavage fragment). Using the calculated values of *F*_−_, *F*_+_, and *S*_−_, and Equation 3, we calculated for the three time points (i.e. those with the highest probability of the first ribosome arriving at the cleavage site; see above) the expected fraction of ribosome-independent cleavage events with a ribosome on the 3’ cleavage fragment (*S*_+_). Finally, we calculated a weighted average of *S*_+_ for the three time points by weighting each time point based on the total number of ribosome-dependent cleavage events at each time point.

#### Expectations from release and binding models

To determine whether the experimentally-observed fraction of mRNAs with a ribosome on the 3’ cleavage fragment was most consistent with the *release* or *binding* model, we compared the fraction of ribosome-dependent cleavage events with a ribosome on the 3’ cleavage fragment with the fraction that we would expect with the *release* and *binding* model. In the *binding* model we expect that a ribosome translates over the siRNA target site to make the target site available for AGO2, which will result in mRNA cleavage with a ribosome on the 3’ cleavage fragment. Therefore, most ribosome-dependent cleavage events will lead to a cleavage with a ribosome on the 3’ cleavage fragment (so we would expect close to 100% of the ribosome-dependent cleavage events to have a ribosome on the 3’ cleavage fragment). Note that in some cases the siRNA binding site may interact with an upstream sequence in the mRNA to mask the target site. In these cases, translation of the upstream sequence by the ribosome may be sufficient to unmask the target site. If cleavage occurs very rapidly after unmasking, before the ribosomes reaches the siRNA binding site, then the ribosome will not end up on the 3’ cleavage fragment, even though it stimulated cleavage through the binding model. Thus, the binding model predicts that slightly less than 100% of cleavage events contain a ribosome on the 3’ cleavage fragment.

In the *release* model we expect that a ribosome collides with AGO2 at the siRNA target site and stimulates AGO2 release from the 5’ and 3’ cleavage fragments. If the first ribosome translating an mRNA collides with AGO2 and stimulates release, no ribosomes will be present on the 3’ cleavage fragment. However, it is also possible that the first (few) ribosome(s) translates the cleavage site, after which AGO2 binds and cleaves the mRNA, and that a subsequent ribosome collides with AGO2 to cause fragment release. In this case we do expect a ribosome on the 3’ cleavage fragment. Thus, to determine the expected fraction of ribosome-dependent cleavage events with a ribosome on the 3’ cleavage fragment for the *collision* model, we need to calculate the fraction of ribosome-dependent cleavage events that is caused by the first ribosome and by subsequent ribosomes.

To estimate the fraction of ribosome-dependent cleavage events caused by the first ribosome versus subsequent ribosomes, we need to calculate the fraction of mRNAs on which the first ribosome has passed the cleavage site at the moment of cleavage. For example, if on 30% of the mRNAs the first ribosome has passed the cleavage site, we would expect that 30% of the ribosome-dependent cleavage events are caused by a subsequent ribosome and will have a ribosome on the 3’ cleavage fragment. To calculate the fraction of mRNAs on which the first ribosome has passed the cleavage site, we made a correction to the first ribosome arrival distribution that was determined previously (see section *‘Arrival time of the first ribosome at the siRNA binding site’*). From the first ribosome arrival distribution we know for each time point the mRNA fraction on which the first ribosome has passed the cleavage site. However, we also expect with the *collision* model that many mRNAs are cleaved when the first ribosome reaches the cleavage site. Thus, if we would expect, based on the ribosome arrival distribution, that on 30% of the remaining mRNAs the first ribosome has passed the cleavage site, the real fraction would be much lower as most of these mRNAs have been cleaved when the first ribosome reached the target site. To correct for mRNA cleavage when the first ribosome reaches the siRNA target site, we need to determine the fraction of mRNAs that is cleaved upon arrival of the first ribosome. To determine the fraction of mRNAs that is cleaved, we looked at the three time points when on most mRNAs the first ribosome reaches the cleavage site, and determined the amount of ribosome-dependent cleavage and divided this by the fraction of mRNAs where the first ribosome arrived during the three time points. Next, we used the fraction of mRNAs that is cleaved upon arrival of the first ribosome to correct the first ribosome arrival distribution; for example, if at a given time point on 30% of the mRNAs the first ribosome arrives at the cleavage site, but also 50% of these mRNAs are cleaved because of ribosome-AGO2 collisions, we expect that after this time point only 15% of these mRNAs are remaining. The first ribosome arrival distribution was corrected in this fashion, which resulted in a new distribution that reports for each time point on which fraction of mRNAs the first ribosome has already passed the cleavage site. As only ribosome-dependent cleavage events caused by subsequent ribosomes lead to a cleavage with a ribosome on the 3’ cleavage fragment, we then calculated the expected fraction of ribosome-dependent cleavage events with a ribosome on the 3’ cleavage fragment for the *release* model based on this corrected ribosome arrival distribution.

### Quantification of smFISH experiments

#### Number of mRNAs

To quantify the number of mRNAs based on smFISH, multiple Z slices were made (with an interslice distance of 0.5 µm each) and maximum intensity Z-projections were created). Dependent on whether we wished to quantify the number of mRNAs in the nucleus and cytoplasm (Figure S1B-F, 2SJ), or tethered to the membrane (Figure 2SJ), maximum projections containing different slices were created. For measurements in the nucleus and cytoplasm, maximum projections which included all slices in which the nucleus was present (based on DAPI) were used (to prevent calling cytoplasmic mRNAs nuclear). To quantify the number of mRNAs tethered to the membrane, maximum projections of the two slices containing the bottom membrane were made. Using TransTrack, the nucleus was identified based on DAPI and the number of mRNAs at each location was quantified.

#### Co-localization between SunTag and PP7 probes

To quantify the percentage of co-localization between the SunTag and PP7 smFISH probes, mRNAs were identified using TransTrack based on the SunTag probe signal (Cy5). For each mRNA we manually determined whether the SunTag smFISH signal co-localized with the PP7 signal (Alexa 594).

#### Transcription site intensities

To quantify the transcription site intensities, maximum intensity Z-projections were created (images were taken with an interslice distance of 0.5 µm each). To ensure that all the fluorescence signal of the transcription site was captured, the maximum intensity projections included all slices in which the nucleus was present (based on DAPI signal). To quantify the fluorescence intensity of the transcription site, an ROI was manually drawn around the transcription site, and the integrated fluorescence intensity was measured. For each spot the background fluorescence intensity was measured in the cytoplasm using a second ROI with the same dimensions. The background fluorescence intensity was subtracted from the transcription site fluorescence intensity.

### Quantification of northern blots

Northern blot images were analyzed using ImageQuant TL. The total intensity of each band was measured and background was subtracted using the manual baseline option (i.e. the background intensity was measured manually). To control for loading differences, the RNA gel was analyzed and both the 18S rRNA and 28S rRNA integrated band intensity were measured. An average normalization factor was calculated based on the 18S rRNA and 28S rRNA integrated intensity and the northern blot band intensities were normalized accordingly.

### Quantification of *in vitro* AGO2 on rate

#### Identifying AGO2 binding events

RNA molecules were first localized in the reference movie through custom code written in IDL. Non-specific interactions of AGO2/siRNA complexes with the chamber surface (i.e. interactions that do not show co-localization with the RNA molecules) are ignored. Next, intensity time traces were created in the measurement movie for each RNA molecule (based on the positions of the RNA molecules in the reference movie) and the resulting intensity time traces were further processed in MATLAB (Mathworks) using custom code. To determine the binding rate, we measured the time between introduction of AGO2/siRNA complexes in the sample chamber and the time when stable binding occurred (stable binding is defined by interactions of > 1 s).

#### Calculating the in vitro AGO2 on-rate

The binding time of AGO2 was calculated as the time between AGO2/siRNA complex introduction into the imaging chamber and the first stable binding event for each RNA molecule. For the short RNA oligo target, the majority of molecules was bound by an AGO2 molecule within our time window of 350 s. To calculate the on-rate, we fit the data with Equation 4.

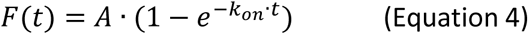

*F* is the fraction of bound molecules, *A* the maximum bound fraction, t the time and *k_on_* the on rate.

For the full length mRNA targets, most molecules were not bound by an AGO2 molecule within our time window of 350 s. Therefore, it was not possible to fit the data with Equation 4 and we instead linearized Equation 4.

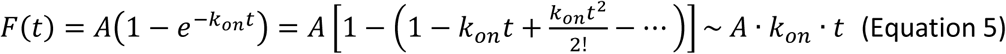

The approximation in Equation 5 is valid as long as the product *k_on_* · *t* is very small. Using Equation 5, we fit the datapoints from the first hundred seconds to determine the value of *A* · *k_on_*. Next, we calculated the on rate (*k_on_*) by dividing with *A*. We took the value of *A* that we had fit with the short oligo target.

### Calculating AGO2 cleavage cycle kinetics

Using the cleavage curves, we wished to estimate the average duration of the AGO2 cleavage cycle *in vivo*. The AGO2 cleavage cycle consists of AGO2/siRNA target binding, mRNA slicing, and fragment release. In the cleavage curves only the moment of fragment release is quantified. To calculate the average duration of the AGO2 cleavage cycle, we have to determine the moment at which the target mRNA becomes accessible for AGO2/siRNA target binding, as target binding can only initiate when the AGO2 target site is unmasked. Target site unmasking can occur independently of ribosomes (e.g. through structural rearrangements) or through ribosomes translating and unfolding the AGO2 target site. For ribosome-dependent cleavage (i.e. unmasking through ribosomes unfolding the AGO2 target site), we can estimate the start of the cleavage cycle (defined here as the moment the target site becomes unmasked) based on the first ribosome arrival time distribution at the AGO2 target site. For ribosome-independent cleavage (i.e. unmasking through structural rearrangements) we cannot estimate the start of the cleavage cycle. Therefore, to estimate the average cleavage cycle duration we focused on ribosome-dependent cleavage events.

To estimate the duration of the AGO2 cleavage cycle, theoretical cleavage curves assuming different durations of the AGO2 cleavage cycle were computed and compared to the experimentally-derived cleavage curves at different siRNA concentrations (10 nM, 0.75 nM, and 0.1 nM). The theoretical cleavage curve is composed of the sum of ribosome-independent cleavage events and ribosome-dependent cleavage events. To calculate the ribosome-independent cleavage curve, we fit the start of the experimentally-derived cleavage curves (first 3.5 minutes after the start of translation) with an exponential function. At these early time points the first ribosome has not reached the AGO2 target site yet, and therefore represent ribosome-independent cleavage. Next, we computed theoretical ribosome-dependent cleavage curves for different durations of the AGO2 cleavage cycle. As we plot the ‘cleavage time’ relative to GFP appearance (i.e. start of translation), the ‘cleavage time’ is the sum of (1) the time it takes for the first ribosome to reach the AGO2 target site (which will unmask the target site and start the cleavage cycle) and (2) the duration of the AGO2 cleavage cycle itself. For the arrival time of the first ribosome at the AGO2 target site we used the first ribosome arrival time distribution (see section *‘arrival time of the first ribosome at the siRNA binding site’*). For the duration of the AGO2 cleavage cycle, we used an exponential distribution with different mean cleavage cycle durations (ranging from 1 s to 30 min). Next, we convoluted the first ribosome arrival time distribution with the different cleavage cycle duration distributions to obtain ribosome-dependent cleavage curves. Finally, we added the ribosome-independent cleavage curve and ribosome-dependent cleavage curves together to obtain theoretical cleavage curves for different durations of the AGO2 cleavage cycle. To calculate the average cleavage cycle duration that best fit with the data (for 10 nM, 0.75 nM, and 0.1 nM), we calculated the sum of squared errors (SSE) between the theoretical cleavage curves (1 s to 30 min duration of the cleavage cycle) and the experimentally-derived cleavage curves (for 10 nM, 0.75 nM, and 0.1 nM).

### Modelling mRNA folding kinetics

#### Simulations of mRNA refolding rate

We found that ribosomes translate and unmask the siRNA target site by unfolding local mRNA structure, facilitating AGO2/siRNA complex binding and mRNA cleavage. After a ribosome unfolds the structure surrounding the siRNA site, the site is available for binding to AGO2/siRNA complexes until structures reform. Therefore, the more frequently an mRNA is translated by ribosomes, the more time the AGO2 target site will be in an unmasked state, which will result in a higher cleavage rate of the mRNA. However, the time that the AGO2 target site is in an unmasked state also depends on the mRNA refolding rate. In case of a very slow refolding rate, an siRNA target site that is unfolded by a ribosome, will not refold to its masked conformation before the next ribosome arrives at the siRNA target site. In this case, a higher translation rate will not increase the time the AGO2 target site will be in an unmasked state and thus does not lead to an increase in the cleavage rate. Therefore, the cleavage rate is dependent on both the translation rate (how often do ribosomes unfold the AGO2 target site) as well as the mRNA refolding rate (how fast do structures reform after a ribosome unfolds the AGO2 target site). To determine the refolding rate of our reporter mRNA, we therefore analyzed the difference in cleavage rates of two reporters with identical coding sequences and siRNA binding sites, but with different translation rates: the KIF18B reporter and the uORF-KIF18B reporter (which has a 3.3 fold lower translation rate, see section *‘Number of ribosomes per mRNA molecule’*). As we know the difference in translation rate between the two reporters, we can determine the expected difference in cleavage rate at different mRNA refolding rates and compare these values to the experimentally-observed difference in cleavage rate of the KIF18B and uORF-KIF18B reporters. To identify the mRNA refolding rate that matches the difference between the experimentally-derived cleavage rate of the KIF18B and uORF-KIF18B reporters best, we performed stochastic simulations to simulate cleavage curves of the KIF18B and uORF-KIF18B reporter at a range of different mRNA refolding rates. Next, we assessed at which mRNA refolding rate the difference between the simulated KIF18B and uORF-KIF18B cleavage curves matched the difference between the experimental cleavage curves by determining an SSE for the experimental cleavage curve with each simulated cleavage curve.

We simulated the cleavage curves for both the KIF18B reporter and the uORF-KIF18B reporter at different siRNA concentrations (10 nM, 0.75 nM, and 0.1 nM). As the cleavage curve is based on the cleavage time of individual mRNAs, we simulated the cleavage time of a single mRNA, and repeated this for many mRNAs to obtain a simulated cleavage curve. To simulate the cleavage time of a single mRNA, we assumed that each mRNA could be cleaved either through; (1) a ribosome-dependent cleavage pathway: ribosome-facilitated AGO2 target binding through target site unmasking by ribosomes translating the siRNA target site; (2) a ribosome-independent cleavage pathway (this was included since mRNAs are also cleaved in the absence of ribosomes translating the siRNA target site). For each mRNA we therefore performed two parallel simulations: we simulated a cleavage time through the ribosome-independent cleavage pathway and through the ribosome-dependent cleavage pathway. The lowest of the two values was used as the cleavage time for that mRNA. To determine the cleavage time through the ribosome-independent pathway, a random number was selected from an exponential distribution that was generated by fitting the cleavage curve obtained in the presence of CHX.

To determine the cleavage time through the ribosome-dependent pathway, we simulated (1) when ribosomes arrive at the siRNA target site, and (2) for each ribosome if cleavage would occur during the time that the target site was unmasked by that ribosome. To simulate when ribosomes arrive at the siRNA target site we distinguished between the first and subsequent ribosomes. The arrival time of the first ribosome at the siRNA target site was simulated by selecting a random number from a gamma distribution, where the shape and scale parameter were taken from the first ribosome arrival distribution at the siRNA target site that we determined previously (see section *‘arrival time of the first ribosome at the siRNA binding site’*). In order to determine the arrival time of the next ribosome at the siRNA target site, we added together two values; (1) a fixed, minimum inter-ribosome arrival time, which was introduced, because each ribosome occupies multiple nucleotides on an mRNA and therefore a minimal nucleotide distance is always present between the centers of two ribosomes; (2) the time between two subsequent translation initiation events. In the simulations we assumed that the minimal distance between two ribosomes is 30 nucleotides (about the size of one ribosome footprint), and we converted the minimal inter-ribosome distance to a minimal inter-ribosome arrival time by dividing the minimal inter-ribosome distance with the average elongation speed of ribosomes on the KIF18B reporter (as was determined from the first ribosome arrival distribution). To simulate the time between two initiation events, we drew a number from an exponential distribution, which was based on the experimentally observed initiation rates of both the KIF18B and uORF-KIF18B reporters.

Next, we determined for each ribosome arriving at the siRNA target site if cleavage would occur during the time that the target site was unmasked by that ribosome; we assume that each ribosome unfolds the siRNA target site when the ribosome passes the site, which can lead to one of three possible outcomes: (1) the mRNA refolds (and will be opened again when the next ribosome arrives); (2) AGO2 binds and cleaves the mRNA; or (3) the mRNA remains open until the next ribosome appears at the site without getting cleaved.

We simulated the time for mRNA refolding and the time for AGO2-mediated mRNA cleavage (both of which are unknown) by drawing two random numbers. (1) The mRNA refolding time was drawn from an exponential distribution. The simulation is run multiple times with different values for the mean of the mRNA refolding time distribution (37 times with a range of 0.1 s to 1000 s). (2) The mRNA cleavage time was drawn from an exponential distribution. The mean of this distribution is the average mRNA cleavage time (which includes the time for AGO2 target binding, mRNA slicing, and fragment release). Simulations were performed using multiple distributions with distinct mean values and the simulation that resulted in the best fit with the data was selected and the mean of the distribution was recorded. The best fit was selected for each simulated mRNA refolding time. (3) The time until the next ribosome arrives at the siRNA binding site is known as we already determined the arrival time of all the ribosomes at the siRNA target site.

To simulate the outcome of a ribosome-dependent mRNA unfolding event (i.e. next ribosome arrives, mRNA refolds, or AGO2 binds and cleaves the mRNA), we selected the simulated process which was the fastest. Finally, the cleavage time through the ribosome-dependent pathway was determined by taking the arrival time of the ribosome after which AGO2-mediated mRNA cleavage occurred, and adding the AGO2 cleavage time. Using the simulations described above, we can simulate cleavage curves for the KIF18B and uORF-KIF18B reporters at different mRNA refolding rates. The values of the parameters used in the simulations can be found in the overview below.

| Parameter | Value |
| --- | --- |
| Cleavage rate (ribosome-independent pathway) | 10 nM control: $6.6396 \cdot 10^{-4} \text{ s}^{-1}$, 10 nM uORF: $6.1039 \cdot 10^{-4} \text{ s}^{-1}$, 0.75 nM control: $7.4634 \cdot 10^{-4} \text{ s}^{-1}$, |
| | 0.75 nM uORF: $6.9877 \cdot 10^{-4} \text{ s}^{-1}$, 0.1 nM control: $3.8190 \cdot 10^{-4} \text{ s}^{-1}$, and 0.1 nM uORF: $3.3234 \cdot 10^{-4} \text{ s}^{-1}$ |
| First ribosome arrival time distribution at the siRNA target site (gamma distribution) | Shape: 15.34 and scale: 18.42. |
| Inter-ribosome distance | 30 nt |
| Ribosome elongation speed | 11.13 nt/second |
| Ribosome initiation rate | Control: $0.0469 \text{ s}^{-1}$ and uORF: $0.0135 \text{ s}^{-1}$ |

During the simulations we assumed that the mRNA refolding rate and the AGO2 cleavage rate are similar between the KIF18B and uORF-KIF18B reporter as both reporters have an identical coding sequence and siRNA target sequence. Next, we wanted to determine for each siRNA concentration (10 nM, 0.75 nM, and 0.1 nM) and each simulated mRNA refolding rate (*k_refold_*), the AGO2 cleavage rate (*k_on_*) that best fit the experimentally-observed difference between the cleavage curves of the KIF18B and uORF-KIF18B reporter. To find the AGO2 cleavage rate that best matched the data, we calculated a total sum of squared errors (*total SSE*) at each simulated mRNA refolding rate, which reports on how well the simulated cleavage curves matched the experimental cleavage curves for both the KIF18B and uORF-KIF18B reporters (see Equation 6).

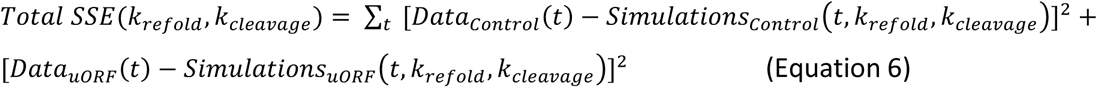

The total sum of squared errors (*total SSE*) is calculated by adding together the sum of squared errors of the KIF18B reporter and the sum of squared errors of the uORF-KIF18B reporter. The sum of squared errors of each reporter represents the difference between the experimentally-derived cleavage curve and the simulated cleavage curve at a specific mRNA refolding rate and AGO2 cleavage rate. Thus, the AGO2 cleavage rate that best fit with the experimental data at each simulated mRNA refolding rate can be found by minimizing the *total SSE*. To minimize the *total SSE* at each mRNA refolding rate, we simulated cleavage curves for a range of mRNA refolding rates (between 0.001 and 10 s^−1^), and at each mRNA refolding rate we searched for the AGO2 cleavage rate that minimized Equation 6. Each individual simulation consisted of 10.000 mRNAs, was performed in 10 replicates using a different random seed in each replicate, and all replicates were averaged. In this way, we obtained a *total SSE* value for each mRNA refolding rate.

To compare the *total SSE* values at different mRNA refolding rates, we first normalized the *total SSE* value at each mRNA refolding rate with the lowest possible SSE value obtained when the experimental KIF18B and uORF-KIF18B cleavage curves were fit independently, at each mRNA refolding rate. To calculate this lowest possible SSE value we fit the KIF18B and uORF-KIF18B cleavage curves independently and calculated an *independent SSE* for both the KIF18B and uORF-KIF18B reporters using Equation 7. The lowest possible SSE value is the sum of both *independent SSE*s.

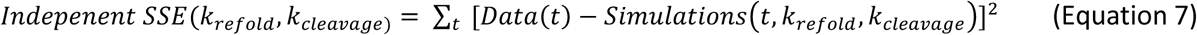

The AGO2 cleavage rate that best fit at each mRNA refolding rate with the experimental data for either the KIF18B or uORF-KIF18B cleavage curve can be found by minimizing the *independent SSE*. To minimize the *independent SSE*, we simulated cleavage curves for a range of mRNA refolding rates (between 0.001 and 10 s^−1^), and at each mRNA refolding rate we searched for the AGO2 cleavage rate that minimized Equation 7. After finding the *independent SSE* for both the KIF18B and uORF-KIF18B reporter at each mRNA refolding rate, we can normalize the *total SSE* using Equation 8.

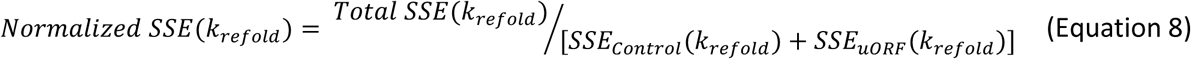

For each simulated mRNA refolding rate, we can normalize the *total SSE* by the sum of the KIF18B and uORF-KIF18B *independent SSE*s that were determined with Equation 7. As the sum of the KIF18B and uORF-KIF18B *independent SSE* represents the minimal *total SSE* possible, we obtain a *normalized SSE*.

#### Simulations of RNA structural dynamics

Our results showed that in the absence of ribosomes translating the siRNA target site, the cleavage rate is very similar at different siRNA concentrations. In contrast, lowering the siRNA concentration in the presence of ribosomes translating the siRNA target site strongly decreased the cleavage rate. These findings together suggest that in the presence of ribosomes translating the siRNA target site the AGO2/siRNA complex concentration is rate-limiting, while in the absence of ribosomes translating the siRNA target site, the availability of the siRNA target site can become another rate-limiting step (i.e. the target site is often in a masked state, unavailable for AGO2/siRNA complex binding). We can use the sensitivity of the cleavage rate to different siRNA concentrations in the absence of ribosomes translating the siRNA target site to probe the structural dynamics of the reporter mRNA; in case mRNAs change between different structural conformations on fast time scales (i.e. fast structural dynamics), it is expected that target site masking and unmasking also occur fast, as the siRNA target site is expected to be masked in only a subset of structural conformations. In this scenario, target site unmasking is not a rate-limiting step and the cleavage rate is expected to be solely determined and thus very sensitive to the siRNA concentration. In contrast, when mRNAs change between different structural conformations on slow time scales (i.e. similar or slower time scales than the AGO2-target binding), target site unmasking becomes rate-limiting as well. In this case the cleavage rate is expected to be less sensitive to the siRNA concentration as the cleavage rate is only partially determined by the siRNA concentration. Thus, to investigate the structural dynamics of our reporter mRNA (i.e. how fast does the siRNA target site change between a masked and unmasked state), we decided to simulate cleavage curves at different siRNA concentrations, and with different unmasking and masking rates. Specifically, we wanted to determine at which unmasking and masking rate, the effect of lowering siRNA concentration on the simulated cleavage curves best matched the experimentally-observed effect of lowering the siRNA concentration.

To determine the effect of lowering the siRNA concentration on the cleavage curve in the absence of ribosomes translating the siRNA target site, we simulated cleavage curves at different siRNA concentrations, and different unmasking and masking rates. To obtain a simulated cleavage curve, we simulated the cleavage time of a single mRNA and repeated this many times. To simulate the cleavage time of a single mRNA, we set-up a simulation where an mRNA can be in three different states; in the first two states the mRNA is intact and either (1) in a *masked* state, unavailable for AGO2/siRNA complex binding and mRNA cleavage, or (2) in an *unmasked* state, available for AGO2/siRNA complex binding and mRNA cleavage. In the third state the mRNA is in (3) a *cleaved* state. As an mRNA is only available for AGO2/siRNA complex binding in the *unmasked* state, transitions can only occur from the *unmasked* to *cleaved* state, and transitions can occur between the *unmasked* and *masked* states. In the simulations, each mRNA starts either in the *masked* or *unmasked* state and transitions between the different states are simulated until the mRNA reaches the *cleaved* state. The time it takes in the simulations to reach the *cleaved* state is the cleavage time for that mRNA. To simulate the transitions and the transition times (how long does it take to transition from one state to the other), random numbers are selected from distributions, as described below.

#### Masked to unmasked transition

The transition time from the *masked* to *unmasked* state is referred to as the unmasking time. The mean unmasking time is unknown and the simulation is run multiple times with different values for the mean unmasking time (16 times with a range of 1 s to 1500 s for the mean unmasking time). The unmasking time is simulated by selecting a random number from an exponential distribution where the mean is the mean unmasking time.

#### Transition from the unmasked to either the masked or cleaved state

When an mRNA is in the *unmasked* state, it can transition either to the *masked* state or to the *cleaved* state. The transition time from the *unmasked* to the *masked* state is referred to as the masking time and the transition time from the *unmasked* to *cleaved* state is referred to as the cleavage time. To determine if an mRNA transitions to the *masked* or *cleaved* state, we simulate in parallel two transition times. (1) A masking time was simulated. The mean masking time is unknown and simulations were performed using multiple exponential distributions with distinct mean values. The simulation that resulted in the best fit with the data was selected, and the mean of the distribution was recorded. The masking time is simulated by selecting a random number from an exponential distribution where the mean is the mean masking time. (2) The AGO2 cleavage time, which is simulated by selecting a random number from an exponential distribution where the mean is the experimentally-derived average AGO2 cleavage time (which includes AGO2 target binding, mRNA slicing, and fragment release). The average AGO2 cleavage time (i.e. the AGO2 cleavage cycle duration) was already determined before at each siRNA concentration (10 nM, 0.75 nM, and 0.1 nM) (see Figure 3B) and was used in the simulations. The shortest of the two simulated transition times determines if an mRNA transitions to the *unmasked* or *cleaved* state.

Finally, we simulate the starting state of each single mRNA, which can be in the *masked* or *unmasked* state. To simulate the starting state, we assume that at the start of our experiment the ratio between the *masked* and *unmasked* state has reached an equilibrium that depends on the average masking, unmasking and cleavage time. For example, if the average masking time is very long while the average unmasking time is very short, it is expected that most mRNAs are in the *unmasked* state. To obtain the ratio between the *masked* and *unmasked* state we run an initial simulation before starting the full simulation, where all mRNAs start in the *masked* state. Next, we evaluate the initial simulation until the ratio of mRNAs in the *masked* and *unmasked* state reaches an equilibrium (which will depend on the masking, unmasking and cleavage times used in this simulation). This ratio is used in the full simulation to determine if an mRNA starts in the *masked* or *unmasked* state.

Thus, we simulate for single mRNAs in which state they start the simulation and how long it takes to transition to the *cleaved* state. By repeating this procedure for many mRNAs, we obtain a simulated cleavage curve. Next, we performed two simulations where we assessed the effect of lowering siRNA concentration, from 10 nM to either 0.75 nM or 0.1 nM, on the cleavage curves in the absence of translating ribosomes. In both simulations we simulated the effect of lowering the siRNA concentration at a range of mean unmasking and masking rates, and compared the simulated cleavage curves to experimentally-derived cleavage curves. For each mean unmasking time (we took a range of 1 s to 1500 s), we searched for the average masking time that resulted in the best fit with the data by calculating a goodness-of-fit score (*total SSE*) using Equation 9.

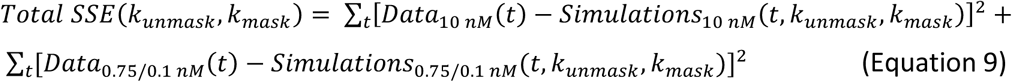

The *total SSE* is calculated by adding together the SSE for the 10 nM cleavage curve in the absence of translating ribosomes (we used the experiments where we added CHX) and the SSE of either the cleavage curve with 0.75 nM siRNA, or 0.1 nM siRNA concentration in the absence of translating ribosomes (we used the experiments where we added CHX). The *total* SSE represents the difference between the experimentally-derived cleavage curve and the simulated cleavage curve at a specific unmasking and masking rate. For a range of mean unmasking times (1 s to 1500 s), we simulated cleavage curves with different mean masking times and selected the masking time that minimized the *total SSE* (Equation 9). Each individual simulation consisted of 10.000 mRNAs, and was done in 5 replicates using a different random seed in each replicate. The average of the 5 replicates was reported. In this way, we obtained a *total SSE* value for each unmasking rate.

